# Design-of-Experiments for Nonlinear, Multivariate Biology: Rethinking Experimental Design through Perturb-seq

**DOI:** 10.64898/2025.12.28.696309

**Authors:** Yuji Okano, Tetsuo Ishikawa, Yasunori Sato, Hideyuki Okano, Kazuhiro Sakurada

## Abstract

Design of experiments (DOE) principles are increasingly applied to biological assays, yet it remains unclear whether the optimality of their foundational assumption—orthogonal decomposition—holds in nonlinear biological systems.

We addressed this question using Perturb-seq as a case study. By benchmarking a design commonly used in Perturb-seq and related experiments against the orthogonal Plackett–Burman (PB) design via simulations, we uncovered a counter-intuitive phenomenon: while orthogonal designs generally excel, the correlated structure inherent to the common design is functionally robust in systems with significant signal amplification. This challenges the blind application of DOE to biology.

Based on these findings, we developed the PB suitability index (PBSI), a simple, parameter-free metric that predicts the optimal design solely from network structure. Our work not only provides practical guidelines for Perturb-seq but also establishes a “biology-oriented DOE” framework, bridging the gap between statistical rigor and biological complexity.

## 1 Introduction

Biological experimentation involves extracting *events* (one-off occurrences) from *processes* (continuous sequences of changes driven by scale interactions) and interpreting these events through the causal inference schemes: correlation, intervention, and counterfactuals [1]. Recent progress in high-throughput quantitative measurements has dramatically expanded the scale of this causal inquiry, necessitating multivariate approaches.

In many multivariate biological experiments, a design that tests one factor at a time against a baseline—which we previously termed the “Control-and-Leave-One-Out (C+LOO) design” [2]—has long served as the standard approach for verifying causal effects. Despite its practical convenience, the C+LOO strategy and its variants are traditionally criticized for their limited capacity to detect interactions [3]. Therefore, the integration of multivariate biological assays and DOE—a statistical framework designed to balance feasibility and resolution under experimental constraints [4]—is gaining increasing attention [5].

However, while DOE provides a principled mathematical foundation—notably orthogonality and decomposability—it remains uncertain to what extent these properties remain optimal in nonlinear, open, and nonequilibrium biological systems [6]. In other words, although DOE is increasingly invoked in modern biology, whether its classical principles can be seamlessly transplanted to nonlinear biological contexts without theoretical reassessment remains an open question. Given that biological experiments often extend to realms where discrete events and continuous processes conceptually fuse together, an epistemological classification of biological contexts—clarifying what can be effectively demonstrated by causal modeling—is essential before applying statistical tools.

To address this gap, we establish two complementary lines of inquiry. First, we examine whether experimental designs used in multivariate biological assays can be systematically extended and analyzed using DOE principles. Second, we evaluate how far the foundational assumption of orthogonal decomposability, central to the DOE discipline, remains meaningful when the generative process is nonlinear.

As an initial contribution toward a biology-oriented DOE framework, we analyze (i) statistical properties of the C+LOO design, (ii) their impacts on the result interpretation, and (iii) their benchmarking against the PB design [4], focusing on Perturb-seq—a CRISPR-based single-cell perturbation assay [7, 8]—as a prototypical case where the C+LOO design is widely employed [2]. We also provide an experimental design guideline for Perturb-seq to support understanding and comparison of experimental designs.

### 2 Results

### 2.1 Inherent correlated structure of the C+LOO design is sensitive to the number of factors *n*

Having one control trial and *n* knockdown (KD) trials, the C+LOO design for *n* factors (∀*n* ∈ ℕ)—which we denote *C*(*n*)—consists of *n* + 1 trials (see Methods 4.1.1 for details). As an example, *C*(9) is shown in Figure 1a.

**Figure 1.**
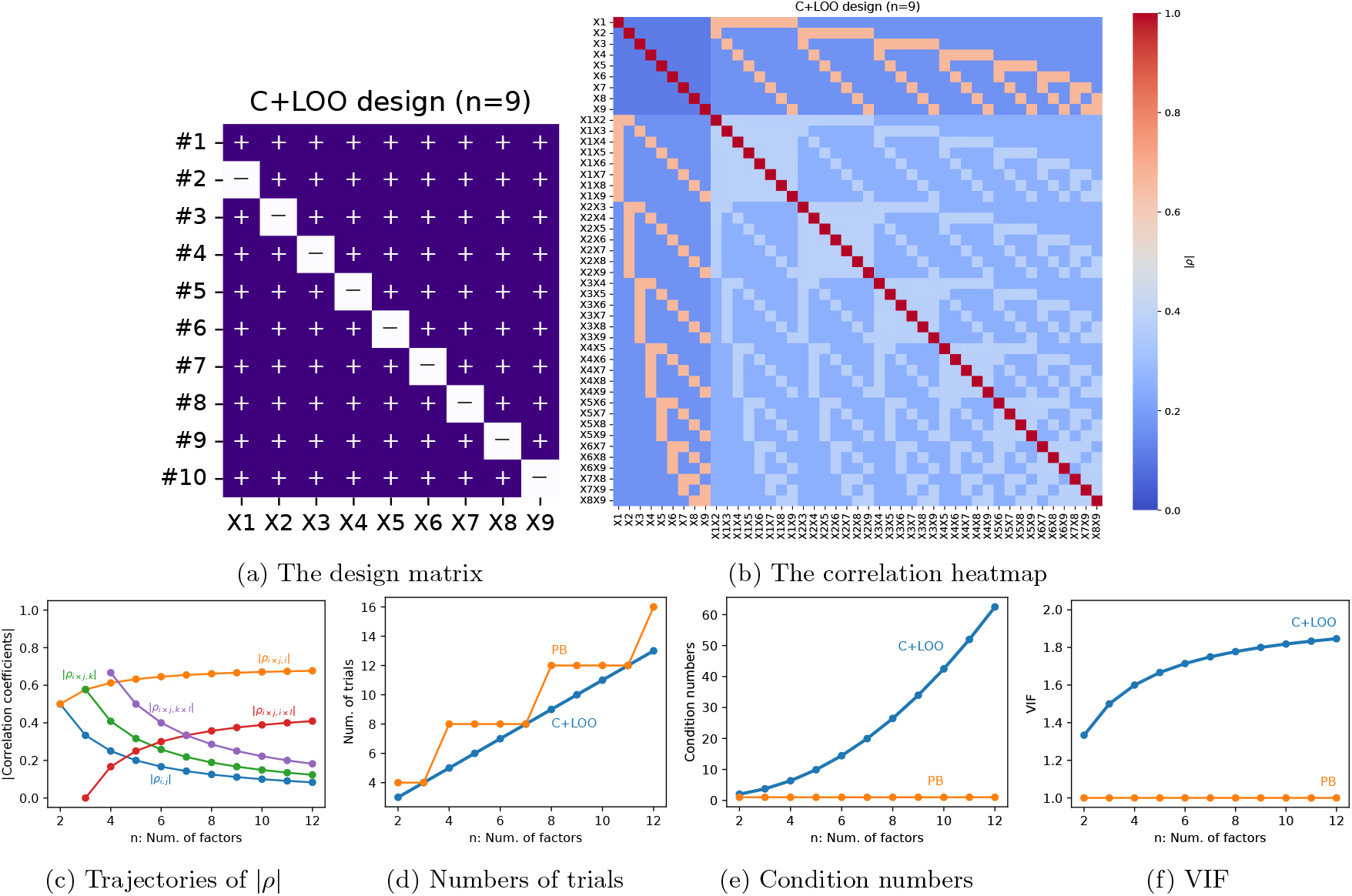
The statistical properties of the C+LOO design depend on the number of factors *n*. **a**: Design matrix of the C+LOO design for nine factors (+, addition of factors;, − subtraction of factors). The rows (#1–#10) represent trials and the columns (X1–X9) represent the factors. **b**: Heatmap of the absolute correlation coefficients (denoted as |*ρ*|) among main effects (X1–X9) and interaction terms (e.g., X1X2· · ·) when they are investigated under the design matrix. **c**: |*ρ*| for C+LOO designs of 2–12 factors (*ρ*_*i,j*_ between two different main effects **x**_*i*_ and **x**_*j*_, blue; *ρ*_*i×j,i*_ between an interaction term **x**_*i×j*_ and one of its corresponding main effects **x**_*i*_, orange; *ρ*_*i×j,k*_ between an interaction **x**_*i×j*_ and a non-corresponding main effect **x**_*k*_, green; *ρ*_*i×j,i×l*_ between two interaction sharing one main effect such as **x**_*i×j*_ and **x**_*i×l*_, red; and *ρ*_*i×j,k×l*_ between two interaction terms with no shared main effect such as **x**_*i×j*_ and **x**_*k×l*_, purple) for ∀*n* ∈ Euclid Math Two, indices *i, j, k, l* ∈ Euclid Math Two such that 1 ≤ *i, j, k, l* ≤ *n* and all indices are distinct. Notably, here we omitted *ρ*_*i,i*_ and *ρ*_*i×j,i×j*_ since they are constant. **d–f** : The numbers of trials (d), condition numbers (e), and VIF (f) for the C+LOO (blue) and PB (orange) designs of 2–25 factors. These results demonstrate that the statistical structure of the C+LOO design is not fixed but evolves with *n*, undermining naive design benchmarking across studies with different target sizes.

By visualizing the absolute values of the Pearson correlation coefficients *ρ* among the main effects and interaction terms of *C*(9) (Figure 1b), we confirmed a distinctive property of the C+LOO design: generally weak correlations among main effects and interaction terms [2].

To examine the behavior for general *n*, we derived closed-form expressions for the correlation coefficients of generic main effects **x**_*i*_ and interaction terms **x**_*i×j*_ (Methods 4.1.4–4.1.5 and Supplementary Information (SI) A– B). As illustrated in Figure 1c for 2 ≤ *n* ≤ 12, some absolute correlations decrease with *n* whereas others increase. While this “trade-off” structure maintains a nonvanishing correlation profile for all *n*, the detailed composition differs according to the value of *n*. In other words, both the magnitude and the pattern of factor correlations in the C+LOO design vary with the number of factors *n*.

Notably, *ρ*_*i×j,i*_ and *ρ*_*i×j,i×l*_ converge to positive values as *n* increases, suggesting that C+LOO becomes increasingly susceptible to confounding from interaction terms as the factor count grows. In sharp contrast, the well-known full factorial (FF) and PB designs ensure *ρ*_*i×j,i*_ = 0 (Methods 4.1.2–4.1.3 and SI C), underscoring this as a characteristic feature of C+LOO.

We further found that the *n*-dependent evolution of these correlations exacerbates multicollinearity. For comparison, we instantiated the PB design—requiring a similar total number of runs as C+LOO for each *n* (Figure 1d and Methods 4.1.3)—and evaluated the condition number and variance inflation factor (VIF). Whereas PB remained constant across *n*, the C+LOO design showed clear increases with *n* (Figures 1e–1f).

In summary, the C+LOO design is characterized by *n*-dependent shifts in the trade-off structure of pronounced factor correlations. The latter presents a significant challenge for comparative studies of experimental designs. Although benchmarking with real single-cell RNA-sequencing (scRNA-seq) datasets is standard practice in comparing scRNA-seq methodologies, the statistical properties of C+LOO change with the number of target genes, making broadly generalizable conclusions difficult. Thus, rather than relying solely on inductive comparisons from individual datasets, one should seek more abstract, design-level principles—grounded in theory and universality—to understand how experimental design influences inference.

### 2.2 Simulated models revealed strengths and weaknesses of the C+LOO design in Perturb-seq data analysis deriving from the inherent correlations

Given the unique factor correlation patterns inherent in the C+LOO design, further investigation is essential to understand how these features influence the conclusions drawn from Perturb-seq experiments.

To evaluate the influence of experimental design on the interpretation of Perturb-seq results, it is essential to formally define its underlying logical framework and the biological phenomena it addresses. We established the following formalization to delineate the boundaries of the research domain where experimental design can intervene:

#### 1. Quantification Framework

Conventional Perturb-seq (using the C+LOO design) quantifies phenotypes resulting from individual gene KD by summarizing them into scalar values (e.g., singlegene markers or phenotypic scores) and analyzing the difference relative to a baseline condition (Figure 2).

**Figure 2.**
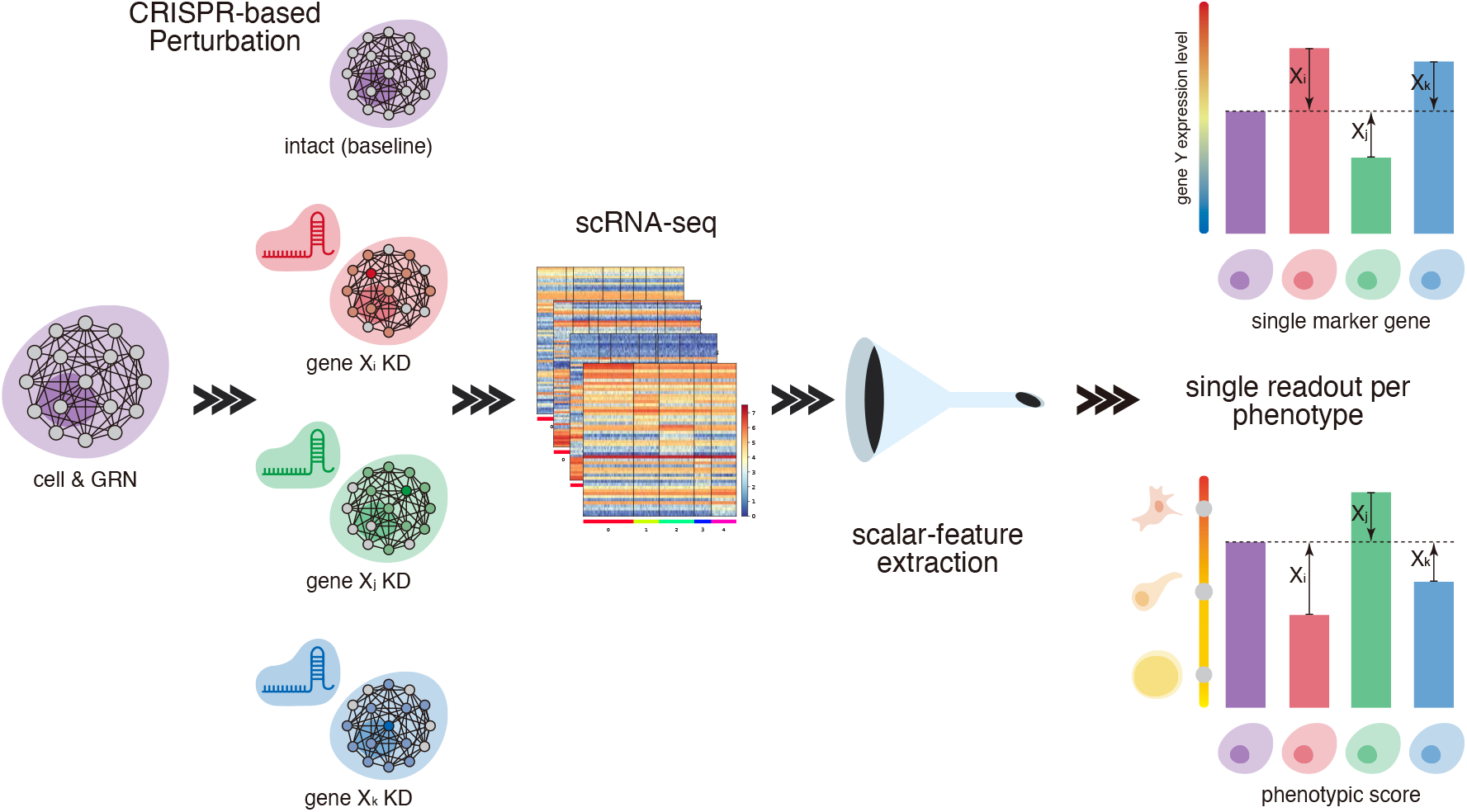
Standard procedure for Perturb-seq experiments and data analysis. Perturb-seq is a combinatorial approach that couples CRISPR-based screening with scRNA-seq [7, 8]. Using a C+LOO design, one-gene-at-a-time KD models are constructed, and changes in gene-expression patterns are profiled by scRNA-seq. The intact condition (purple cell) serves as the baseline, and each KD model (pink, green, blue cells) is evaluated in a pairwise comparison (e.g., Dunnett’s test [9]) against this baseline. To enhance interpretability for a phenotype, the readout is often (i) single-gene markers—commonly used proxies for specific phenotypes, including lineage markers (e.g., *HES1*), master regulators (e.g., *NANOG, FOXP3*), and oncogenes (e.g., *MYC, KRAS*)—or (ii) phenotypic scores that aggregate multiple marker genes (e.g., via SCENIC [10]). These readouts share a common property: they are scalar. As an example, the right panels illustrate pairwise comparisons to the baseline: the top-right uses the expression level of gene Y as the readout, and the bottom-right uses a phenotypic score. In both cases, comparisons are shown for genes X_i_ (pink), X_j_ (green), and X_k_ (blue). Thus, a standard Perturb-seq workflow can be abstracted as follows: from scRNA-seq data acquired under a C+LOO design, extract a phenotype-aligned scalar feature and assess the KD-induced difference by pairwise comparison to the baseline. In this study, to examine how the experimental design and scalar readout shape downstream biological inference, we explicitly consider the underlying GRN as a mediating variable. For further details, see Methods 4.2 and SI D.1.

#### 2. Nature of the Causal Model

This methodology relies on a discrete causal model between the perturbation and the phenotype, treating the detailed, intermediate mechanism as a black box (Methods 4.2 and SI D.1).

#### 3. Limitation of Applicability

Due to the observational “blank time” inherent in Perturb-seq, the model’s application is restricted to systems where the gene regulatory network (GRN) structure is static (time-invariant). Discussions regarding the efficiency of experimental design are therefore only valid under this constraint (Methods 4.2 and SI D.1).

Based on this formalization, we devised a methodology to systematically evaluate the impact of experimental design. Since empirical analysis using real data presents limitations for constructing systematic knowledge (SI D.2), we developed a minimalist simulator that extracts only the essential elements of Perturb-seq (Methods 4.3): (i) a static GRN, (ii) nonlinear interactions based on a specified true GRN structure, and (iii) scalar output. Through this generalized model, we aim to derive universal principles governing the relationship between experimental design and result interpretation, ultimately providing guiding principles for decision-making in future Perturb-seq experiments.

Based on this framework, we investigated the impact of the inherent correlations within the C+LOO design. We first established three interpretable scenarios (Methods 4.3.1–4.3.3) and constructed corresponding simulators—Models Φ, Ψ, and Λ—for each (Figures 3a– 3c). We confirmed that the simulators reproduce Taylor’s power law (TPL), a ubiquitous property of scRNA-seq data [11–15], ensuring biological plausibility for the subsequent benchmarks (Methods 4.3.5 and SI D.3).

**Figure 3.**
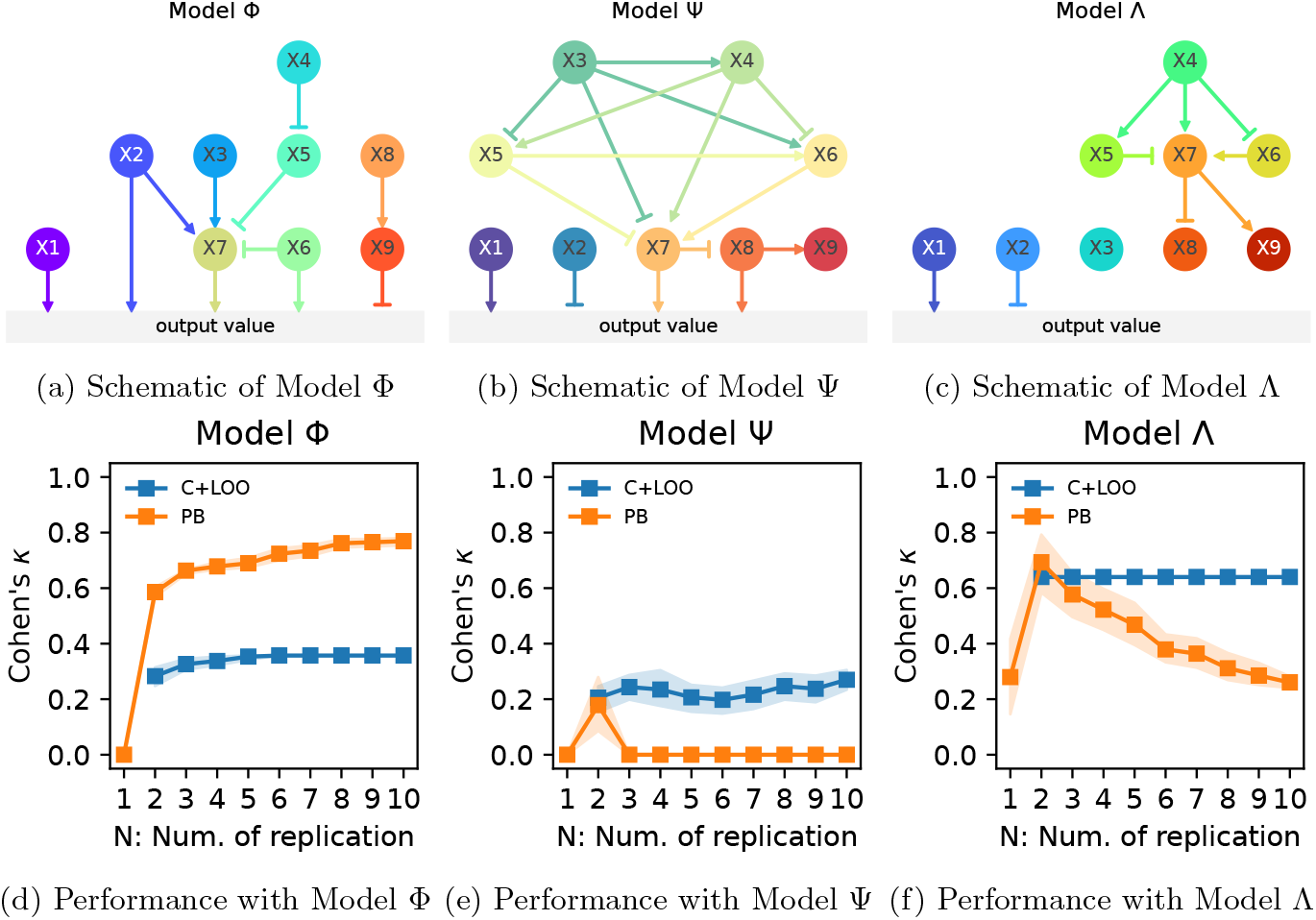
Simulated experiments highlighted the pros and cons of the C+LOO design. **a–c**: Schematics of the minimalist simulators. Constructed upon a GRN framework, these models feature nine interacting genes (X1–X9) regulating a non-negative scalar output via positive (pointed arrows) or negative (T-shaped arrows) edges, consistent with conventions in Boolean networks—a popular class of biological network modeling [17]. Model Φ (a) incorporates nine factors modulating the output via multiple direct and indirect pathways (Methods 4.3.1). Model Ψ (b) emulates a complex cascade featuring both synergistic and antagonistic interactions (Methods 4.3.2). Model Λ (c) represents a sparse factor scenario (poor candidate filtering) where the majority of genes exert no influence (Methods 4.3.3). All simulators share a unified architecture: (i) directed acyclic graph (DAG)-based GRN structures; (ii) parameters sampled from log-normal distributions; (iii) scalar outputs mimicking Perturb-seq readouts (e.g., single-gene markers or phenotypic scores); and (iv) rectified linear unit (ReLU) activation to introduce biological nonlinearity (Methods 4.3). Biological relevance was ensured by validating adherence to TPL—a ubiquitous power-law scaling between means and variances observed in scRNA-seq data [11–15] (SI D.3). **d–f** : Performance comparison of the C+LOO (blue) and PB (orange) designs based on the *κ* metric across replication levels *N* = 1–10. Markers represent the mean over 30 random seeds, with shaded areas indicating 95% bootstrap CIs. Data for the C+LOO design at *N* = 1 are omitted as ANOVA requires replication (SI E). For comparisons with Dunnett’s test (a representative pairwise method showing lower noise robustness), refer to Figures S8–S10 and SI G.4.

Using these simulators, we evaluated how the C+LOO design’s correlated structure affects the interpretation of experimental results through the following evaluation pipeline:

#### 1. Definition of ground truth main effects

To quantify the accuracy of analytical results from C+LOO and PB designs, ground truth main effects were defined for each gene, serving as an anchor for evaluating performance (Methods 4.4.6).

#### 2. Simulation of experiments

Simulations based on the C+LOO and comparative PB designs were conducted to generate scalar output values for each condition.

#### 3. Analysis of experimental results

Given that conventional pairwise comparisons suffer from context-dependency due to reliance on specific baselines (detailed in SI E), we adopted standard DOE multivariate analyses—specifically multivariate linear regression (MLR) and analysis of variance (ANOVA)—to evaluate the directionality and statistical significance of the estimated effects.

#### 4. Benchmarking experimental design performance

The concordance between the regulatory states inferred by the C+LOO and PB designs and the ground truth labels was quantified using weighted Cohen’s *κ* coefficients (Methods 4.5.1), with uncertainty estimated via 95% bootstrap confidence intervals (CIs); comparative benchmarks against pairwise testing are detailed in SI G.4.

The relative performance of the designs varied across simulators (Figures 3d–3f): the PB design generally outperformed in Model Φ, whereas the C+LOO design performed better in Models Ψ and Λ. We also observed consistent trends across different Gaussian noise levels (SI G), with one exception in Model Λ: The C+LOO performed better under low noise conditions, whereas the PB design showed a clear advantage under high noise.

Importantly, inherent correlation patterns play a central role in determining design performance. The characteristic differences between the two designs—and their performance trends across simulators—persisted even after applying D-optimization, a common strategy for improving design matrices by reducing factor correlations [16] (SI H):

- D-optimized C+LOO (DO-C+LOO) designs exhibited more intensive correlation patterns than PB and D-optimized PB (DO-PB) designs
- D-optimization improved the performance of the C+LOO in scenarios favoring PB designs, while degrading it in cases where C+LOO outperformed

These simulated results collectively show that the performance of experimental designs is closely linked to the network structures of the simulated models. In systems regulated by parallel cascades—as modeled in Model Φ—the C+LOO design may underperform compared to the PB design, suffering from undesirable “contribution leakage” (shown in SI F) due to confoundings. However, situations where the target phenomenon is predominantly governed by a cascade involving complex interactions—such as Model Ψ—can present counterintuitive instances: the “leakage” of interactions via *ρ*_*i×j,i*_ coincidentally yields a more robust estimation of main effect signs compared to the PB design with the opposing strategy of orthogonal decoupling.

Results from Model Λ can also be explained with the network structure and factor correlations. Model Λ reflects a scenario with poor candidate gene selection where most factors do not influence the target outcome. Under such circumstances, ensuring sign consistency can be difficult due to the numerical instability of the model (SI G.3).

Since both multivariate analyses and pairwise testing operate within the general linear model framework, they fundamentally rely on inverting the model matrix (Methods 4.4.3–4.4.4). Consequently, design-inherent factor correlations are unavoidable in the C+LOO design, even when factors are independent. Depending on the network structure among factors, such inherent correlation structures can yield robustness, impair inference, or show no consistent effect on design performance.

### 2.3 Deriving a topological indicator for experimental design selection

Thus far, we have analyzed the C+LOO design from a DOE perspective, elucidating its correlation structure. Our simulations of three biologically plausible scenarios revealed that while inherent correlations can lead to erroneous conclusions, they can paradoxically yield robustness in specific contexts—a phenomenon counterintuitive to traditional DOE principles. This underscores a critical reality: there is no single “omnipotent” experimental design in biology; rather, the optimal strategy is strictly context-dependent.

How, then, should researchers select an appropriate design? Bridging this gap between theory and practice requires a framework that translates qualitative domain knowledge into quantitative decision-making. To achieve this, we adopted a data-driven approach: we first employed machine learning (ML) to identify key structural features governing design preference—C+LOO or PB— using exhaustive simulations, and subsequently distilled these findings into a simple, scalable indicator named the PBSI.

We began by defining a set of terms to quantitatively describe GRN structures, including *regulations, pathways, cascades, regulation weight, factor weight*, and *cascade length* (see Figure 4 and Methods 4.7.1 for definitions). Based on these concepts, we engineered 31 features designed to be scalable regardless of the number of factors (Methods 4.7.2).

**Figure 4.**
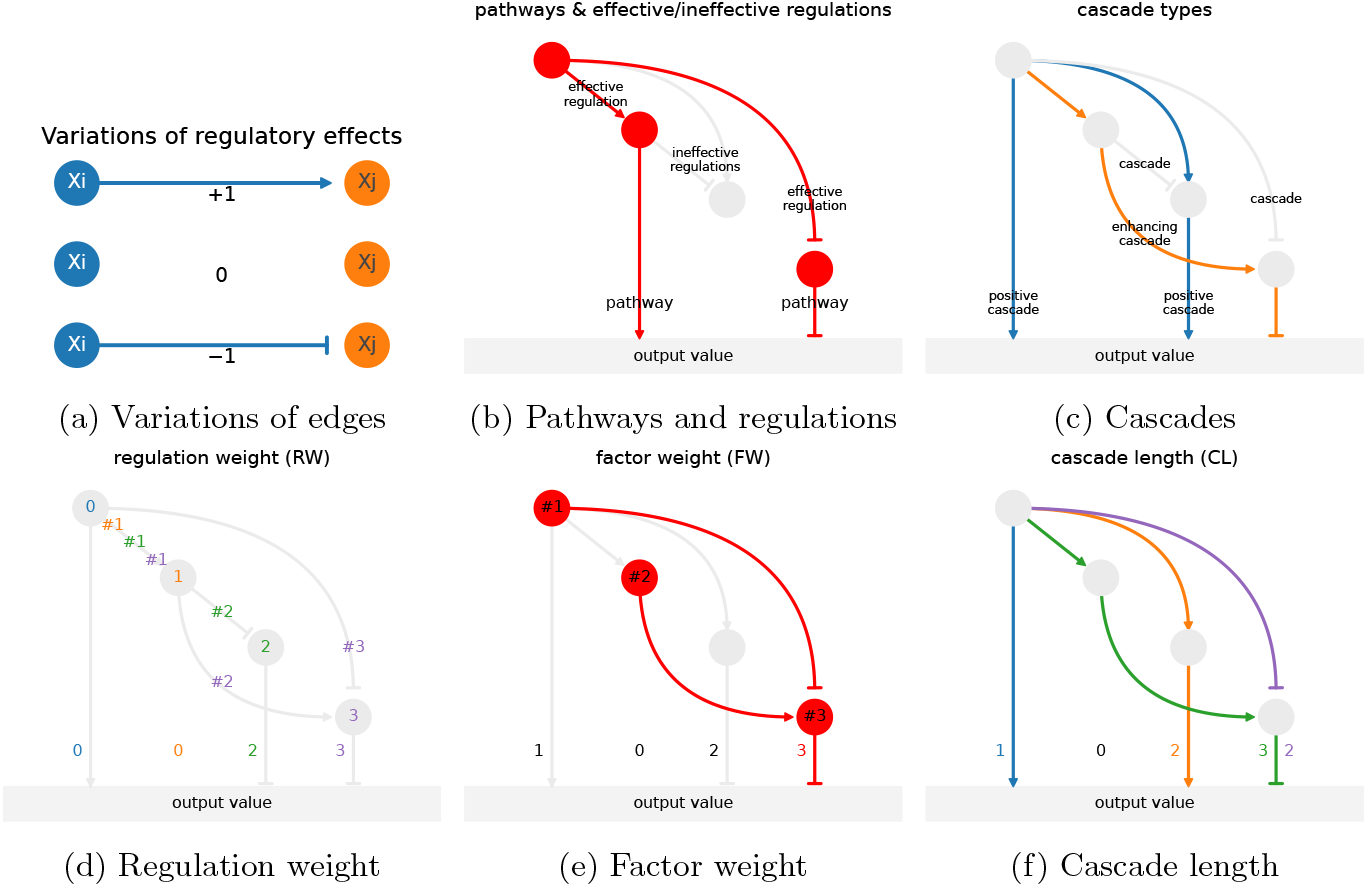
Terminology for simulated model networks. **a**: The three states of edges in the GRNs are considered—having positive (+1), null (0), or negative effects (− 1). **b**: Regulations represent inter-factor edges, whereas pathways are edges between a factor and the output value. A regulation is defined as effective if it can ultimately influence output values via pathways, and ineffective otherwise. **c**: A cascade is defined as a linear series of regulations including the target pathway. An enhancing cascade (orange and blue) consists solely of positive regulations and becomes a positive cascade (blue) if the terminal pathway is also positive. Note that positive cascade is a subdivision of enhancing cascade. **d**: Regulation weight is defined as the number of regulations corresponding to respective pathways. **e**: Factor weight is defined as the number of factors corresponding to respective pathways. **f** : Cascade length is defined as the number of edges (i.e., regulations and pathways) in a cascade corresponding to respective pathways. If a pathway belongs to multiple cascades, its cascade length is determined by the longest cascade containing it (e.g., the rightmost pathway is part of the green cascade, resulting in a cascade length of 3).

To uncover the relationship between these features and experimental design performance, we developed Exhaustive Search Model of four factors (ESM4), a framework generating simulators with 3^10^ exhaustive structural configurations (Figure 5a). For each simulator, we evaluated the performance of C+LOO and PB designs using the *aptitude score*—an extension of Cohen’s *κ* (Methods 4.5.2)—and binned them into “C+LOO-suited” or “PB-suited” classes (Figure 5b). Using the 31 network features, we trained a binary classification gradient boosting decision tree (GBDT) model to predict the “PB-suited” class (Methods 4.8.2–4.8.3). The receiver operating characteristic (ROC) curve shows that the model achieved high predictive performance with an area under the ROC (AUROC) of 0.892 (Figure 5c), and Shapley additive explanations (SHAP) analysis revealed the top 10 features contributing to the prediction (Figure 5d).

**Figure 5.**
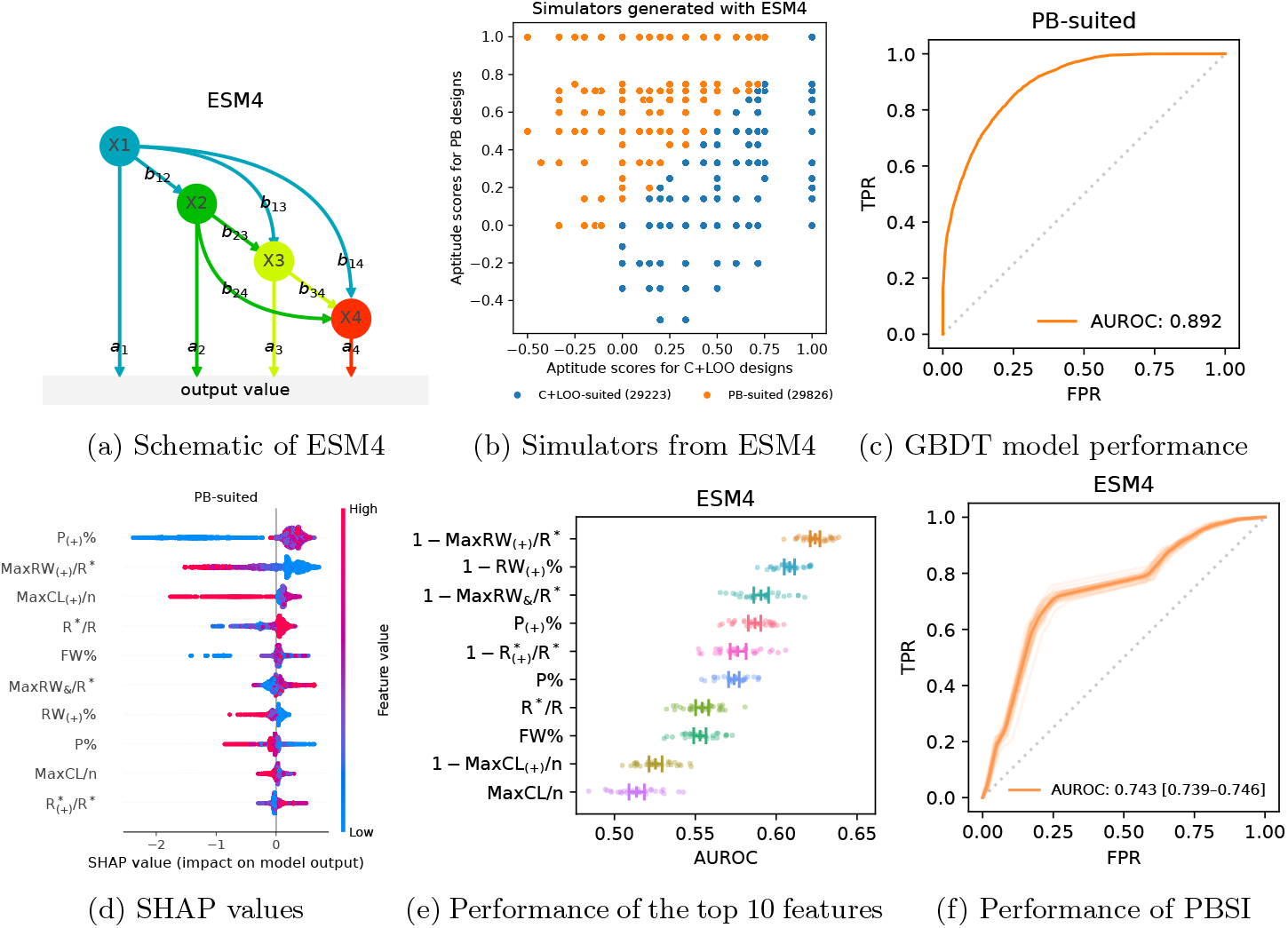
A data-driven approach identified network features associated with high performance of PB relative to C+LOO. **a**: Example network structure generated from ESM4. **b**: Classification of the 3^10^ ESM4 simulators for the ML task. Using aptitude scores for C+LOO 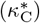 and PB 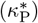, simulators were binned into “C+LOO-suited” (blue; 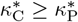) and “PB-suited” (orange; 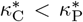) classes. Since C+LOO is the *de facto* standard, the ML model specifically targets the “PB-suited” class to identify scenarios where PB outperforms C+LOO. Numbers in parentheses indicate the count of simulators in each category. **c**: ROC curve of the GBDT model predicting the “PB-suited” class. The horizontal axis represents the false positive rate, and the vertical axis represents the true positive rate. **d**: Beeswarm plot of the top 10 features with the highest mean absolute SHAP values for the “PB-suited” class. Markers are colored according to their corresponding feature values (pink, high; blue, low), and their positions on the horizontal axis indicate the SHAP values. **e**: AUROC scores of the top 10 features with the highest SHAP scores. The error bars indicate the mean and 95% bootstrap CIs. For features associated with the “C+LOO-suited” class, values were inverted so that all AUROC scores were ≥ 0.5. **f** : ROC curve of the PBSI showing its predictive performance (mean AUROC and the 95% bootstrap CI) for the “PB-suited” class.

While the GBDT model was effective, training separate ML models for arbitrary numbers of factors is practically infeasible. We therefore sought a simple, model-free surrogate metric usable for any *n*. Although individual top features identified by SHAP showed only modest discrimination (AUROC ≈ 0.5–0.6; Figure 5e), combining them yielded a robust index. We defined PBSI as the mean of the three features: P_(+)_%, 1 − MaxRW_(+)_*/*R^∗^, and 1 − RW_(+)_% (SI I.1). On the ESM4 dataset, PBSI achieved an AUROC of 0.743, outperforming any single feature (Figure 5f).

To validate the scalability of PBSI beyond four-factor networks, we developed the Exhaustive Search Model of nine factors (ESM9) framework, a validation dataset for nine-factor network structures, finding that PBSI consistently outperformed all other features and composite indices in predicting design suitability (SI I.2).

Conceptually, the three components of PBSI characterize a network topology that avoids the convergence of complex positive interactions—a structural feature where the C+LOO design typically yields robustness. To verify this mechanism, we employed newly designed nine-factor simulators explicitly engineered to test these topological characteristics (Methods 4.8.8). These included Model Π (pure parallel topology yielding high PBSI), Model Δ (entangled topology driving signal amplification, yielding low PBSI), and crucially, Model Σ (entangled topology where amplification is dampened by a negative terminal edge, yielding high PBSI).

Our benchmarking confirmed that mere topological entanglement is insufficient to necessitate the C+LOO design. Instead, it is the signal amplification—triggered by unbroken chains of positive interactions—that causes model misspecification in ANOVA, thereby rendering the orthogonal PB design suboptimal. PBSI successfully captures the presence or absence of this amplification structure, accurately predicting the distinct design preferences across these diverse topologies (SI I.3).

In situations where neither the *de facto* standard C+LOO nor the theoretically high-performing PB is universally superior, PBSI offers a breakthrough: it determines design suitability based solely on GRN topology. This requirement is minimalistic compared to the complexity of a full-scale Perturb-seq experiment.

To demonstrate this applicability, we examined the consistency among biological context, inferred GRN topology, and PBSI scores using unperturbed plain scRNA-seq data. Focusing on TNF signaling (PB-candidate) and Myc response (C+LOO-candidate) as representative drivers of the S-phase (Figures 6a–6c), and filtering genes based on correlation profiles (Figures 6d–6e), we inferred GRNs positioning the S-phase score as the downstream output. Consistent with our simulations, the TNF network exhibited a fragmented parallel architecture with a negative terminal edge suppressing amplification, correctly yielding PBSI *>* 0.6 (Figures 6f–6g)—consistent with the threshold derived from our benchmarking with ESM9 and Models Φ and Ψ (SI I.3). Conversely, the Myc network displayed a convergence of positive interactions culminating in a positive terminal edge—a hallmark of signal amplification— correctly yielding PBSI *<* 0.6 (Figures 6h–6i). Further details and additional cases ensuring reproducibility are provided in SI I.4.

**Figure 6.**
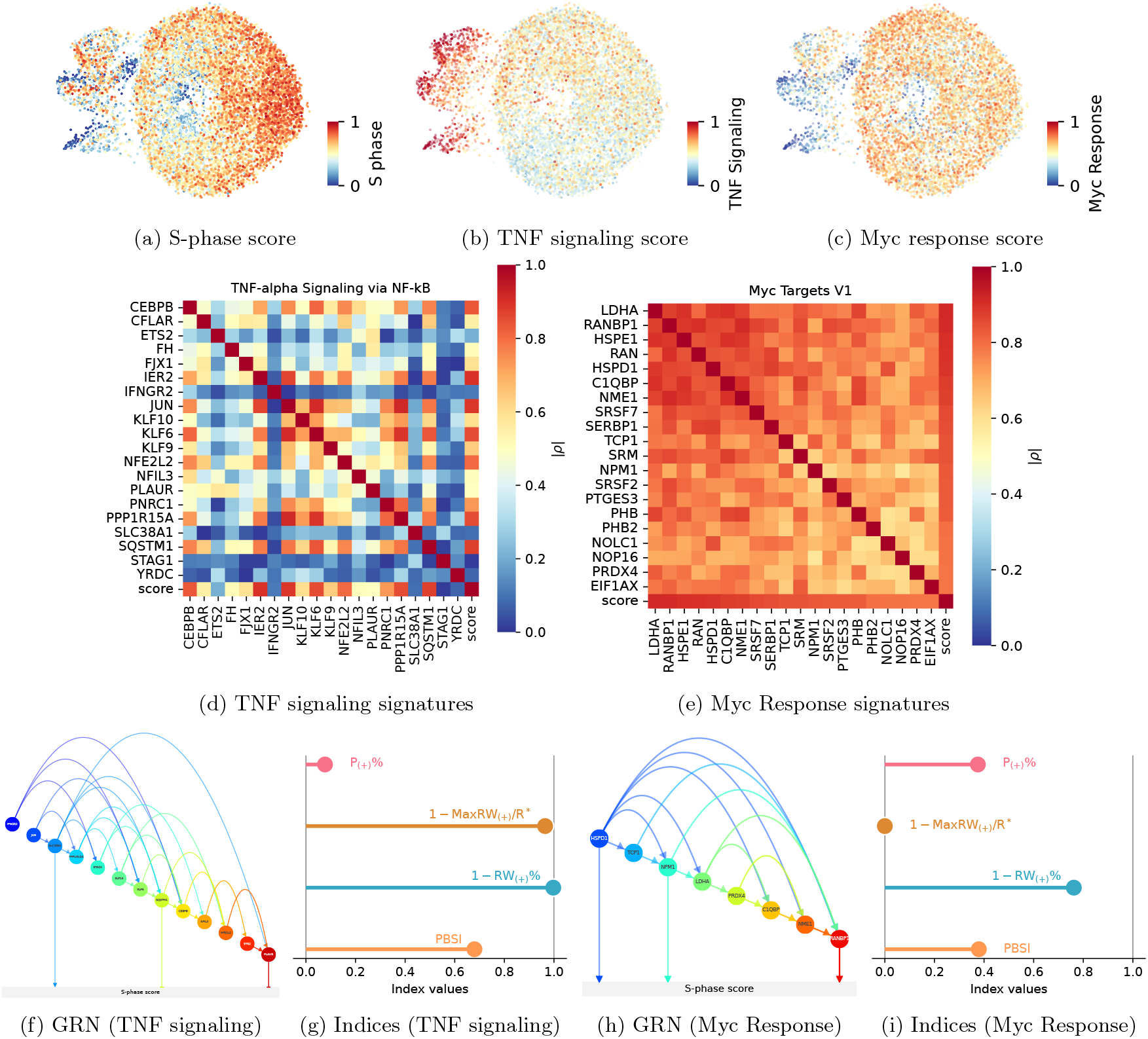
Identification of GRN topologies favoring the PB or C+LOO designs in real-world scRNA-seq data. **a–c**: Uniform Manifold Approximation and Projection (UMAP) plots of K562 cells (non-targeting control condition from Replogle et al. [18]). Cells are colored by the S-phase score (a), TNF signaling score (b), and Myc response score (c). These scores summarize the expression levels of E2F targets, TNF-*α* signaling via NF-*κ*B, and Myc targets V1 genes registered in the MSigDB Hallmark 2020 collection, respectively (see Methods 4.9.2). **d–e**: Heatmaps of the 20 selected genes and phenotypic scores. The absolute Pearson correlation coefficients are shown for the 20 low-correlated genes associated with TNF signaling (d) and the 20 high-correlated genes associated with Myc response (e). The “score” in each panel represents the phenotypic score for the respective biological context. See Methods 4.9.3 for detailed selection criteria. **f–g**: The inferred GRN representing the effect of TNF signaling on the S-phase score (f) and its four topology indices (g). The high PBSI score (*>* 0.6) indicates that the GRN, consisting of low-correlated TNF-*α*-related genes, exhibits fragmented cascades, as reflected by the high values of 1 − MaxRW_(+)_*/*R^∗^ and 1 − RW_(+)_%. Although the topology includes some entangled substructures, the presence of negative terminal edges effectively prevents signal amplification. **h–i**: The inferred GRN representing the effect of Myc response on the S-phase score (h) and its four topology indices (i). The low PBSI score (*<* 0.6) indicates that the GRN, consisting of high-correlated Myc-related genes, exhibits chains of positive gene–gene interactions, as reflected by the low values of 1 − MaxRW_(+)_*/*R^∗^ and 1 − RW_(+)_%. See Methods 4.9.4 for detailed GRN inference methods.

These empirical results suggest that the theoretical dichotomy observed in our simulations—the superiority of orthogonal designs versus their underperformance due to signal amplification-induced model misspecification—is not a hypothetical artifact but a tangible reality in actual biological systems. This demonstration elevates PBSI from a theoretical construct to a powerful, parameter-free “nudge” for experimental design. Notably, PBSI is a concept born neither from biology alone nor from DOE alone, but from their organic integration—a framework we term the *biology-oriented DOE*.

## 3 Discussion

In Perturb-seq, C+LOO has been adopted almost by default across diverse biological systems. By introducing a DOE perspective, we formalized design performance as a quantity to be predicted and compared, in addition to showcasing the PB design as a potential alternative. Our analysis, spanning theoretical simulations and realworld validation, empirically demonstrated that the established dogma of DOE—the unconditional supremacy of orthogonality—does not always hold true in nonlinear, overdispersed biological contexts. This discovery necessitates a paradigm shift towards a “biology-oriented DOE” framework.

Uncovering the universal principles of design suitability through exhaustive search represents a fundamental theoretical advancement in the interdisciplinary integration of biology and DOE. This comprehensive understanding, unattainable solely through empirical comparisons of specific biological contexts, establishes a robust theoretical foundation that transcends the limitations of individual case studies. Operationalizing these principles into the simple PBSI metric and presenting its applicability to real scRNA-seq data served as a critical step to showcase that this theoretical abstraction is indeed anchored in biological reality.

However, true interdisciplinary integration is achieved not when a theory is proposed, but when it survives the complexity of real-world implementation. To ensure that our “biology-oriented DOE” avoids becoming a mere abstraction and effectively navigates the constraints of wetlab reality, we distill our findings into the following actionable policies. Note that these guidelines serve as a high-level navigational compass; for comprehensive technical protocols, mathematical rationales, and epistemological discussions, we direct readers to the extensive documentation in SI J.

- **Candidate gene selection**: Ensure plausibility. For C+LOO, a large candidate list mitigates undesirable factor correlations; we specifically recommend a threshold of *n* ≥ 11 factors (SI J.1).
- **Experimental design selection**: Prioritize C+LOO for systems with significant signal amplification. Otherwise, adopt PB as the robust default. Refer to PBSI for the optimal design. Quantitatively, we recommend adopting the PB design when PBSI *>* 0.6 (SI J.2).
- **Preliminary experiments**: Leverage existing resources—such as public databases and GRN inference tools—to estimate network topology before wet-lab experimentation (SI J.1–J.2). In the near future, large-scale Perturb-seq models will further aid in virtual sample size planning (SI J.4).
- **Strategies without preliminary data**: In the absence of data, hypothesize GRNs using prior domain knowledge (SI J.3).
- **Analytical framework**: Transition from pairwise testing to MLR/ANOVA to enhance statistical power without additional cost (SI J.4).
- **Result interpretation**: Exercise caution in generalization. Mechanistic insights inferred from experiments are not always guaranteed to hold in different experimental settings; interpretation must not be decoupled from experimental premises (SI J.5).
- **Combinatorial effect verification**: While *posthoc* pair addition to C+LOO is common, we caution against its theoretical complexity. Future work should rigorously compare this approach with established DOE methods for interaction analysis (SI J.6).

The application of DOE principles extends beyond individual experiments to the meta-scale of biological research. With the advent of foundation models, the field is increasingly attempting to adapt to the nonlinear and overdispersed complexity of biological systems through sheer data volume and model complexity [19]. While large-scale Perturb-seq models share this approach, they often face challenges in predicting combinatorial effects within extrapolation regimes [20]. Here, strategic experimental design can play a pivotal role by altering how the data space is populated, potentially leading to more efficient model construction than standard datasets. Analogous to physics-informed machine learning, which enhances generalization by constraining model complexity through physical laws [21], largescale biological models could benefit from structural constraints derived from experimental designs and the essential logic of phenotype generation. In this regard, the simulator architecture proposed herein—representing the minimal constitution of biological plausibility—could serve as a foundational testbed for developing such sophisticated, structure-aware models.

In addition to offering a principled basis for design selection, this study holds significant epistemological importance by reframing Perturb-seq as a discrete causal model connecting perturbations to phenotypes, given the inherent “blank time” between intervention and observation. This perspective necessitates a distinction between biological phenomena that can be adequately handled by discrete causal models and those that cannot. In this context, we propose a shift in perspective from strictly “scientific understanding” to “engineering control.” That is, by advancing Perturb-seq as a discrete causal model, we can adopt an engineering approach: optimizing pre-defined phenotypic metrics to control the system, even if the internal regulatory mechanisms remain partially black-boxed.

While biology has traditionally prioritized mechanistic elucidation, the rise of black-box optimization [22] suggests that complete interpretability is not always a necessary condition for controlling complex systems, provided that the system’s behavior is stable and empirically reproducible. Importantly, this engineering stance does not deviate drastically from the traditions of biomedical science; randomized controlled trials, for instance, establish causality without necessarily waiting for a complete mechanistic explanation. Thus, an engineering approach that temporarily bypasses mechanism is compatible with biomedical pragmatism. Fortunately, this does not imply a loss of scientific insight; scRNA-seq data measured during Perturb-seq still allows for the *post-hoc* inference of mechanistic insights, preserving the ability to bridge control and understanding.

Ultimately, perhaps our most significant contribution lies in providing an early case study suggesting that the functional integration of biology and mathematics—a realm where mathematical elegance and simplicity are often revered—requires a fundamental epistemological reframing. We hope that this study serves as a cornerstone for this interdisciplinary integration, accelerating the evolution of multivariate biology in the era of data science.

## 4 Methods

### 4.1 Design matrices and their statistics

#### 4.1.1 An overview of the C+LOO design

The C+LOO design consists of (i) a control trial in which all factors included (i.e., all genes are intact); and (ii) additional trials in which the factors are subtracted one at a time (i.e., single genes are knocked down). This design has traditionally been applied to biological experiments [2]—such as the discovery of the four pluripotency-inducing factors [23].

From a DOE perspective, the C+LOO design can be conceptualized as a variant of the One-Factor-At-a-Time (OFAT) family [3]. However, the status of OFAT is nuanced; while traditionally criticized for its statistical inefficiency and inability to detect interactions, it retains prevalence in engineering practice due to its intuitive simplicity [3].

Unlike generalized OFAT strategies—which often involve multi-level factors without a fixed reference [3]— the C+LOO design represents a specialized framework. It is distinctly characterized by its reliance on a fixed baseline (the “all-intact” control) and a subtractive approach to verify the causal necessity of each factor relative to that baseline.

When we denote the C+LOO design matrix for *n* factors (∀*n* ∈ ℕ) as *C*(*n*) symbolizing the addition and subtraction of factors with ± 1, *C*(5) with five factors for another example is given as follows:

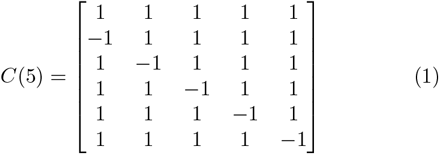

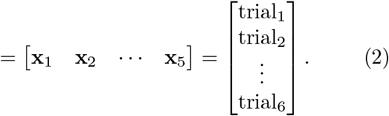

Note that the column vectors (i.e., **x**_*i*_ where ∀*i* ∈ ℕ s.t. 1 ≤ *i* ≤ *n*) indicate the assignments of experimental conditions for the corresponding factors, and the row vectors (i.e., trial_*j*_ where ∀*j* ∈ ℕ s.t. 1 ≤ *j* ≤ *n* + 1) indicate the overall experimental conditions per run. In general, *C*(*n*) can be represented as follows:

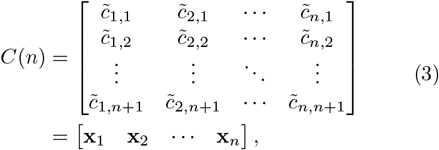

where 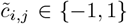 represents experimental conditions (1, addition of the factor; −1, subtraction of the factor). Note that 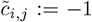 if and only if *j* = *i* + 1; otherwise, 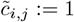.

#### 4.1.2 An overview of the FF design

The experimental designs with complete control experiments—running all possible conditions across 2^*n*^ trials for *n* factors—are called the FF design. In theory, the FF design offers the highest resolution among experimental designs. The FF design is known to show no multicollinearity since any effects from the presence or absence of other factors are canceled out by calculating the mean values. Further information is provided in SI C.1.

#### 4.1.3 An overview of the PB design

The PB design is a popular class of screening design introduced in 1946 [4, 24]. Based on a Hadamard matrix, the PB design assumes a first-degree polynomial model, where the results are explained by a linear combination of the products of factors and their coefficients [25].

The PB design is unique in having only 4 × *k* (*k* ∈ ℕ) trials, making it suitable for up to (4 × *k*) − 1 factors [24]. Since PB designs involve a comparable number of trials to C+LOO designs for a given *n*, we adopted the PB design as the comparison counterparts to the C+LOO design. Regarding the C+LOO design, the total number of trials *n*_max_ for given numbers of replication ∀*N* ∈ ℕ and factors ∀*n* ∈ ℕ is given as follows:

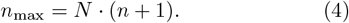

On the other hand, the *n*_max_ that the PB design requires for given ∀*N* ∈ ℕ and ∀*n* ∈ 99 is given as follows:

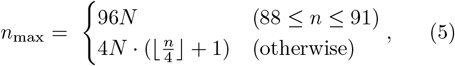

where ⌊⌋ is the floor function. It is noteworthy that the PB design is defined for 4 × *k* (*k* ∈ ℕ) trials up to 100 except for 92 [26].

Further information is provided in SI C.2.

#### 4.1.4 Design matrices, main effects, and interaction terms

We denote a general experimental design *D*(*n, m*) with *n* factors and *m* trials as follows:

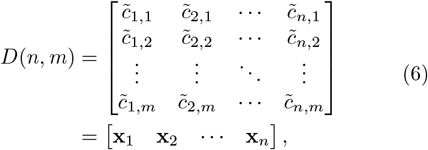

where 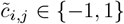 (1 ≤ *i* ≤ *n*, 1 ≤ *j* ≤ *m*) assigns specific conditions. Following the notation of the C+LOO design introduced in Equation (1), we denote the *i*-th column vector corresponding to the *i*-th factor of the design matrix as **x**_*i*_.

The main effects are defined as the independent contributions of individual factors [27]. Although they can be defined for specific levels of given factors (e.g., the main effect 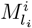 of the *i*-th factor at the *l*_*i*_-th level), the two-level *n*-factor model simply deals the main effect *M*^*i*^ under existence of the *i*-th factor by introducing an assumption 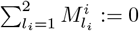 (thus,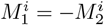).

Similarly, the second-order interactions are defined at respective levels of the given two factors (e.g., the secondorder interaction 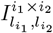 of the *i*_1_-th and *i*_2_-th factors at the 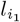-th and 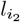-th levels). Since 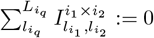 holds for all *q* ∈ { 1, 2 }, the following equation holds in the twolevel *n*-factor models:

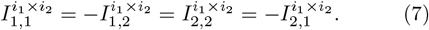

Hence, we exclusively consider the interaction 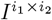 under co-existence or joint absence of the *i*_1_-th and *i*_2_-th factors.

Since the FF design covers complete experimental conditions, the definitions of the main effect *M*^*i*^ of the *i*-th factor and the second-order interaction term 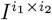 between the *i*_1_-th and *i*_2_-th factors can be derived using the column vectors **x**_1_–**x**_*n*_ of two-level *n*-factor FF designs (1 ≤ *i, i*_1_, *i*_2_ ≤ *n*) with *N* replication as follows:

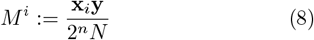

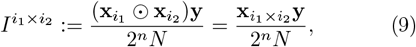

where **y** is the observation vector, ⊙ refers the elementwise (i.e., Hadamard) product of vectors, and 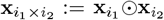 is the column vector for the second-order interaction between the *i*_1_-th and *i*_2_-th factors—indicating the existence and absence of the interaction. Although *M*^*i*^ and 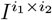 are frequently scaled with 2^*n*−1^*N* or 2^*n*−2^*N* in DOE-contexts, here we align the definition to MLR coefficients for consistency.

Notably, the Pearson correlations among main effects and interaction terms can be calculated with the column vectors—instead of explicitly calculating *M*^*i*^ or 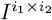 from a concrete **y**. In the field of DOE, this technique is extended to other experimental designs to visualize the characteristics of them without performing actual experiments. Adopting the same idea, we identify the column vectors **x**_*i*_ or **x**_*i×j*_ as a main effect or an interaction in this study for simplicity.

#### 4.1.5 Factor correlations in the C+LOO design

To understand the statistical properties of the C+LOO design, we derived the formula on the correlation coefficients among main effects and interaction terms at various conditions.

As we mentioned in Methods 4.1.4, **x**_*i*_ denotes the main effect of the *i*-th factor, and **x**_*i×j*_ refers to the interaction term between the *i*-th and the *j*-th factors (∀*n* ∈ ℕ, ∀*i, j* ∈ ℕ such that 1 ≤ *i, j* ≤ *n*, and *i* ≠ *j*). We also introduce correlation coefficients *ρ*_*i,j*_ := Corr(**x**_*i*_, **x**_*j*_), *ρ*_*i×j,k*_ := Corr(**x**_*i×j*_, **x**_*k*_), and *ρ*_*i×j,k×l*_ := Corr(**x**_*i×j*_, **x**_*k×l*_) while Corr(,) is the formula for the Pearson correlation coefficient. It is also noteworthy that Corr(,) is symmetrical, therefore, *ρ*_*k,i×j*_ := Corr(**x**_*k*_, **x**_*i×j*_) = *ρ*_*i×j,k*_. Furthermore, *ρ*_*i×j,k*_ = *ρ*_*j×i,k*_ = *ρ*_*k,j×i*_ = *ρ*_*k,i×j*_ because of the symmetry of the Hadamard multiplication in interaction terms.

To evaluate how much of correlations caused in *C*(*n*) at different levels of *n*, we calculated the correlation coefficients *ρ*_*i,j*_, *ρ*_*i×j,i*_, *ρ*_*i×j,k*_, *ρ*_*i×j,i×l*_, and *ρ*_*i×j,k×l*_ (∀*n* ∈ ℕ, ∀*i, j, k, l* ∈ ℕ such that 1 ≤ *i, j, k, l* ≤ *n* and (*i* − *j*) · · (*i* − *k*) · (*i* − *l*) · (*j* − *k*) · (*j* − *l*) · (*k* − *l*) ≠ 0) at specific conditions as follows:

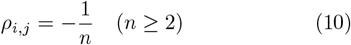

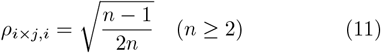

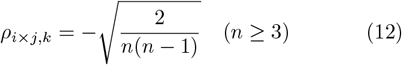

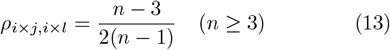

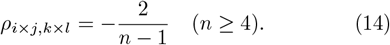

Notably, here we omitted *ρ*_*i,i*_ and *ρ*_*i×j,i×j*_ since they are constant. More rigorous descriptions are provided in SI A–B.

We also calculated the limit values of them to have the following equations:

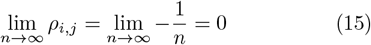

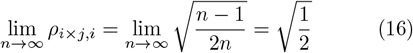

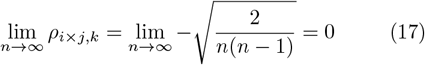

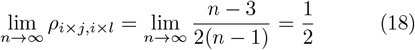

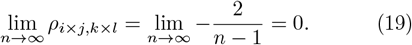

Note that the limit operations are all considered as discrete since the correlation functions are defined for *n* ∈ ℕ.

#### 4.1.6 Visualization of design matrices, correlation heatmaps, and the correlation curves

We created the C+LOO and FF design matrices using NumPy [28] and pandas [29]. We generated the PB design matrix using pyDOE [30] and other Python packages. The column vectors for interaction terms were calculated with pandas and other Python packages.

We created the design matrices and correlation heatmaps with NumPy, pandas and other Python packages. We also visualized the five correlation functions by evaluating Equations (10)–(14) for values ranging from 2 to 12 using NumPy.

#### 4.1.7 Model matrices

The model matrix is a matrix representing independent variables (including the intercept) of the model. As a design matrix represents the states of the manipulated variables, it can serve as a part of the model matrix for MLR models.

For example, the model matrix *X* for the “additive model”—an MLR model comprising main effects and a constant term (Methods 4.4.2)—based on an experimental design *D*(*n, m*) can be given as follows:

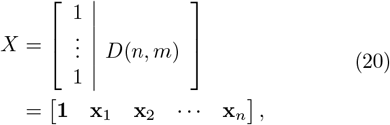

where **1** is the *m*-dimensional vector which elements are all 1.

#### 4.1.8 Condition numbers and VIF

We calculated condition numbers and VIF values of a factor based on the model matrices of additive models (Methods 4.1.7 and 4.4.2). Then, we also plotted them using NumPy, statsmodels [31], and other Python packages. Notably, all factors in an additive model show the same VIF values when the model matrix is based on the C+LOO or PB designs. Thus, we reported VIF values of one factor per *n*.

### 4.2 Formalization of Perturb-seq

To rigorously evaluate the impact of experimental design, we first formalized the theoretical framework of Perturbseq and delineated the scope of biological systems where DOE principles are applicable.

First, we define Perturb-seq as a framework that establishes discrete causal models mapping perturbation inputs (e.g., gene KD) to phenotypic outputs (scalar metrics derived from high-dimensional scRNA-seq data) (Figure S3). This approach inherently treats the intermediate molecular mechanisms between perturbation and observation as a black box, inferring causal relationships by contrasting perturbed states against an “all-gene intact” baseline.

Second, we distinguish between two classes of biological systems based on the temporal dynamics of their underlying GRNs during the interval between perturbation and observation:

- **Systems with time-varying GRN topology:** Systems where the GRN structure itself evolves in a condition-dependent manner (e.g., sign reversals, rewiring, or module coupling/decoupling as shown in Figure S4a). In these systems, the static snapshot nature of scRNA-seq leads to a loss of traceability, making mechanistic inference theoretically intractable without time-series data.
- **Systems with static GRN topology:** Systems where the GRN structure remains static (time-invariant), and perturbations manifest solely as quantitative changes in gene expression levels within a fixed topology (Figure S4b).

We posit that the discrete causal modeling used in Perturb-seq is primarily robust in systems with static GRN topology, where the structural invariance ensures that the input–response relationship can be reliably approximated. Consequently, this study restricts its scope to such static systems to isolate the effects of experimental design from the confounding factors of network evolution.

A detailed derivation of this formalization and illustrative examples of GRN dynamics are provided in SI D.1.

### 4.3 Minimalist simulators

Perturb-seq is primarily utilized to directly link perturbed genes with phenotypes, often bypassing the intricacies of underlying regulatory networks to interpret transcriptomic changes (SI D.1). However, to gain deeper insights into how experimental design shapes observable outcomes, it is crucial to dissect the impact of perturbations on signaling cascades in detail.

Given the difficulty of isolating the true effects of experimental design choices in real-world data (where the governing equations remain unknown), we constructed minimalist simulators of signaling cascades. These simulators provide tangible insights into how differences in experimental design alter the biological interpretation of Perturb-seq (see SI D.2 for a detailed rationale).

All minimalist simulators developed in this study share a unified architecture: gene–gene regulatory relationships are represented as a directed acyclic graph (DAG) mimicking transcriptional control, which ultimately regulates a scalar output value representing the phenotype (corresponding to single-gene markers or phenotypic scores). To balance computational efficiency with biological plausibility, we employed the rectified linear unit (ReLU) activation function. This choice introduces nonlinearity—preventing the output from reducing to a simple linear sum of gene expression levels—and effectively emulates the threshold-like behavior observed in biological systems while keeping computational costs low [32–35].

Consequently, our simulator exhibits a structure analogous to a real-valued extension of Boolean networks [36], similar to those employed in mathematical models of cellsignaling pathways [17]. Furthermore, model parameters were initialized via random sampling from log-normal distributions to recapitulate the long-tail distributions frequently observed in real-world biology.

#### 4.3.1 Model Φ: a basic model simulating biological mechanisms

Model Φ is a toy simulation of Perturb-seq analysis applied to complex gene regulation, involving nine factors (X1–X9) and multiple regulatory routes that influence the output value either directly or indirectly. Among the models presented in this study, it serves as the most neutral baseline compared to Models Ψ and Λ, given its balanced configuration of pathways.

Let *n*_max_ denote the total number of trials—e.g., *n*_max_ = 10*N* for *N* replicates of *C*(9). Define the following for all *i, j* ∈ ℕ such that 1 ≤ *i* ≤ 9 and 1 ≤ *j* ≤ *n*_max_:

- *x*_*i,j*_: observed value of factor *i* in trial *j*
- *y*_*j*_: output value of trial *j*
- *v*_*i*_ ∼ Lognormal(1, (0.8)^2^): latent constant of factor *i*
- *c*_*i,j*_ ∈ { 0, 1 }: perturbation indicator (0, factor included; 1, factor removed)
- *K* := {1, 2, 6, 7, 9}
- *L* := {27, 37, 45, 57, 67, 89}
- *a*_*k*_ ∼ Lognormal(2, (0.3)^2^) for *k* ∈ *K*: direct coefficients on *y*_*j*_
- *b*_*ℓ*_ ∼ Lognormal(1, (0.5)^2^) for *ℓ* ∈ *L*: regulatory coefficients for inter-factor influence
- *ε*_*i,j*_, *ε*_*y,j*_ ∼ N (0, *σ*^2^): Gaussian noise for *x*_*i,j*_ and output *y*_*j*_
- ReLU: ReLU function, defined as *f* (*x*) = max(0, *x*)

Here, 𝒩 (*µ, σ*^2^) denotes the normal distribution, and Lognormal(*µ, σ*^2^) denotes a log-normal distribution such that ln *X* ∼ 𝒩 (*µ, σ*^2^). The variables *v*_*i*_, *a*_*k*_, and *b*_*ℓ*_ are shared across trials and were generated using NumPy with a fixed random seed to ensure reproducibility and positivity. Note that the index *ℓ* in *b*_*ℓ*_ refers to a specific regulatory connection between factors (e.g., *b*_27_ affects the influence from X2 to X7), rather than to a particular factor itself.

The model equations are defined as:

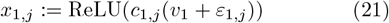

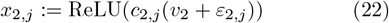

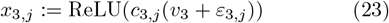

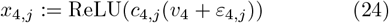

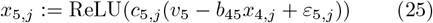

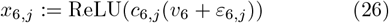

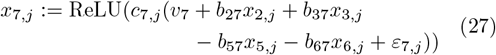

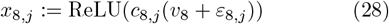

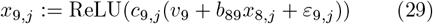

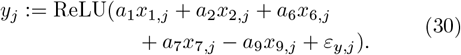

#### 4.3.2 Model Ψ: a highly complex regulatory circuit of interactions

Model Ψ represents a more intricate network of interactions than Model Φ. Using the similar notations introduced in Model Φ, we define the following for all *i, j* ∈ ℕ such that 1 ≤ *i* ≤ 9 and 1 ≤ *j* ≤ *n*_max_:

- *x*_*i,j*_, *y*_*j*_, *c*_*i,j*_, *ε*_*i,j*_, *ε*_*y,j*_, ReLU: identical to those in Model Φ
- *K* := {1, 2, 7, 8}
- *L* := {34, 35, 36, 37, 45, 46, 47, 56, 57, 67, 78, 89}
- *a*_*k*_ ∼ Lognormal(2, (0.3)^2^) for *k* ∈ *K*
- *b*_*ℓ*_ ∼ Lognormal(1, (0.5)^2^) for *ℓ* ∈ *L*

As in Model Φ, the parameters *v*_*i*_, *a*_*k*_, and *b*_*ℓ*_ are shared across trials and were sampled using NumPy with a fixed random seed.

The model equations are given by:

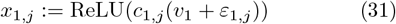

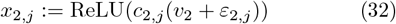

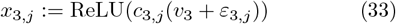

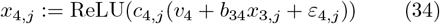

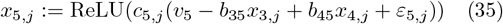

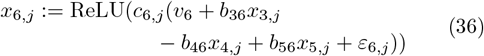

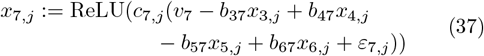

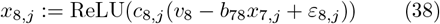

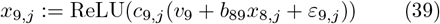

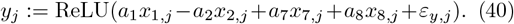

#### 4.3.3 Model Λ: a model with poorly curated candidates

Model Λ mimics a situation where the candidate genes are poorly filtered so that the majority of them shows no influence on the target phenomenon. With the similar notations introduced in Model Ψ, we define the following for all *i, j* ∈ ℕ such that 1 ≤ *i* ≤ 9 and 1 ≤ *j* ≤ *n*_max_:

- *x*_*i,j*_, *y*_*j*_, *c*_*i,j*_, *ε*_*i,j*_, *ε*_*y,j*_, ReLU: identical to those in Models Φ and Ψ
- *K* := {1, 2}
- *L* := {45, 46, 47, 57, 67, 78, 79}
- *a*_*k*_ ∼ Lognormal(2, (0.3)^2^) for *k* ∈ *K*
- *b*_*ℓ*_ ∼ Lognormal(1, (0.5)^2^) for *ℓ* ∈ *L*

As in Models Φ and Ψ, the parameters *v*_*i*_, *a*_*k*_, and *b*_*ℓ*_ are shared across trials and were sampled using NumPy with a fixed random seed.

The model equations are given by:

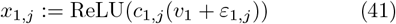

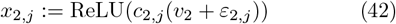

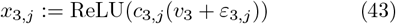

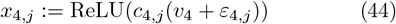

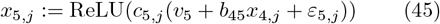

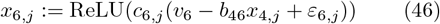

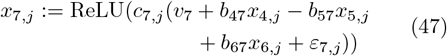

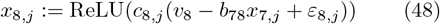

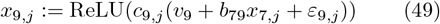

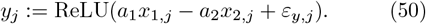

#### 4.3.4 Implementation

We generated random variables using NumPy, setting different seeds for pseudorandom numbers in different trials. By default, Gaussian noise followed 𝒩 (0, 1). To investigate the impact of noise intensity, we also evaluated *σ* ∈ {0.5, 2, 4}.

Regarding replication, we simply repeated the experiments sequentially without shuffling. In real-world experiments, it is essential to randomize the execution order—specifically, the row order of the design matrix— to minimize the impact of covariates associated with the trial sequence [37] (e.g., improvements in technical proficiency or workflow efficiency). However, for simplicity, we assumed no such covariates were associated with the order of experiments and treated the output values as independent and identically distributed variables.

#### 4.3.5 Validation of the biological relevance of the simulators

Although Models Φ, Ψ, and Λ were constructed without any specific network motifs, we examined whether these simple networks could reproduce the nonlinear and overdispersed characteristics of real-world biological systems. Specifically, we tested whether the output values of these simulators followed the TPL, a log-log relationship between means and variances observed across a variety of biological systems, including scRNA-seq data [11–15].

As detailed in SI D.3, we performed FF-based simulations for each model (*N* = 10), varying the Gaussian noise levels (*σ* ∈ { 0.5, 1, 2, 4 }) to generate output values under a wide range of experimental conditions. From the generated data, the means and variances of output values were calculated within each experimental condition defined by the FF design. We then fitted linear regression models to the log-transformed means and variances to obtain the coefficient of determination (*R*^2^) and the scaling exponent (*β*), which represent the degree of conformity to the TPL.

The implementation was carried out in Python using packages including scikit-learn [38].

### 4.4 Statistical analyses

#### 4.4.1 Statistical testing for pairwise comparisons

In SI E and G, we performed Dunnett’s test [9] using the Python package, SciPy [39], with the control condition defined as the trial including all factors. Statistical significance was set at *α* = 0.05; *p*-values were reported as ∗ for *p <* 0.05, ∗∗ for *p <* 0.01, ∗∗∗ for *p <* 0.001, and N.S. for not significant. Plots were generated with Python packages, and we displayed error bars representing the mean ± SD.

We also conducted per-comparison power analysis for *k* treatment groups. The required quantities—the residual degrees of freedom *ϕ*, noncentrality parameters *λ*_*i*_, correlation matrix *R*, test statistics *T*_*i*_, and power 1 − *β*_*i*_— were defined as follows:

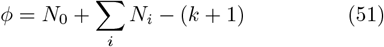

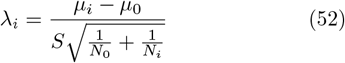

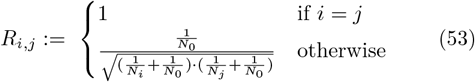

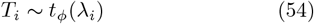

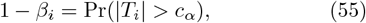

where 1 ≤ *i, j* ≤ *k*; *N*_0_ and *N*_*i*_ are the sample sizes in the control and the *i*-th treatment group, respectively; *µ*_0_ and *µ*_*i*_ are the corresponding group means; *S* denotes the common standard deviation (pooled across groups); *t*_*ϕ*_(*λ*_*i*_) is the noncentral *t*-distribution with *ϕ* degrees of freedom and noncentrality *λ*_*i*_; Pr denotes the probability function; and *c*_*α*_ is the two-sided Dunnett critical value that controls the family-wise error rate at *α*.

For estimation, *µ*_0_ and *µ*_*i*_ were taken as the sample group means. The common standard deviation was estimated as

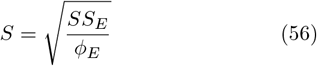

(see Equations (71)–(72) for the definition of *SS*_*E*_ and *ϕ*_*E*_). The Dunnett two-sided critical value *c*_*α*_ [9] corresponding to *α* = 0.05 was obtained using the R package mvtnorm [40] by inverting the central multivariate *t*-distribution with correlation matrix *R*.

#### 4.4.2 Additive models

Under the *additive model* —a multivariate model assuming no interaction terms—we modeled *n* factors at a replication level *N* (∀*n, N* ∈ ℕ), the output value 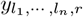 when ∀*i*-th factors are assigned to their *l*_*i*_-th level (∀*i*, ∃*L*_*i*_, ∀*l*_*i*_ ∈ ℕ such that 1 ≤ *i* ≤ *n* and 1 ≤ *l*_*i*_ ≤ *L*_*i*_) at replication index *r* (1 ≤ *r* ≤ *N*) is modeled as follows:

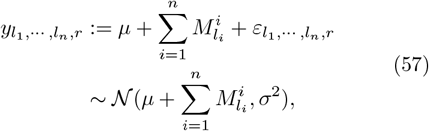

where *µ* is the population mean, 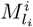 is the main effect of the *i*-th factor at the *l*_*i*_-th level, and 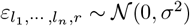 is the Gaussian noise term.

As described in Methods 4.1.4, main effects 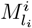 can be simplified to *M*^*i*^ to have the following:

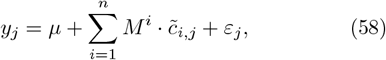

where 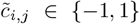 is the element of a design matrix (refer to Equation (6)) assigning specific experimental conditions. Notably, 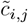 can be introduced by rescaling *c*_*i,j*_—the perturbation indicators introduced in Models Φ, Ψ, and Λ—with 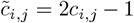. Moreover, *y*_*j*_ and *ε*_*j*_ are equivalent to 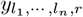 or 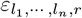 introduced in Equation (57), but indexed with the trial number *j*.

With a design matrix *D*(*n, m*) = [**x**_1_ ··· **x**_*n*_], Equation (58) can be represented as follows:

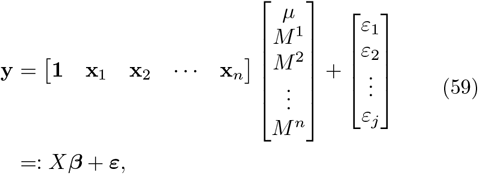

where **y** is the vector for output values *y*_*j*_, *X* is the model matrix, and ***β*** is the coefficient vector containing *µ* and *M*^*i*^. ***ε*** is the residual error vector following the *m*-dimensional Gaussian distribution 𝒩_*m*_(**0**, *σ*^2^**I**_*m*_), where **I**_*m*_ denotes the *m* × *m* identity matrix.

#### 4.4.3 MLR

To visualize the results from additive models with a simple method, we created ordinary least square MLR models referring Equation (59). By minimizing the mean squared error, the estimated coefficient vector 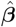 is determined as follows:

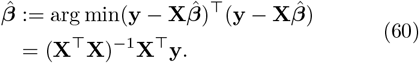

We implemented MLR models with statsmodels and visualized the MLR coefficients with bar charts using Python packages.

#### 4.4.4 ANOVA

We adopted Type II ANOVA, as recommended by Langsrud (2003) [41]. Since interaction terms are undetectable in OFAT designs (the superfamily of C+LOO; see Methods 4.1.1) [3] and PB designs rely on a first-degree polynomial model [25], Type II ANOVA is deemed appropriate for these contexts as it allows for analysis without postulating interaction terms [41].

To perform Type II ANOVA, we introduce three different model matrices *X*_Ω_, *X*_*i*_, and *X*_Ω−*i*_ as follows:

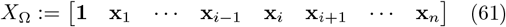

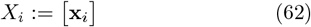

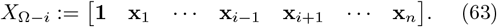

Notably, *X*_Ω_ is the model matrix of additive models, whereas *X*_*i*_ and *X*_Ω−*i*_ represent the partial models of *X*_Ω_ which parameters are exclusive and complementary to each other.

With these model matrices, the additive model can be derived as follows:

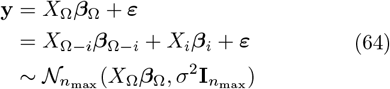

where ***β***_Ω_, ***β***_Ω−*i*_, and ***β***_*i*_ are the coefficient vectors for the respective model matrices.

Under Type II ANOVA, the sum of squares for the *i*-th factor *SS*(*X*_*i*_) are indirectly derived from the total sum of squares *SS*(*X*_Ω_) subtracted with *SS*(*X*_Ω−*i*_) for given observations **y** as follows:

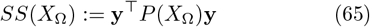

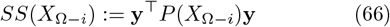

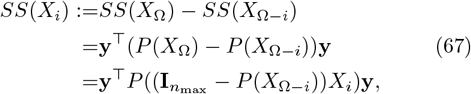

where *P* (*A*) := *A*(*A*^⊤^*A*)^−1^*A*^⊤^ denotes the orthogonal projection matrix of an arbitrary matrix *A*.

Regarding *SS*(*X*_*i*_), the following equations hold:

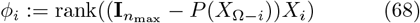

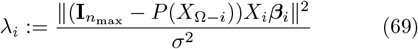

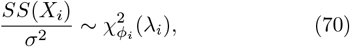

where 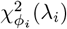 denotes a noncentral chi-squared distribution with *ϕ*_*i*_ degrees of freedom and a noncentrality parameter *λ*_*i*_.

Similarly, the error sum of squares *SS*_*E*_ is defined as follows:

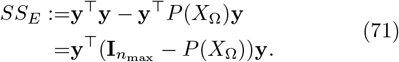

In practice, the noncentrality parameter *λ*_*E*_ for *SS*_*E*_*/σ*^2^ is assumed to be zero to avoid the doubly noncentral *F* distribution regarding the *F*-statistics (refer to SI F). Therefore, the following equations hold:

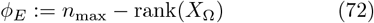

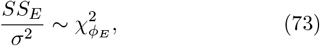

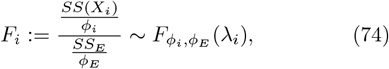

where 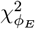 denotes a chi-squared distribution with *ϕ*_*E*_ degrees of freedom, *F*_*i*_ is the *F*-statistic for the *i*-th main effect, and 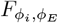 (*λ*_*i*_) denotes the noncentral *F*-distribution with *ϕ*_*i*_ and *ϕ*_*E*_ degrees of freedom and a noncentrality parameter *λ*_*i*_.

In the statistical testing on the *i*-th factor main effect, the null hypothesis *H*_0_ assumes ***β***_*i*_ := [_*M*_^*i*^] = **0**. Hence, the *p*-values are calculated under assumption that *λ*_*i*_ = 0.

In contrast, in the power analysis of the statistical testing, the statistical power 1 − *β* is calculated under the noncentral *F*-distribution. More precisely, the upper probability of the noncentral *F*-distribution is calculated for an *F* such that the upper probability of the corresponding *F*-distribution equals the *α* used in the statistical testing. The noncentrality parameter *λ*_*i*_ was estimated as follows:

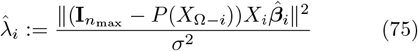

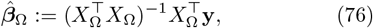

where 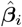 corresponds to the *i*-th element (or sub-vector) of the estimated coefficient vector 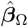.

Notably, the implementation described above is mathematically equivalent to the formulas derived from the definitions of Type II ANOVA in Langsrud (2003) [41] and the general linear model framework in Muller and Peterson (1984) [42].

We implemented Type II ANOVA, power analysis, and visualization with Python packages including statsmodels and SciPy.

#### 4.4.5 Visualization of MLR/ANOVA results

In Figures S6b–S6c, we visualized the results from MLR and ANOVA for Model Φ-based additive models using lollipop plots. To visualize both the direction and magnitude of the factor effects, we calculated the signed partial eta-squared values 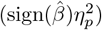, defined as the product of the sign of the MLR coefficient 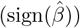 and the partialeta squared 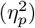 [43]. The partial eta-squared for the *i*-th factor 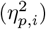 is defined as follows:

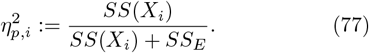

We computed the MLR coefficients and sums of squares using statsmodels and derived the signed metrics using NumPy.

#### 4.4.6 Main effect estimation

After generating simulated results based on the FF design with 100 different random seeds, we calculated the main effects according to Equation (8) and estimated the bootstrap 95% CIs with SciPy, setting the number of bootstrapping *n*_boot_ = 10000. Then we interpreted the results based on the CIs as follows:

- “Up” if the lower bound of the CI is above 0
- “N.S.” the CI includes 0
- “Down” if the upper bound of the CI is below 0

We leveraged these labels as the ground truth when comparing the experimental design performance.

#### 4.4.7 Half-normal plot

The half-normal plot shown in Figure S6e is a graphical method to visualize the factor effects which is versatile even when ANOVA does not work—e.g., *ϕ*_*E*_ = 0. The effects are suggested to be significant when the absolute values of the standardized effects are larger than the halfnormal quantiles [44, 45].

We created a half-normal plot for the C+LOO-based Model Φ results at *N* = 1. The half-normal quantiles were calculated using SciPy implementation, and the standardized effects were computed with NumPy. Then, we visualized these results with Matplotlib and other Python packages.

### 4.5 Performance evaluation metrices

#### 4.5.1 Weighted Cohen’s *κ*

When evaluating the main effects in the simulators from generated results, we applied the following criteria to interpret the multivariate models: the effect of a factor is positive if the MLR coefficient is positive and ANOVA suggests *p <* 0.05; negative if the coefficient is negative and *p <* 0.05; and N.S. if *p* ≥ 0.05.

We quantified the agreement between result interpretations by calculating weighted Cohen’s *κ* coefficients relative to the ground truth labels (Methods 4.4.6). Although Cohen’s *κ* is limited to evaluating categorical concordance and does not capture the quantitative magnitude of effects, we adopted this metric to prioritize qualitative accuracy. In exploratory biological experiments— particularly in domains where mechanistic consensus remains unestablished—the primary objective is often to elucidate the Boolean topology of the governing mechanisms or GRNs. Consequently, we defined model performance based on the fidelity of qualitative classification rather than quantitative precision.

To reduce strictness in identifying statistically significant effects as N.S. (or vice versa) and to relatively penalize predicting effects in opposite directions, we applied linear weights (0, 0.5, and 1) when calculating *κ* values. We implemented this method using Python packages, including NumPy, pandas, and scikit-learn, and visualized the results with Matplotlib and seaborn.

#### 4.5.2 Aptitude score

Cohen’s *κ* is defined as follows [46]:

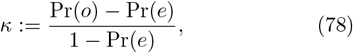

where Pr(*o*) and Pr(*e*) denote the observed and expected agreement, respectively. As shown in Equation (78), *κ* is undefined when Pr(*e*) = 1, a limitation that also applies to Weighted Cohen’s *κ*.

To ensure robust evaluation of experimental design performance under diverse conditions, we extended the standard weighted Cohen’s *κ* used in Models Φ, Ψ, and Λ by introducing the *aptitude score*, which returns raw accuracy values in cases where *κ* is undefined.

To compute aptitude scores, we measured the agreement between the results predicted by the multivariate model based on the experimental design of interest (C+LOO or PB) and the ground truth labels.

We implemented this metric using Python packages, including NumPy, pandas, and scikit-learn.

### 4.6 The D-criterion and D-optimization

With respect to the model matrix *X* (introduced in Methods 4.1.7), *X*^⊤^*X* is called the information matrix, and (*X*^⊤^*X*)^−1^ is the dispersion matrix [16]. The determinant of the dispersion matrix det(*X*^⊤^*X*)^−1^ is called the D-criterion.

D-optimization is a process of choosing the model matrix, as well as the design matrix which forms the model matrix, that minimizes the D-criterion. Such matrices are considered optimal for experimental designs because the lowest D-criterion indicates that the coefficients of the multivariate model are as independent of each other as possible [16]. Note that minimizing the determinant of the dispersion matrix are equivalent to maximizing the determinant of the information matrix [16].

We visualized the D-criterion values with Python packages including Matplotlib, NumPy, seaborn, and statsmodels. We performed D-optimization using an R package AlgDesign [47] and visualized the results with Python packages.

### 4.7 Terminology and features for simulator network structures

#### 4.7.1 Terminology for feature engineering

To numerically characterize network structures, here we introduce a terminology to describe network structures of simulated models. *pathway, regulation*, and *cascade*. A *pathway* refers to an edge that directly leads to the output value, while a *regulation* denotes an edge connecting two factors. Each edge can take one of three forms—positive, null, or negative (Figure 4a)—resulting in a power-of-three number of possible network structures for the simulated models. A *cascade* is a chain of a pathway and one or more regulations that are linearly connected. Notably, a single pathway without any upstream regulations is also a cascade.

Furthermore, if a regulation is directly or indirectly connected to a pathway—that is, if it forms part of a cascade—we refer to it as an *effective* regulation. In contrast, any regulation that lacks a downstream pathway is considered *ineffective* (Figure 4b).

To highlight impactful cascades that influence the output value, we introduce *enhancing cascades* and *positive cascades* (Figure 4c). A cascade is considered *enhancing* if it either has no regulations or if all existing regulations are positive. An enhancing cascade is *positive* when the pathway is also positive. Thus, all positive cascades are enhancing. Having at least one negative regulation, the effect of the pathway in non-enhancing cascades is directly or indirectly downregulated within the cascade itself.

Then, for all pathways, we define *regulation weight* and *factor weight*. A *regulation weight* is a lump-sum number of regulations belonging to cascades with the pathway of interest. For example, we present a graphical example in Figure 4d. The most upstream factor has zero regulation weight. The factor marked with orange has a regulation from the blue one but any pathway downstream. In such a case, regulation weight is not defined. For the sake of consistency, we assign zero instead. The factor marked with green has two upstream regulations—edges from the blue one to the orange one, and from the orange one to the green one—hence, the regulation weight becomes two. Having three regulations upstream, the purple factor has a pathway with regulation weight of three. Similarly, a *factor weight* is the number of factors involving to the pathway of interest. As Figure 4e shows, factor weight for the pathway in red is three since there are three factors directly or indirectly involving the pathway. Notably, we also assigned zero factor weight when a factor has no pathway.

As an alternative feature, we introduce *cascade length*, a total number of edges in the cascade containing the pathway of interest. Since a pathway can be a part of multiple cascades, the largest cascade represents the pathway in such cases. Thus, the cascade length for the pathway on the right in Figure 4f is three.

#### 4.7.2 The 31 features of the simulator network structures

Based on the terminology introduced above, we defined 31 features—including pathway-related, regulation-related, regulation weight-related, factor weight-related, and cascade length-related features—for a network with *n* factors, *p* pathways, and *r* regulations (*n, r, p* ∈ ℤ such that *n* ≥ 1 and *r, p* ≥ 0). Notably, we defined the values to always fall within the range [0, 1] for all *n* to gain generalizability for any number of factors.

As pathway-related features, we introduce *pathway coverage* P%, *positive pathway coverage* P_(+)_%, *negative pathway coverage* P_(−)_%, and *pathway positivity* P_(+)_*/*P, and *pathway negativity* P_(−)_*/*P as follows:

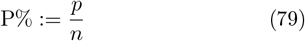

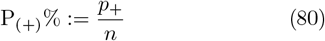

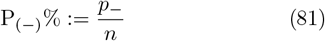

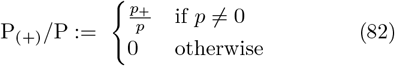

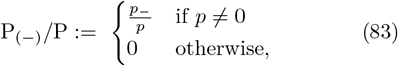

where *p*_+_ and *p*_−_ denote the number of positive and negative pathways, respectively. The metrics P%, P_(+)_%, and P_(−)_% reflect the abundance of all, positive, and negative pathways, respectively, and are computed as ratios normalized by the number of factors *n*, which defines the maximum possible number of pathways. In contrast, P_(+)_*/*P and P_(−)_*/*P represent the proportions of positive and negative pathways relative to the total number of pathways.

Subsequently, we introduce regulation-related features—namely, *regulation coverage* R%, *positive regulation coverage* R_(+)_%, *negative regulation coverage* R_(−)_%, *regulation positivity* R_(+)_*/*R, *regulation negativity* R_(−)_*/*R, *regulation effectivity* R^∗^*/*R, *effective regulation positivity* 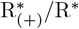, and *effective regulation negativity* 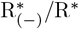 —as follows:

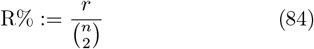

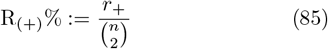

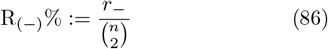

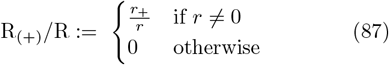

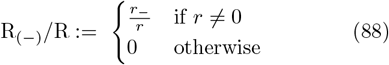

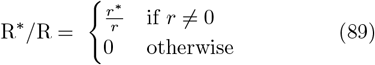

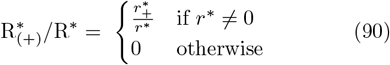

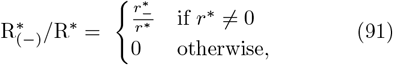

where *r*_+_, *r*_−_, *r*^∗^, 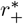, and 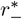 denote the number of positive, negative, effective, positive effective, and negative effective regulations, respectively. Given that the theoretical maximum number of regulations for *n* factors is 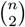, the values R%, R_(+)_ %, and R_(−)_ % are normalized by this quantity. The ratios R_(+)_*/*R, R_(−)_*/*R, and R^∗^*/*R are computed relative to the total number of regulations, while 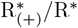 and 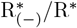 reflect the proportions of positive and negative edges among effective regulations, respectively.

Then, we introduce regulation weight-related features: *maximum regulation density* MaxRW*/*R^∗^, *regulation weight coverage* RW%, *maximum positive regulation density* MaxRW_(+)_*/*R^∗^, *positive regulation weight coverage* RW_(+)_%, *maximum enhancing regulation density* MaxRW_&_*/*R^∗^, and *enhancing regulation weight coverage* RW_&_%.

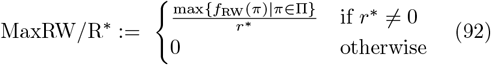

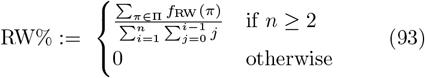

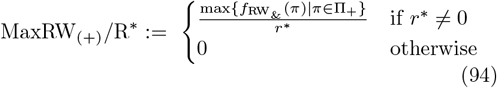

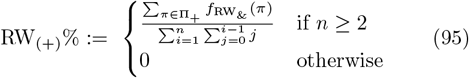

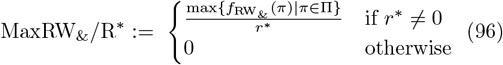

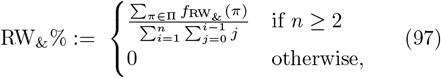

where Π and Π_+_ denote the sets of all pathways and positive pathways, respectively; *f*_RW_ and 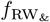 are functions that return the total regulation weights of a given pathway or those restricted to enhancing cascades (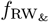 returns zero for pathways of non-enhancing cascades); and max denotes the function returning the maximum value in each set. The values MaxRW*/*R^∗^, MaxRW_(+)_*/*R^∗^, and MaxRW_&_*/*R^∗^ respectively represent the impact of maximum regulation weight among cascades, positive cascades, and effective cascades, divided by the number of effective regulations. RW%, RW_(+)_%, and RW_&_% represent the coverages of regulation weights of respective categories normalized by the theoretical maximum number 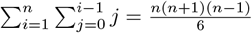.

Similarly, factor weight-related factors, such as *maximum factor density* MaxFW*/*n, *factor weight coverage* FW%, *maximum positive factor density* MaxFW_(+)_*/*n, *positive factor weight coverage* FW_(+)_%, *maximum enhancing factor density* MaxFW_&_*/*n, and *enhancing factor weight coverage* FW_&_% are defined as follows:

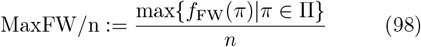

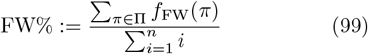

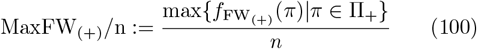

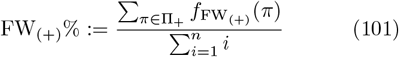

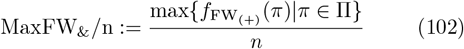

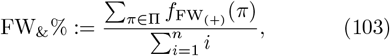

where *f*_FW_ and 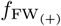 denotes functions returning factor weights or those of positive cascades for a given pathway, respectively.

Finally, we introduce cascade length-related features, *maximum cascade length scale* MaxCL*/*n, *cascade length coverage* CL%, *maximum positive cascade length scale* MaxCL_(+)_*/*n, *positive cascade length coverage* CL_(+)_%, *maximum enhancing cascade length scale* MaxCL_&_*/*n, and *enhancing cascade length coverage* CL_&_%, as follows:

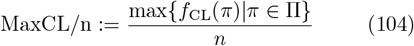

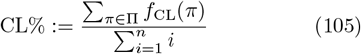

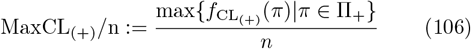

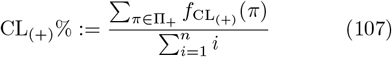

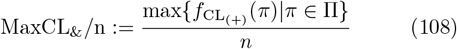

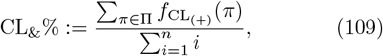

where *f*_CL_ and 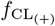 denotes functions returning cascade lengths or those of positive cascades for a given pathway, respectively.

### 4.8 Exhaustive search of network structure with four factors

#### 4.8.1 ESM4: a template for diverse simulated models of different network structures

To explore all possible network structures for certain features that may serve as indicators of the effectiveness of the C+LOO design, we created ESM4, a template model of network structures that can generate various simulated models.

Define the following for all *i, j* ∈ ℕ such that 1 ≤ *i* ≤ 4 and 1 ≤ *j* ≤ *n*_max_:

- *x*_*i,j*_, *y*_*j*_, *c*_*i,j*_, *ε*_*i,j*_, *ε*_*y,j*_, ReLU: identical to those in Models Φ and Ψ
- *K* := {1, 2, 3, 4}
- *L* := {12, 13, 14, 23, 24, 34}
- *a*_*k*_ ∼ Lognormal(2, (0.3)^2^) for *k* ∈ *K*
- *b*_*ℓ*_ ∼ Lognormal(1, (0.5)^2^) for *ℓ* ∈ *L*
- *s*_*λ*_ ∈ {−1, 0, 1} for *λ* ∈ *K* ∪ *L*: edge status indicator (−1, negative; 0, null; and 1, positive)

Notably, *s*_*λ*_ modulates the connectivity patterns and the polarity of factor effects. By varying *s*_*λ*_ values, ESM4 can generate 59,049 different simulated models.

As in the other simulated models, the parameters *v*_*i*_, *a*_*k*_, and *b*_*ℓ*_ are shared across trials and network structures. They were sampled using `NumPy` with a fixed random seed.

The model equations are defined as follows:

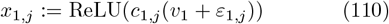

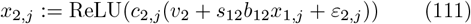

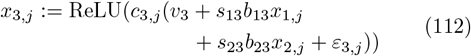

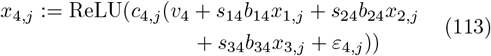

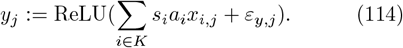

Downstream analyses—namely, simulated experiments and ANOVA—for the models generated with ESM4 were performed in the same way with the other simulated models.

#### 4.8.2 Datasets for the ML model

Using the ESM4 template, we generated 59,049 simulators—all possible network structures for simulators consisted of four factors and their DAGs. After evaluating the C+LOO- and PB-aptitude scores for each simulator at *N* = 3, we classified them into the binary classifications—”C+LOO-suited” and “PB-suited”—referring their aptitude scores for C+LOO 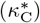 and PB 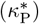 designs. In cases where 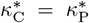, the C+LOO design was defined as the “C+LOO-suited” class, since it requires a smaller total number of trials (Figure 1d and Methods 4.1.3). We also calculated the 31 feature values detailed in Methods 4.7.2 for each simulator.

After the formation of the dataset consisting of the simulator categories and the 31 network features, we performed a random hold-out to split the dataset into the training, validation, and test sets in approximate proportions of 3:1:1.

#### 4.8.3 ML binary classification model

To find plausible features that are associated with high performance of the PB design with a data-driven manner, we created an ML model to predict the simulator classes from the 31 network features. We adopted a GBDT model provided as LightGBM [48]. To optimize the hyper parameters in the LightGBM model, we adopted the `LightGBMTuner`, a LightGBM model employed in `Optuna` [49] that wraps LightGBM training and hyper-parameter optimization.

The performance of the GBDT model was evaluated with an ROC curve and its AUROC value. We implemented the ML model evaluation with `scikit-learn` and other Python packages.

#### 4.8.4 SHAP scores

To understand which feature has the largest contribution to the classification, we computed SHAP scores to visualize the correspondence of feature values to the model output. Since the Python package SHAP [50] can analyze tree-based pretrained models including LightGBM, we passed the GBDT model introduced in Methods 4.8.3 and created a beeswarm plot for the “PB-suited” class showing the top 10 features with the highest impact on the ML model output.

#### 4.8.5 PBSI

To determine which of the C+LOO or PB designs is more suitable for a given phenomenon, it is important to identify the conditions under which the PB design can out-perform the C+LOO design, which is the standard choice for Perturb-seq experiments. Compared to creating ML models based on a comprehensive dataset of network features and experimental design performance for a desired number of factors—as we did with ESM4, having an indicator that predicts high performance with the PB design is an efficient way to streamline experimental planning.

Since single features did not show sufficient prediction performance for the “PB-suited” class (Figure 5e), we developed an integrative metric identifying high performance of the PB design compared to the C+LOO design. We term this metric the PBSI, defined as follows:

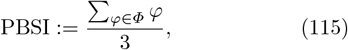

where *Φ* := { P_(+)_%, 1 − MaxRW_(+)_*/*R^∗^, 1 − RW_(+)_% }. Notably, we engineered PBSI by exhaustively searching all possible combinations of the top six individual features in terms of predictive performance—1 − MaxRW_(+)_*/*R^∗^, 1 − RW_(+)_%, 1 − MaxRW_&_*/*R^∗^, P_(+)_%, 1 − 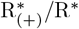, and P%—and selecting the combination that yielded the highest AUROC in the ESM4 dataset (SI I.1).

#### 4.8.6 Prediction performance evaluation for the individual network features and composite indices

As described in Methods 4.7.2, the 31 features were designed to take values within the range [0, 1]. The composite indices constructed by calculating the arithmetic means of feature combinations also shared this property. Using these values directly, we evaluated the ROC curves, AUROC, and average precision (AP) scores to measure the predictive performance of each metric for the “PB-suited” class.

We applied StratifiedKFold from scikit-learn to divide the ESM4 or ESM9 datasets (Methods 4.8.7) into 30 subsets, and calculated ROC curves and AUROC scores for each fold. For the ROC curves, we linearly interpolated the curves across folds using SciPy, accompanied by the raw ROC curves from each fold. For the AUROC values, we calculated the mean and 95% bootstrap CIs across folds using SciPy and visualized them as error bars or annotated them directly in the figures.

Since PBSI was designed as a simple, model-free metric, it was calculated as the arithmetic mean of P_(+)_%, 1 − MaxRW_(+)_*/*R^∗^, and 1 − RW_(+)_%. To examine the potential influence of unoptimized weighting, we built logistic regression models following the same procedure, fitting weights for each component and comparing their predictive performance with that of PBSI (SI I.1–I.2). The logistic regression models were implemented using scikit-learn.

#### 4.8.7 ESM9: another exhaustive search model for nine factors

To evaluate the scalability of the features and metrics associated with relative high performance of PB designs, we created ESM9, another template model that can generate diverse simulators.

Define the following for all *i, j* ∈ ℕ such that 1 ≤ *i* ≤ 9 and 1 ≤ *j* ≤ *n*_max_:

- *x*_*i,j*_, *y*_*j*_, *c*_*i,j*_, *ε*_*i,j*_, *ε*_*y,j*_, ReLU: identical to those in Models Φ and Ψ
- *K* := {*i* | 1 ≤ *i* ≤ 9}
- *L* := {10*i*^∗^ + *i*_∗_ | 1 ≤ *i*^∗^ ≤ 8, *i*^∗^ + 1 ≤ *i*_∗_ ≤ 9}
- *a*_*k*_ ∼ Lognormal(2, (0.3)^2^) for *k* ∈ *K*
- *b*_*ℓ*_ ∼ Lognormal(1, (0.5)^2^) for *ℓ* ∈ *L*
- *s*_*λ*_ ∈ {−1, 0, 1} for *λ* ∈ *K* ∪ *L*: edge status indicator (−1, negative; 0, null; and 1, positive)

Notably, *s*_*λ*_ modulates the connectivity patterns and the polarity of factor effects. By varying *s*_*λ*_ values, ESM9 can generate 2,954,312,706,550,833,698,643 (= 3^45^) different simulators. To reduce computational cost, we randomly sampled 20,000 network configurations and evaluated the C+LOO- and PB-aptitude scores at *N* = 3.

As in the other simulated models, the parameters *v*_*i*_, *a*_*k*_, and *b*_*ℓ*_ are shared across trials and network structures. They were sampled using NumPy with a fixed random seed.

The model equations are defined as follows:

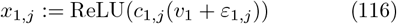

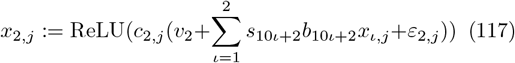

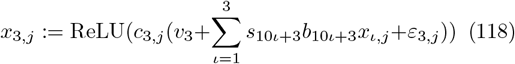

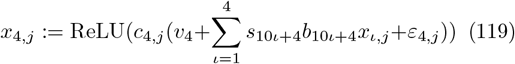

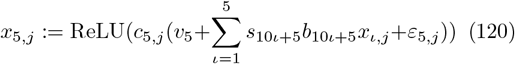

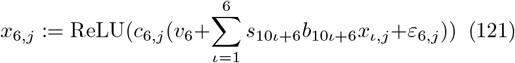

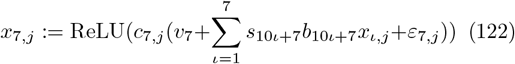

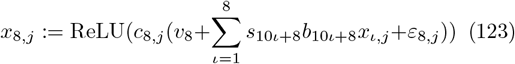

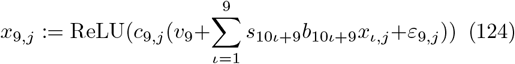

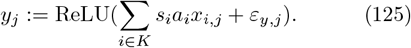

Downstream analyses for the models generated with ESM9 were performed in the same way with the other simulators.

#### 4.8.8 Benchmarking C+LOO and PB designs using PBSI-guided simulators

In SI I.3, we benchmarked the C+LOO and PB designs using newly introduced simulators, Models Π, Δ, and Σ. The network configurations for these models within the ESM9 framework were explicitly engineered based on PBSI logic to verify whether the relative performance of the two designs could be predicted and controlled by the PBSI score.

Model Π was constructed to represent a high-PBSI (“PB-suited”) architecture characterized by pure parallelism. In this model, all direct pathways were enabled while all inter-factor regulations were disabled. Formally, the edge weights were assigned as *s*_*k*_ = 1 for pathway edges *k* ∈ *K* and *s*_*ℓ*_ = 0 for regulatory edges *ℓ* ∈ *L*, ensuring distinct, independent signal transmission for each factor.

In contrast, Model Δ was constructed to represent a low-PBSI (“C+LOO-suited”) architecture driven by signal amplification. The edge weights were assigned as follows:

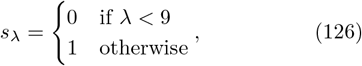

where index *λ* ∈ *K* ∪ *L* represents the ordered edges in the ESM9 topology. This configuration results in a structure with a single positive pathway at the most downstream level, coupled with the full set of positive inter-factor regulations permitted in ESM9. This topology forces all upstream signals to merge and amplify through a dense regulatory network before converging on a single output node, thereby minimizing P_(+)_% and ensuring large values of MaxRW_(+)_*/*R^∗^ and RW_(+)_%.

Model Σ serves as a critical control: a system that is topologically entangled (like Model Δ) but yields a high PBSI score (like Model Π). Modifying the Model Δ architecture, we assigned edges as follows:

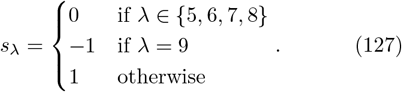

This configuration introduces two key alterations. First, the terminal pathway edge (*λ* = 9) is set to − 1, inverting the signal amplification generated by the positive regulations. Second, additional positive pathways (indices *λ* ∈ {1, 2, 3, 4}) are enabled. These extra parallel paths elevate the numerator term P_(+)_%, while the low covariance maintained by the dampened interactions ensures that the overall PBSI score remains high despite the presence of complex crosstalk.

Using the same benchmarking protocols applied to Models Φ, Ψ, and Λ, we evaluated the performance of the C+LOO and PB designs with these simulators.

### 4.9 scRNA-seq data analysis

To validate the biological interpretation of PBSI— specifically, that parallel gene modules exhibit high PBSI while signal amplification networks exhibit low PBSI— we analyzed the Perturb-seq dataset of K562 cells published by Replogle et al. (2022) [18]. The raw single-cell data (`K562 essential raw singlecell 01.h5ad`) were downloaded, and only the non-targeting (control) condition was utilized for this study. Our analysis aimed to construct GRNs corresponding to various biological phenomena and examine the relationship between their network topologies and PBSI scores. Implementation details are available in the GitHub repository described in Data and Code Availability 5.

#### 4.9.1 Data preprocessing

Data preprocessing was performed using `Scanpy` [51]. Quality control metrics, including total counts, number of genes by counts, and the percentage of mitochondrial genes, were visualized to confirm that the deposited data had been appropriately preprocessed. Doublet removal was executed using the `Scanpy` interface for `Scrublet` [52]. Subsequently, total count normalization (scaling to 10,000 counts per cell), log-transformation, highly variable gene selection, Principal Component Analysis, construction of the nearest neighbor graph, and dimensionality reduction via Uniform Manifold Approximation and Projection (UMAP) [53] were performed using standard `Scanpy` workflows. To facilitate robust variable selection and causal discovery, we aggregated the cells into MetaCells using the `SEACells` algorithm [54], resulting in the assignment of 213 MetaCells. These MetaCells were used as pseudobulk samples for the subsequent GRN inference.

#### 4.9.2 Phenotype scoring

For the S-phase score and other phenotype scores, gene sets were retrieved from the MSigDB Hallmark 2020 collection [55] via `GSEApy` [56]. Genes included in the dataset were extracted from the sub-categories, accounting for gene symbol aliases using `MyGene` [57]. Scoring was performed using the `score genes` function implemented in `Scanpy`. The raw scores were standardized to Z-scores and subsequently transformed using a sigmoid function to rescale the values to the range [0, 1]. Specifically for the S-phase score, we calculated the score using genes from the “E2F Targets” category, explicitly excluding those that overlap with the “G2-M Checkpoint” category to ensure phase specificity.

#### 4.9.3 Feature selection for GRNs

We determined the target configuration for each GRN— either parallel or entangled—based on the visualization of Pearson correlation heatmaps for gene sets within the sub-categories of the MSigDB Hallmark 2020 collection. Theoretically, GRN inference aims to construct graphs representing causal regulatory structures. However, given that statistical dependencies and correlation coefficients are often employed as surrogate measures in the context of causal discovery [58], we selected gene sets with low mutual correlations for parallel GRNs and those with high correlations for entangled GRNs.

To identify gene sets for parallel GRNs, we selected the top 20 genes with low mutual correlation by performing Recursive Feature Elimination using LASSO regression as the estimator, with the phenotypic score of each sub-category as the target variable. Conversely, for entangled GRNs, we selected the top 20 genes exhibiting the highest individual correlation with the phenotypic score of the respective sub-category. All implementations were conducted using standard Python packages, including `scikit-learn` and `Pandas`.

#### 4.9.4 Bayesian network inference and PBSI evaluation

To simulate a scenario where a phenotypic score serves as the primary readout—specifically, the impact of individual gene knockdowns on the S-phase score—we performed structural inference of GRNs using the 20 candidate genes selected for each sub-category and the S-phase score. We employed a Bayesian network-based causal discovery framework, which learns caulaisty from data structure, rather than specific biological domain knowledge (e.g., transcription factor binding), allowing for flexible feature selection and the inclusion of non-genetic variables such as phenotypic scores.

The single-cell data were standardized using the `StandardScaler` from `scikit-learn` and subsequently averaged per MetaCell. Using these aggregated data, we estimated the DAG structures by maximizing the Bayesian Information Criterion through a Hill-Climb search implemented in `Pgmpy` [59]. To ensure the S-phase score was positioned as the terminal node, we identified all genes upstream of the S-phase score and extracted the corresponding induced subgraphs using `NetworkX` [60]. The regulatory signs (upregulation or downregulation) for each edge were assigned based on the signs of the Pearson correlation coefficients between each gene and the S-phase score. This process resulted in a signed DAG where the output variable is located at the lowest downstream level, consistent with our simulation framework.

Finally, we evaluated P_(+)_%, 1 − MaxRW_(+)_*/*R^∗^, 1 − RW_(+)_%, and PBSI for the inferred GRNs. Since the number of factors in these GRNs varied depending on the identified upstream components, we implemented a dynamic programming-based algorithm using NumPy to efficiently compute these topology metrics for varying graph sizes.

### 4.10 AI usage declaration

In this work, we used large language models, including ChatGPT and Gemini, for code review and language proof editing. All code and text were initially drafted by YO. Following proofreading by the AI models aimed at bug detection and readability improvement, all suggestions were scrutinized, verified, and modified if needed by YO. The manuscript drafted by YO was further reviewed by all other authors.

No AI models were used to create figures, including conceptual diagrams—namely, Figure 2 and Figures S3– S4—which were created by YO using Adobe Illustrator.

## 5 Data and Code Availability

The scRNA-seq data analyzed in this study were obtained from the Figshare repository associated with Replogle et al. (2022) [18] (https://plus.figshare.com/articles/dataset/Mapping_information-rich_genotype-phenotype_landscapes with genome-scale Perturb-seq Replogle et al 2022 processed Perturb-seq datasets/20029387). All analysis code is available in our GitHub repository (https://github.com/yo-aka-gene/WhyDOE).

## 6 Author information

YO conceived the method and performed the experiments with input from TI and KS. YO assessed the statistical validity of the method under supervision from YS. YO developed biological interpretability of the method with input from YS, HO, and KS. YO wrote the manuscript with input from TI, YS, HO, and KS. All the authors reviewed the manuscript.

## 7 Acknowledgements

TI has received a grant from Japan Society for the Promotion of Science (JSPS) KAKENHI Grant Number JP25K21344.

We would like to express our sincere gratitude to Drs. Hikaru Sugimoto (The University of Tokyo), Herbert Yao (University of British Columbia), Profs. Shinya Kuroda (The University of Tokyo), and Nozomu Yachie (University of British Columbia) for their critical reading and their valuable suggestions.

## Abbreviations

ANOVA: analysis of variance
AP: average precision
AUROC: area under the ROC
C+LOO: Control-and-Leave-One-Out
CI: confidence interval
DAG: directed acyclic graph
DO-C+LOO: D-optimized C+LOO
DO-PB: D-optimized PB
DOE: design of experiments
ESM4: Exhaustive Search Model of four factors
ESM9: Exhaustive Search Model of nine factors
FF: full factorial
GBDT: gradient boosting decision tree
GRN: gene regulatory network
KD: knockdown
ML: machine learning
MLR: multivariate linear regression
OFAT: One-Factor-At-a-Time
PB: Plackett–Burman
PBSI: PB suitability index
ReLU: rectified linear unit
ROC: receiver operating characteristic
scRNA-seq: single-cell RNA-sequencing
SHAP: Shapley additive explanations
SI: Supplementary Information
TPL: Taylor’s power law
UMAP: Uniform Manifold Approximation and Projection
VIF: variance inflation factor

