## Supplemental Information for "Design-of-Experiments for Nonlinear, Multivariate Biology: Rethinking Experimental Design through Perturb-seq"

**A Mathematical notation** **ii**
A.1 Basic notation . . . ii
A.2 Notation for the C+LOO design and related statistics . . . ii
A.3 Interaction terms . . . iii

**B Correlation coefficients of the C+LOO design** **iii**
B.1 Correlation coefficients between the main effects . . . iii
B.2 Correlations between main effects and interaction terms . . . iv
B.3 Correlation coefficients between the interaction terms . . . v

**C Experimental designs from DOE** **vi**
C.1 The FF design and experimental costs . . . vi
C.2 An overview of the PB design . . . vii

**D Dissecting Perturb-seq capabilities and limitations via a minimalist simulator** **viii**
D.1 Formalization of Perturb-seq and the temporal dynamics of GRN structures . . . viii
D.2 Rationale for a minimalist simulator . . . xi
D.3 Taylor's power law (TPL) suggests our simulators model "canonical behavior" of biological systems . xii

**E A generic workflow of C+LOO-based Perturb-seq vs. DOE-originated multivariate analysis** **xiii**

**F Impact of the inherent correlations: Interaction Leakage in the C+LOO design** **xiii**

**G Correspondence between simulator configurations and experimental design performance** **xvi**
G.1 Model $\Phi$: a basic model simulating signaling cascades . . . xvi
G.2 Model $\Psi$: a highly complex regulatory circuit of interactions . . . xvii
G.3 Model $\Lambda$: a model with poorly curated candidates . . . xvii
G.4 C+LOO-based multivariate models vs. Dunnett's test models . . . xvii

**H Investigating the role of C+LOO-inherent correlations via D-optimization** **xvii**

**I PBSI** **xxiv**
I.1 Identification of PBSI . . . xxiv
I.2 Scalability of PBSI . . . xxiv
I.3 Explainability of PBSI: validation via representative topological features . . . xxiv
I.4 Biological relevance of PBSI-related GRN topologies . . . xxxiii

**J Practical guidelines for experimental design in multivariate biology** **xli**
J.1 Selection of candidate genes . . . xli
J.2 Experimental design choice: C+LOO vs. PB . . . xlii
J.3 Strategies in the absence of preliminary scRNA-seq data . . . xliii
J.4 Analytical framework: Shifting from pairwise comparisons to MLR/ANOVA . . . xliv
J.5 Result interpretation: on translating readouts from statistical analysis into biological language . . . xliv
J.6 Perspectives for detecting combinatorial effects . . . xlv

### A Mathematical notation

#### A.1 Basic notation

To facilitate the discussion of the C+LOO design, we first introduce basic notation. First, we introduce the following notation:

$$\underline{x} := \{\alpha \in \mathbb{N} \mid 1 \leq \alpha \leq x\}. \tag{S-1}$$

Then, we denote an $n$-dimensional vector ($\forall n \in \mathbb{N}$) $\mathbf{x} \in \mathbb{R}^n$ using Equation (S-1) as follows:

$$\mathbf{x} := \begin{bmatrix} x_1 \\ x_2 \\ \vdots \\ x_n \end{bmatrix} =: (x_i)_{i \in \underline{n}} \tag{S-2}$$

where $\forall i \in \underline{n}$, $x_i \in \mathbb{R}$. Likewise, we introduce $\mathbf{1}_n$ as follows:

$$\mathbf{1}_n := (1)_{i \in \underline{n}} = \begin{bmatrix} 1 \\ 1 \\ \vdots \\ 1 \end{bmatrix}. \tag{S-3}$$

Next, we denote the Hadamard multiplication $\odot$ as follows (let $\forall \mathbf{x}, \mathbf{y} \in \mathbb{R}^n$ such that $\mathbf{x} = (x_i)_{i \in \underline{n}}$ and $\mathbf{y} = (y_i)_{i \in \underline{n}}$):

$$\mathbf{x} \odot \mathbf{y} := (x_i \cdot y_i)_{i \in \underline{n}} = (y_i \cdot x_i)_{i \in \underline{n}} = \mathbf{y} \odot \mathbf{x}. \tag{S-4}$$

Note that $\mathbf{x} \odot \mathbf{1}_n = \mathbf{x}$. In addition to these linear-algebraic notations, the Kronecker delta $\delta_{i,j}$ is given as follows:

$$\delta_{i,j} = \begin{cases} 1 & (i = j) \\ 0 & (i \neq j) \end{cases}. \tag{S-5}$$

Finally, we define the expected value $\mathrm{E}[\mathbf{x}]$, variance $\mathrm{Var}[\mathbf{x}]$, and correlation coefficient $\mathrm{Corr}(\mathbf{x}, \mathbf{y})$ of the random variables $\mathbf{x}, \mathbf{y} \in \mathbb{R}^n$ as follows:

$$\mathrm{E}[\mathbf{x}] := \frac{\sum_{i \in \underline{n}} x_i}{n} \tag{S-6}$$

$$\mathrm{Var}[\mathbf{x}] := \mathrm{E}[(\mathbf{x} - \mathrm{E}[\mathbf{x}] \cdot \mathbf{1}_n) \odot (\mathbf{x} - \mathrm{E}[\mathbf{x}] \cdot \mathbf{1}_n)] = \mathrm{E}[\mathbf{x} \odot \mathbf{x}] - \{\mathrm{E}[\mathbf{x}]\}^2 \tag{S-7}$$

$$\mathrm{Corr}(\mathbf{x}, \mathbf{y}) := \frac{\mathrm{E}[(\mathbf{x} - \mathrm{E}[\mathbf{x}] \cdot \mathbf{1}_n) \odot (\mathbf{y} - \mathrm{E}[\mathbf{y}] \cdot \mathbf{1}_n)]}{\sqrt{\mathrm{Var}[\mathbf{x}] \cdot \mathrm{Var}[\mathbf{y}]}} = \frac{\mathrm{E}[((x_i - \mathrm{E}[\mathbf{x}]) \cdot (y_i - \mathrm{E}[\mathbf{y}]))_{i \in \underline{n}}]}{\sqrt{\mathrm{Var}[\mathbf{x}] \cdot \mathrm{Var}[\mathbf{y}]}}. \tag{S-8}$$

#### A.2 Notation for the C+LOO design and related statistics

The $i$-th column vector $\mathbf{x}_i$ of $C(n)$ design matrix ($\forall n \in \mathbb{N}$, $\forall i \in \underline{n}$) corresponding to $i$-th factor is denoted as follows:

$$\mathbf{x}_i := (1 - 2\delta_{i+1,j})_{j \in \underline{n+1}}. \tag{S-9}$$

Note that the elements of $\mathbf{x}_i$ are conditioned as follows:

$$1 - 2\delta_{i+1,j} = \begin{cases} -1 & (i + 1 = j) \\ 1 & (i + 1 \neq j) \end{cases}. \tag{S-10}$$

Accordingly, the expected value $\mathrm{E}[\mathbf{x}_i]$ and the variance $\mathrm{Var}[\mathbf{x}_i]$ of $\mathbf{x}_i$ are given as follows:

$$\mathrm{E}[\mathbf{x}_i] = \frac{n-1}{n+1} \tag{S-11}$$

$$\mathrm{Var}[\mathbf{x}_i] = \mathrm{E}[\mathbf{x}_i \odot \mathbf{x}_i] - \{\mathrm{E}[\mathbf{x}_i]\}^2 = 1 - (\frac{n-1}{n+1})^2 = \frac{4n}{(n+1)^2}. \tag{S-12}$$

##### A.3 Interaction terms

When $n \geq 2$ and $\forall i, j \in \underline{n}$ such that $i \neq j$, we define the interaction term $\mathbf{x}_{i\times j}$ of $\mathbf{x}_i$ and $\mathbf{x}_j$ as follows:

$$\mathbf{x}_{i\times j} := \mathbf{x}_i \odot \mathbf{x}_j = (1 - 2(\delta_{i+1,k} + \delta_{j+1,k}))_{k\in\underline{n+1}}. \tag{S-13}$$

The elements of $\mathbf{x}_{i\times j}$ are conditioned as follows:

$$1 - 2(\delta_{i+1,k} + \delta_{j+1,k}) \;=\; \begin{cases} -1 & (i+1 = k \;\vee\; j+1 = k) \\ 1 & (\text{otherwise}) \end{cases}. \tag{S-14}$$

Note that the interaction terms are symmetrical (i.e., $\forall i, j \in \underline{n}$, $\mathbf{x}_{i\times j} = \mathbf{x}_i \odot \mathbf{x}_j = \mathbf{x}_j \odot \mathbf{x}_i = \mathbf{x}_{j\times i}$). Furthermore, the expected value $\mathrm{E}[\mathbf{x}_{i\times j}]$ and the variance $\mathrm{Var}[\mathbf{x}_{i\times j}]$ of $\mathbf{x}_{i\times j}$ are introduced as follows:

$$\mathrm{E}[\mathbf{x}_{i\times j}] = \frac{n-3}{n+1} \tag{S-15}$$

$$\mathrm{Var}[\mathbf{x}_{i\times j}] = \mathrm{E}[\mathbf{x}_{i\times j} \odot \mathbf{x}_{i\times j}] - \{\mathrm{E}[\mathbf{x}_{i\times j}]\}^2 = 1 - (\frac{n-3}{n+1})^2 = \frac{8(n-1)}{(n+1)^2}. \tag{S-16}$$

#### B Correlation coefficients of the C+LOO design

##### B.1 Correlation coefficients between the main effects

$\forall i, j \in \underline{n}$, we define the correlation coefficient $\rho_{i,j}$ of $\mathbf{x}_i$ and $\mathbf{x}_j$ as follows (this notation is introduced for short):

$$\rho_{i,j} := \mathrm{Corr}(\mathbf{x}_i, \mathbf{x}_j) = \frac{(n+1)^2}{4n} \cdot \mathrm{E}[((\frac{2}{n+1} - 2\delta_{i+1,k}) \cdot (\frac{2}{n+1} - 2\delta_{j+1,k}))_{k\in\underline{n+1}}]. \tag{S-17}$$

###### B.1.1 When $i = j$

When $i = j$, the correlation coefficient $\rho_{i,i}$ is given as follows:

$$\rho_{i,i} = \frac{(n+1)^2}{4n} \cdot \mathrm{E}[((\frac{2}{n+1} - 2\delta_{i+1,k})^2)_{k\in\underline{n+1}}]. \tag{S-18}$$

Note that $(\frac{2}{n+1} - 2\delta_{i+1,k})^2$ is conditioned as follows:

$$(\frac{2}{n+1} - 2\delta_{i+1,k})^2 \;=\; \begin{cases} \frac{4n^2}{(n+1)^2} & (i+1 = k) \\ \frac{4}{(n+1)^2} & (i+1 \neq k) \end{cases}. \tag{S-19}$$

Therefore, $\mathrm{E}[((\frac{2}{n+1} - 2\delta_{i+1,k})^2)_{k\in\underline{n+1}}]$ in Equation (S-18) and $\rho_{i,i}$ are given as follows:

$$\mathrm{E}[((\frac{2}{n+1} - 2\delta_{i+1,k})^2)_{k\in\underline{n+1}}] = \frac{4}{(n+1)^2} \cdot \frac{n\cdot 1 + n^2}{n+1} = \frac{4n}{(n+1)^2} \tag{S-20}$$

$$\rho_{i,i} = \frac{(n+1)^2}{4n} \cdot \frac{4n}{(n+1)^2} = 1. \tag{S-21}$$

Although $\mathrm{Corr}(\mathbf{x}_i, \mathbf{x}_i) = 1$ by definition, we aimed to demonstrate the calculation process for reference, and check if the notation of the correlation coefficients introduced in Equation (S-17) is well-defined.

###### B.1.2 When $i \neq j$

If $n \geq 2$, there exists a pair of numbers $i, j \in \underline{n}$ such that $i \neq j$. Considering such conditions, the following equations hold:

$$(\frac{2}{n+1} - 2\delta_{i+1,k}) \cdot (\frac{2}{n+1} - 2\delta_{j+1,k}) \;=\; \begin{cases} \frac{-4n}{(n+1)^2} & (i+1 = k \;\vee\; j+1 = k) \\ \frac{4}{(n+1)^2} & (\text{otherwise}) \end{cases} \tag{S-22}$$

$$\mathrm{E}[((\frac{2}{n+1} - 2\delta_{i+1,k}) \cdot (\frac{2}{n+1} - 2\delta_{j+1,k}))_{k\in\underline{n+1}}] = \frac{4}{(n+1)^2} \cdot \frac{(n-1)\cdot 1 + 2\cdot(-n)}{n+1} = -\frac{4}{(n+1)^2}. \tag{S-23}$$

Given Equations (S-17) and (S-23), $\rho_{i,j}$ is introduced as follows:

$$\rho_{i,j} = \frac{(n+1)^2}{4n} \cdot \frac{-4}{(n+1)^2} = -\frac{1}{n}. \tag{S-24}$$

#### B.2 Correlations between main effects and interaction terms

Extending Equation (S-17), we define the correlation coefficient $\rho_{i\times j,k}$ between $\mathbf{x}_{i\times j}$ and $\mathbf{x}_k$ ($n \geq 2$, $\forall i, j, k \in \underline{n}$) as follows:

$$\rho_{i\times j,k} := \mathrm{Corr}(\mathbf{x}_{i\times j}, \mathbf{x}_k) = \frac{(n+1)^2}{4\sqrt{2n(n-1)}} \cdot \mathrm{E}[(\{\frac{4}{n+1} - 2(\delta_{i+1,l} + \delta_{j+1,l})\} \cdot (\frac{2}{n+1} - 2\delta_{k+1,l}))_{l\in\underline{n+1}}]. \tag{S-25}$$

It is noteworthy that $\rho_{k,i\times j} := \mathrm{Corr}(\mathbf{x}_k, \mathbf{x}_{i\times j}) = \rho_{i\times j,k}$ because of the symmetry of the correlation function Equation (S-8). Therefore, we investigate only the property of $\rho_{i\times j,k}$, and the same results apply for $\rho_{k,i\times j}$ as well as $\rho_{j\times i,k}$ and $\rho_{k,j\times i}$.

##### B.2.1 When $i = k$ or $j = k$

First, we consider a correlation coefficient of the main effect of a factor (e.g., $\mathbf{x}_i$) and its associated interaction term (e.g., $\mathbf{x}_{i\times j}$). Given the symmetry of interaction terms, we hypothesize that $i = k$ (so we consider $\rho_{i\times j,i}$) and the following equations are introduced:

$$\rho_{i\times j,i} = \frac{(n+1)^2}{4\sqrt{2n(n-1)}} \cdot \mathrm{E}[(\{\frac{4}{n+1} - 2(\delta_{i+1,l} + \delta_{j+1,l})\} \cdot (\frac{2}{n+1} - 2\delta_{i+1,l}))_{l\in\underline{n+1}}] \tag{S-26}$$

$$\{\frac{4}{n+1} - 2(\delta_{i+1,l} + \delta_{j+1,l})\} \cdot (\frac{2}{n+1} - 2\delta_{i+1,l}) = \begin{cases} \frac{4n(n-1)}{(n+1)^2} & (i+1 = l) \\ \frac{-4(n-1)}{(n+1)^2} & (j+1 = l) \\ \frac{8}{(n+1)^2} & (\text{otherwise}) \end{cases} \tag{S-27}$$

$$\mathrm{E}[(\{\frac{4}{n+1} - 2(\delta_{i+1,l} + \delta_{j+1,l})\} \cdot (\frac{2}{n+1} - 2\delta_{i+1,l}))_{l\in\underline{n+1}}] = \frac{4(n-1)}{(n+1)^2}. \tag{S-28}$$

Given Equations (S-26) and (S-28), $\rho_{i\times j,i}$ is formulated as follows:

$$\rho_{i\times j,i} = \frac{(n+1)^2}{4\sqrt{2n(n-1)}} \cdot \frac{4(n-1)}{(n+1)^2} = \sqrt{\frac{n-1}{2n}}. \tag{S-29}$$

The same results apply for $\rho_{j\times i,i}$, $\rho_{i,i\times j}$, and $\rho_{i,j\times i}$.

##### B.2.2 When $i \neq k$ and $j \neq k$

If $n \geq 3$, there exists a triplet of numbers $i, j, k \in \underline{n}$ such that $i \neq j$, $i \neq k$ and $j \neq k$. Under such a condition, the following equations hold:

$$\{\frac{4}{n+1} - 2(\delta_{i+1,l} + \delta_{j+1,l})\} \cdot (\frac{2}{n+1} - 2\delta_{k+1,l}) = \begin{cases} \frac{-4(n-1)}{(n+1)^2} & (i+1 = l \ \vee \ j+1 = l) \\ \frac{-8n}{(n+1)^2} & (k+1 = l) \\ \frac{8}{(n+1)^2} & (\text{otherwise}) \end{cases} \tag{S-30}$$

$$\mathrm{E}[(\{\frac{4}{n+1} - 2(\delta_{i+1,l} + \delta_{j+1,l})\} \cdot (\frac{2}{n+1} - 2\delta_{k+1,l}))_{l\in\underline{n+1}}] = \frac{-8}{(n+1)^2}. \tag{S-31}$$

Therefore, $\rho_{i\times j,k}$ is given as follows:

$$\rho_{i\times j,k} = \frac{(n+1)^2}{4\sqrt{2n(n-1)}} \cdot \frac{-8}{(n+1)^2} = -\sqrt{\frac{2}{n(n-1)}}. \tag{S-32}$$

The same results apply for $\rho_{j\times i,k}$, $\rho_{k,i\times j}$ and $\rho_{k,j\times i}$.

##### B.3 Correlation coefficients between the interaction terms

Finally, we consider correlation coefficients among interaction terms. $\forall n \geq 2$ and $\forall i, j, k, l \in \underline{n}$ such that $i \neq j$ and $l \neq k$, we define the correlation coefficient $\rho_{i\times j, k\times l}$ of interaction terms $\mathbf{x}_{i\times j}$ and $\mathbf{x}_{k\times l}$ as follows:

$$\rho_{i\times j,k\times l} := \mathrm{Corr}(\mathbf{x}_{i\times j}, \mathbf{x}_{k\times l}) = \frac{(n+1)^2}{8(n-1)} \cdot \mathrm{E}[(\{\frac{4}{n+1} - 2(\delta_{i+1,m} + \delta_{j+1,m})\} \cdot \{\frac{4}{n+1} - 2(\delta_{k+1,m} + \delta_{l+1,m})\})_{m \in \underline{n+1}}] \tag{S-33}$$

In order to avoid redundancy, here we introduce a notation $\Xi_x^c(a,b)$ (where $\forall x \in \mathbb{N}$, $\forall a, b, c \in \underline{x}$ such that $a \neq b$) as follows:

$$\Xi_x^c(a,b) := \frac{4}{x} - 2(\delta_{a,c} + \delta_{b,c}). \tag{S-34}$$

Therefore, using Equation (S-34), Equation (S-33) can be formulated as follows:

$$\rho_{i\times j,k\times l} = \frac{(n+1)^2}{8(n-1)} \cdot \mathrm{E}[(\Xi_{n+1}^m(i+1, j+1) \cdot \Xi_{n+1}^m(k+1, l+1))_{m \in \underline{n+1}}]. \tag{S-35}$$

###### B.3.1 When $\mathbf{x}_{i\times j} = \mathbf{x}_{k\times l}$

When $\mathbf{x}_{i\times j} = \mathbf{x}_{k\times l}$, equations regarding the correlation coefficient $\rho_{i\times j,i\times j}$ are given as follows:

$$\rho_{i\times j,i\times j} = \frac{(n+1)^2}{8(n-1)} \cdot \mathrm{E}[(\{\Xi_{n+1}^m(i+1, j+1)\}^2)_{m \in \underline{n+1}}] \tag{S-36}$$

$$\{\Xi_{n+1}^m(i+1,j+1)\}^2 = \begin{cases} \frac{4(n-1)^2}{(n+1)^2} & (i+1 = m \ \vee \ j+1 = m) \\ \frac{16}{(n+1)^2} & (\text{otherwise}) \end{cases} \tag{S-37}$$

$$\mathrm{E}[(\{\Xi_{n+1}^m(i+1, j+1)\}^2)_{m \in \underline{n+1}}] = \frac{8(n-1)}{(n+1)^2}. \tag{S-38}$$

Accordingly, the following equation holds:

$$\rho_{i\times j,i\times j} = \frac{(n+1)^2}{8(n-1)} \cdot \frac{8(n-1)}{(n+1)^2} = 1. \tag{S-39}$$

Again, it is trivial that $\rho_{i\times j,i\times j} = \mathrm{Corr}(\mathbf{x}_{i\times j}, \mathbf{x}_{i\times j}) = 1$. Yet, we could confirm that Equation (S-35) is well-defined with Equations (S-36)–(S-39).

###### B.3.2 When one factor is shared between the two interaction terms (e.g., $\mathbf{x}_{i\times j}$ and $\mathbf{x}_{i\times l}$)

Next, we consider situations where the two interaction terms share one factor. In other words, the following conditions meet the criterion ($n \geq 3$ and $\forall i, j, k, l \in \underline{n}$ such that $i \neq j$ and $k \neq l$):

1. $i = k$ and $j \neq l$
2. $i = l$ and $j \neq k$
3. $j = k$ and $i \neq l$
4. $j = l$ and $i \neq k$.

As the same results will apply for each condition, we demonstrate the first case and consider the correlation coefficient $\rho_{i\times j,i\times l}$ for $\mathbf{x}_{i\times j}$ and $\mathbf{x}_{i\times l}$. Accordingly, the following equations are introduced:

$$\rho_{i\times j,i\times l} = \frac{(n+1)^2}{8(n-1)} \cdot \mathrm{E}[(\Xi_{n+1}^m(i+1, j+1) \cdot \Xi_{n+1}^m(i+1, l+1))_{m \in \underline{n+1}}] \tag{S-40}$$

$$\Xi_{n+1}^m(i+1,j+1) \cdot \Xi_{n+1}^m(i+1,l+1) = \begin{cases} \frac{4(n-1)^2}{(n+1)^2} & (i+1 = m) \\ \frac{-8(n-1)}{(n+1)^2} & (j+1 = m \ \vee \ l+1 = m) \\ \frac{16}{(n+1)^2} & (\text{otherwise}) \end{cases} \tag{S-41}$$

$$\mathrm{E}[(\Xi_{n+1}^m(i+1,j+1)\cdot\Xi_{n+1}^m(i+1,l+1))_{m\in\underline{n+1}}]=\frac{4(n-3)}{(n+1)^2}. \tag{S-42}$$

According to those equations, $\rho_{i\times j,i\times l}$ is given as follows:

$$\rho_{i\times j,i\times l}=\frac{(n+1)^2}{8(n-1)}\cdot\frac{4(n-3)}{(n+1)^2}=\frac{n-3}{2(n-1)}. \tag{S-43}$$

##### B.3.3 When none of $i,j,k,l$ are equal

When $n\geq 4$, there exists a quadruple of numbers $i,j,k,l\in\underline{n}$ such that $(i-j)(i-k)(i-l)(j-k)(j-l)(k-l)\neq 0$. In other words, $i$, $j$, $k$, and $l$ are distinct numbers, with no two being the same. Then, the following equations hold:

$$\Xi_{n+1}^m(i+1,j+1)\cdot\Xi_{n+1}^m(k+1,l+1)\ =\ \begin{cases}\frac{-8(n-1)}{(n+1)^2} & (i+1=m\ \vee\ j+1=m\ \vee\ k+1=m\ \vee\ l+1=m)\\ \frac{16}{(n+1)^2} & (\text{otherwise})\end{cases} \tag{S-44}$$

$$\mathrm{E}[(\Xi_{n+1}^m(i+1,j+1)\cdot\Xi_{n+1}^m(k+1,l+1))_{m\in\underline{n+1}}]=-\frac{16}{(n+1)^2}. \tag{S-45}$$

Hence, $\rho_{i\times j,k\times l}$ in such a case is given as follows:

$$\rho_{i\times j,k\times l}=\frac{(n+1)^2}{8(n-1)}\cdot\frac{-16}{(n+1)^2}=-\frac{2}{n-1}. \tag{S-46}$$

### C Experimental designs from DOE

#### C.1 The FF design and experimental costs

FF design runs all possible conditions across $2^n$ trials for $n$ factors (Figure S1). For statistical models involving multiple variables (and their interaction terms), multivariate analysis methods such as MLR and ANOVA—both standard approaches in DOE—are well-suited.

However, this design is often impractical, especially when the number of candidate genes is large. For instance, using the FF design for identifying pluripotency-inducing factors from 24 candidate genes—an experiment performed using the C+LOO design in 2006 [TY06]—would require 16,777,216 conditions. This would necessitate an equal number of distinct sgRNA cocktails if performed with Perturb-seq. Beyond the experimental burden, the computational cost for scRNA-seq data analysis would also escalate to an astronomical level.

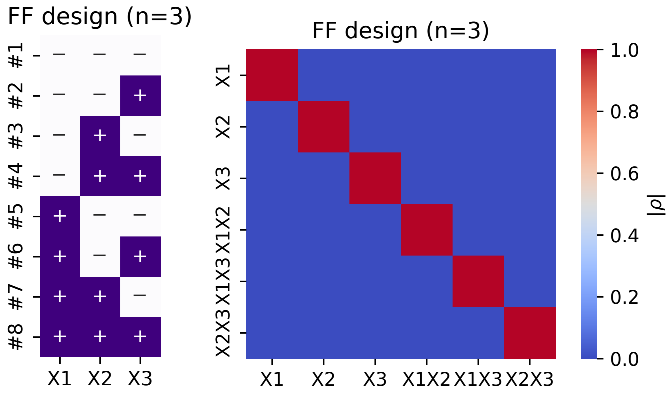


(a) Design matrix (b) Correlation heatmap

**Figure S1: The FF design**

**a–b**: (a) Design matrix and (b) correlation heatmap of the FF designs for 3 factors. The signs in the design matrix indicate inclusion or exclusion of factors. Notably, it is a unique characteristic of the FF design to yield zero correlations for all elements other than $\rho_{i,i}$.

#### C.2 An overview of the PB design

In terms of the balance between utility in experiments and cost-efficiency, the PB design (Figure S2a) serves a popular alternative as a class of screening design. This Hadamard-matrix-based design results in MLR models with lower D-criterion values compared to C+LOO-based MLR models (Figure S2b), indicating that model coefficients are more independent of each other [dABK+95]. Thus, the PB design is well-suited for screening experiments (in terms of DOE) because the factors and their associated interaction terms show no collinearity (i.e., $\rho_{i,j} = 0$, $\rho_{i \times j, i} = 0$, and $\rho_{i \times j, i \times l} = 0$) with a relatively small number of trials (Figure S2c). As a result, the experimenters can reduce the number of candidate factors by focusing on $\mathbf{x}_i$ without being misled by $\mathbf{x}_{i \times j}$. However, for certain purposes, further investigation using experimental designs with higher resolutions is required during the optimization process, as $\rho_{i, j \times k} \neq 0$ and $\rho_{i \times j, k \times l} \neq 0$.

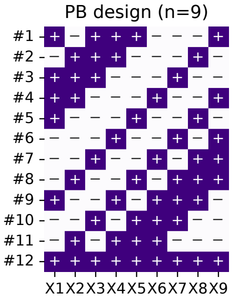


(a) Design matrix

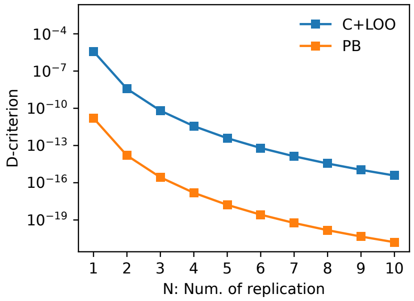


(b) D-criterion comparison

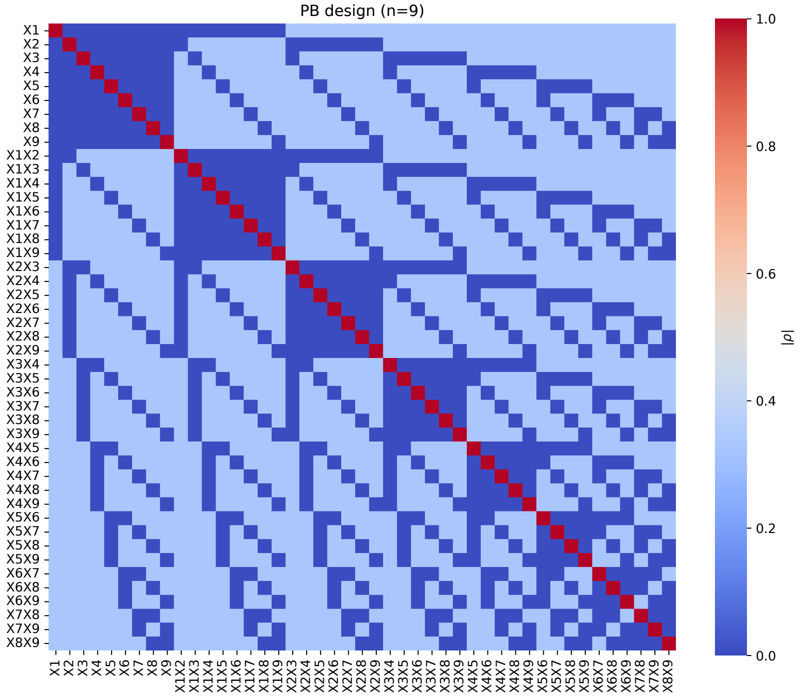


(c) Correlation heatmap

**Figure S2: The PB design matrix and its correlation heatmap**

**a**: The design matrix of the PB design for nine factors (+, addition of factors; −, subtraction of factors). **b**: The D-criterion values of the additive model matrices based on the C+LOO design (blue) and PB design (orange) for nine factors at different numbers of replication ($1 \leq N \leq 10$). **c**: The heatmap of the absolute values of the correlation coefficients (denoted as $|\rho|$) among main effects and interaction terms of factors when they are investigated under the design matrix above.

### D Dissecting Perturb-seq capabilities and limitations via a minimalist simulator

#### D.1 Formalization of Perturb-seq and the temporal dynamics of GRN structures

To assess the impact of experimental design on the interpretation of Perturb-seq results, it is necessary to first formulate what Perturb-seq targets and the scheme by which it elucidates biological phenomena. In other words, to fundamentally understand how changes in experimental design alter result interpretation, we must grasp the inherent framework and limitations of Perturb-seq, and then clarify the scope of controllability—specifically, the range of variables that can be manipulated through experimental design adjustments. Therefore, we first attempted to structure the rationale by which Perturb-seq provides mechanistic explanations for complex biological phenomena.

From a mechanistic perspective, the theoretical framework of Perturb-seq is illustrated in Figure S3. Perturb-seq is a framework that establishes KD models based on the C+LOO design. By comparing deviations from an "all-gene intact" baseline, it describes the relative contribution of each gene to the phenotype via a causal model. Due to its inherent nature, Perturb-seq cannot directly observe the intermediate phenomena occurring between the perturbation and the scRNA-seq readout. Specifically, this process is treated as a "black box" system, formulated as a causal model regarding the correspondence—or discrete mapping—between the input (perturbation condition) and the output (phenotypes evaluated via scalar metrics). While unobserved internal mechanisms remain a black box in principle, they can be inferred by leveraging the high-throughput nature of the data, similar to standard scRNA-seq analysis without a causal structure.

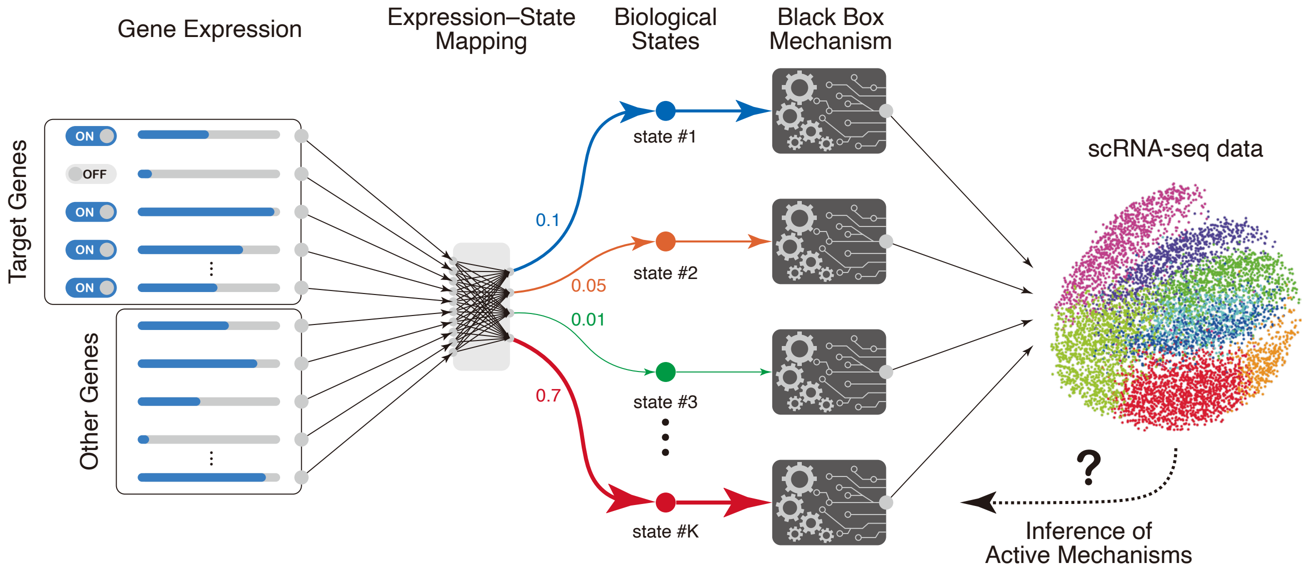


**Figure S3: A canonical theoretical framework for Perturb-seq**

In Perturb-seq, selected target genes are discretely controlled (e.g., ON/OFF), producing a gene-expression pattern at the time of perturbation from the combination of manipulated targets and all other genes. This expression pattern maps—often in a complex way—onto an ontology of biological states (e.g., Gene Ontology terms and related descriptors), enabling biological interpretation of cellular states from the transcriptome. For convenience, we assume a finite set of states, with each cell represented by membership probabilities over these states. Each ontology category is associated with mechanisms specific to the experimental system; perturbations preferentially engage mechanisms linked to the cell's state at the time of intervention. Because scRNA-seq is a snapshot without temporal information, the full sequence of active mechanisms between perturbation and measurement remains effectively a black box, though aspects can be inferred from the observed data. In practice, comparing baseline and perturbed conditions estimates the causal effect of the discrete input (the KD) on the output phenotype; mechanistic pathways are not identified directly by this causal contrast but are instead inferred downstream from the scRNA-seq readout.

A critical perspective here involves determining (i) what changes can occur in the underlying mechanisms, and (ii) distinguishing what can and cannot be examined within this framework of discrete causal modeling for inputs and outputs. To formally address the cellular changes induced by perturbations, we consider the temporal evolution of GRNs from the point of perturbation.

From the perspective of temporal GRN dynamics, post-perturbation system changes can be classified into two simple categories as follows:

1. Cases where the GRN structure exhibits condition-dependent temporal changes (Figure S4a)
2. Cases where the GRN structure remains static, with only gene expression levels changing in a condition-dependent manner (Figure S4b)

First, we consider systems where the GRN structure exhibits condition-dependent temporal evolution. As a concrete model case, Figure S4a illustrates the dynamic changes in the GRN within a Perturb-seq experiment targeting genes $X_1$–$X_6$. The initial condition is shown in the left panel; we assume the GRN remains static under the baseline (intact) condition. We considered the following three patterns of structural changes in the GRN that may occur between perturbation and observation:

- Sign reversal of gene–gene interactions: The case of $X_1$ KD. Between the intervention and observation, the regulatory directionality (positive/negative) of the paths $X_1 \to X_3$ and $X_1 \to X_2 \to X_6$ is reversed.
- Gain and loss of gene–gene interactions (Rewiring): The case of $X_2$ KD. The effect of the intervention on $X_2$ propagates to $X_5$ via a newly emerged path through $X_6$ ($X_6 \to X_5$). However, at the time of observation, the direct regulatory link $X_2 \to X_6$ has disappeared, obscuring the path taken to reach $X_6$.
- Coupling and decoupling at the module level: The case of $X_3$ KD. $X_3$ temporarily couples with an external GRN module, exerting an influence on $X_4$ through that pathway. However, since the original coupling is decoupled at the time of observation, this indicates that not only genes within the model but also unknown factors outside the model can act as confounders.

A common issue across these cases is that "point" observations via scRNA-seq cannot capture the structural changes occurring during the "blank time" between perturbation and observation. For instance, dynamic processes such as the timing of sign reversal or the transient generation and disappearance of pathways are theoretically impossible to infer from static snapshots. Consequently, the traceability of detailed mechanisms is lost, making accurate estimation of causal effects difficult.

Such condition-dependent temporal evolution of GRN structures is a phenomenon prominently observed in *biological processes* accompanied by dynamic remodeling of gene expression profiles; a concrete example is the temporal acquisition of responsiveness to glial-inducing stimuli in neural stem cells [TBR+15].

Of course, if the structural changes in the GRN occur with high reproducibility, it is possible to infer the causal relationship between "intervention and phenotype" using Perturb-seq. However, elucidating the full picture of the underlying mechanisms would require measurements at time intervals fine enough that the GRN structure remains effectively static. Furthermore, given that the true GRN structure is unknown in real data and must be inferred from noisy datasets before applying the above reasoning, this presents a formidable challenge when considering realistic experimental costs and technical constraints.

Next, let us consider a system where the GRN structure remains time-invariant, again using Perturb-seq targeting genes $X_1$–$X_6$ as a concrete example (Figure S4b). In this system, the structural changes described earlier—such as sign reversals of gene–gene interactions, network rewiring, or the coupling and decoupling of modules—do not occur. Instead, differences in KD conditions manifest solely as quantitative variations in the expression levels of $X_1$–$X_6$ within a fixed network topology.

The critical distinction between this system and one where the GRN structure varies dynamically with conditions lies in the static nature of the GRN. Because the structure is invariant, the system is robust against causal disconnections arising from the observational "blank time." Consequently, provided the input–response relationship is effectively explored, the underlying mechanism is, in principle, inferable. In other words, a static GRN is a prerequisite for the discrete causal models of Perturb-seq to be essentially solvable. Furthermore, it is only within this type of system that the efficiency of data space exploration can be meaningfully discussed using DOE concepts.

In contrast to systems exhibiting condition-dependent GRN transformation (conceptually corresponding to biological processes), the system described here maintains a consistent regulatory architecture without fundamental remodeling of the regulatory logic governing the cell's gene expression patterns. Such stability is typically observed in immediate responses, potentially associated with *biological events*.

Consequently, the discrete causal modeling of Perturb-seq is most rigorously applicable to these systems with fixed GRN structures. Moreover, the efficiency of DOE-based experimental design is theoretically grounded only within this stable domain. Therefore, the remainder of this paper focuses on systems exhibiting such static GRN topology.

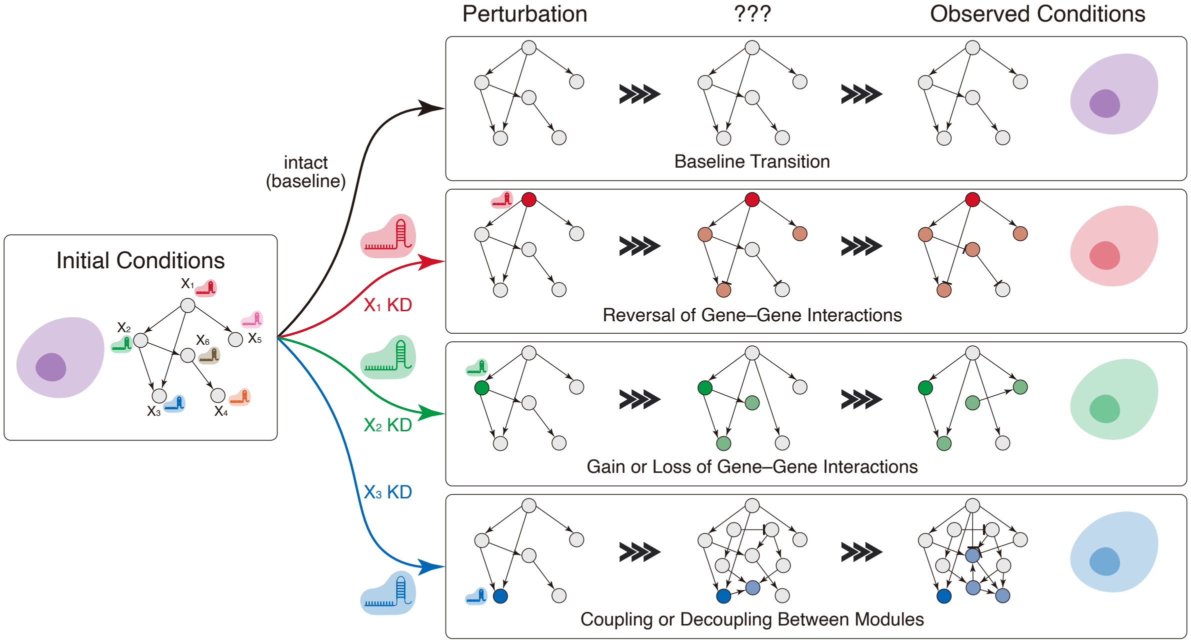


(a) Systems with time- and condition-dependent GRN dynamics

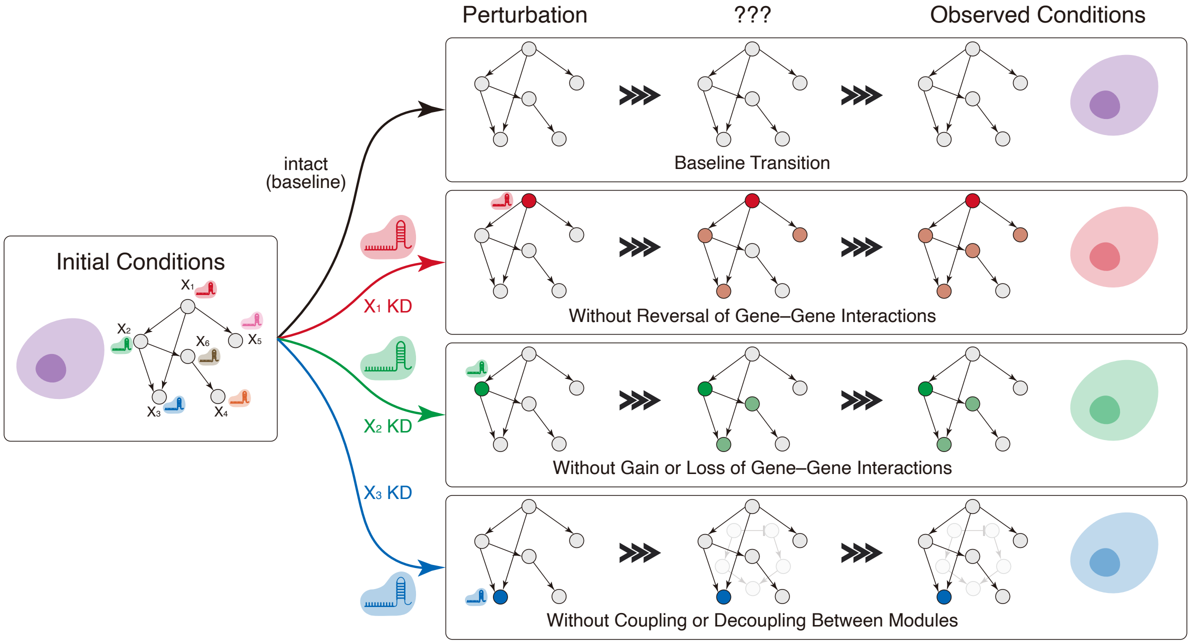


(b) Systems with a stationary GRN structure

**Figure S4: Temporal changes in GRN structure and system suitability for Perturb-seq**

**a–b**: Systems whose GRN structure reconfigures between perturbation and measurement in a condition-dependent, time-varying manner (a), versus systems with a stationary GRN (b). Because Perturb-seq observes only the GRN state at the measurement time, the dynamics in (a) cannot be traced (e.g., baseline transitions, sign reversals, edge gain/loss, or module coupling/decoupling). Thus, detailed mechanistic analysis with Perturb-seq is most appropriate when the GRN remains stationary across conditions, as in (b). For simplicity, we illustrate a baseline GRN over genes $X_1$–$X_6$ that is static; each row depicts a Perturb-seq experiment targeting one gene. Nodes in vivid colors (red/green/blue) mark the perturbed (KD) gene in each condition; nodes in dimmer hues indicate downstream genes affected by the perturbation. Arrows indicate positive regulation, and the T-shaped arrows indicate negative regulation.

#### D.2 Rationale for a minimalist simulator

To evaluate the performance of experimental designs in Perturb-seq, the most intuitive approach is to directly compare the outcomes obtained by applying multiple designs. Indeed, in the development of scRNA-seq analysis methods—such as dropout imputation [PMC+21] and batch integration [LBC+22]—benchmarking using multiple real-world datasets has become a standard practice. However, when comparing Perturb-seq experimental designs, several confounding factors and limitations arise, making evaluation based solely on real or existing virtual data insufficient. These challenges include: (i) confounding with batch effects; (ii) the unknowability of true gene effects and the true GRN structure; (iii) the vastness of the Perturb-seq data space; (iv) the prohibitive costs of exhaustive experiments; (v) technical limitations of generative model-based simulations; and (vi) the dependency of the C+LOO design on the number of factors.

First, isolating the differences caused by experimental design from batch effects is difficult. Because scRNA-seq measurements are sensitive to batch effects—systematic variations between sequencing runs that can arise even from subtle day-to-day changes in laboratory procedures [LBC+22]—discrepancies between Perturb-seq datasets cannot be attributed with confidence to experimental-design differences alone.

Second, the "true gene effects" and "true GRN structure," which serve as the gold standard for evaluation, remain unknown. In benchmarking experimental designs, an intuitive scheme would be to establish a ground truth for gene effects and assess how accurately each design reproduces it. However, given that gene functions are highly context-dependent [KPG+24], and scRNA-seq data inherently links samples and genes in a dual relationship (implying that observed gene functions are data-dependent), establishing an *a priori* ground truth for gene behavior under the specific conditions of Perturb-seq is extremely difficult. As discussed in Supplementary Information (SI) D.1, the success of Perturb-seq depends on temporal robustness of GRN structures; without a known ground truth for the GRN structure, the "anchor point" for comparing experimental designs is fundamentally unstable.

Third, the absence of this anchor point stems from the vastness of the Perturb-seq exploration space. While comprehensive databases such as the Human Cell Atlas [RTL+17] have been established for standard scRNA-seq, Perturb-seq involves the selection of target genes and often non-physiological conditions, resulting in a much larger data space. Consequently, it is difficult to determine the extent to which findings derived from a small number of datasets—focusing on specific components of a biological system—are generalizable to broader Perturb-seq research questions.

Fourth, from the perspective of DOE, using an FF design as an anchor point is a potential solution. Since the FF design covers all control experiments, it offers the highest resolution and could serve as an ideal benchmark. However, an FF design requires $N \cdot 2^n$ trials for $n$ factors and replication level $N$. For a large scale of $n$, the experimental cost becomes prohibitive (see SI C.1). Furthermore, establishing experimental conditions that achieve effective KD for up to $n$ simultaneous CRISPR targets while excluding side effects such as off-target activity presents significant technical difficulties.

Fifth, as an alternative, utilizing simulators such as GEARS [RHL24] or scGPT [CWM+24] to virtually generate Perturb-seq data for FF or other designs has been considered. However, their predictive accuracy for unseen two-gene perturbations remains limited, with reported correlation coefficients of around 0.2–0.6 even in state-of-the-art models [VTWP+25]. This underscores the difficulty in predicting unobserved combinatorial effects—especially nonlinear ones. Consequently, accurately simulating Perturb-seq experiments with desired experimental designs that require simultaneous multi-gene KD is currently unrealistic. Hence, it is difficult to regard reconstructed Perturb-datasets based on counterfactual experimental designs as sufficiently reliable for such evaluation.

Finally, the factor-number dependency of the de facto standard C+LOO design also hinders the generalization of empirical findings. The parameters related to the inherent correlation structure of the C+LOO design change according to the number of factors, $n$. Even if one were to perform benchmarking using a small $n$ where establishing an FF design anchor is feasible, the properties of the C+LOO design in that context would differ from those in high-throughput screenings (large $n$), making the results likely inapplicable. While testing across different scales of $n$ might seem like a solution, when combined with the aforementioned challenges, it raises the barrier for systematic comparison using empirical methods.

In light of these challenges, this study adopts a "minimalist simulator" approach that abstracts the essence of the Perturb-seq process, rather than relying on empirical benchmarking with real datasets. Through this approach, we aim to approach universal laws governing the relationship between experimental design performance and underlying GRN mechanisms, ultimately providing guiding principles to assist decision-making in future Perturb-seq research planning.

##### D.3 Taylor's power law (TPL) suggests our simulators model "canonical behavior" of biological systems

As described in Methods 4.3, our minimalist simulator architecture is composed of the following six steps: (i) computation of a linear combination of upstream factors ($b_\ell x_{i,j}$); (ii) addition of baseline expression potentials ($v_{i,j}$) and Gaussian noise ($\varepsilon_{i,j}$); (iii) modulation by perturbation status ($c_{i,j}$); (iv) application of a ReLU nonlinearity to obtain $x_{i,j}$; (v) computation of a second linear combination ($a_k x_{i,j}$) plus observation noise $\varepsilon_{y,j}$; and (vi) a final ReLU yielding $y_j$. This structure preserves the explainability of linear combinations, while the ReLU steps mimic biological nonlinearities [JKRL09, NIGM18] without sacrificing the advantages of linear aggregation. Specifically, adopting ReLU functions creates a structure analogous to real-valued Boolean networks used in signaling models [KK22]. Additionally, parameters ($a_k$, $b_\ell$, $v_{i,j}$) are sampled from log-normal distributions to capture biological overdispersion.

To assess whether the simulators reproduce realistic variability, we tested whether their outputs follow TPL

$$\log \mathrm{Var}[X] = \log \alpha + \beta \log \mathrm{E}[X], \tag{S-47}$$

where $\alpha$ and $\beta$ are fitted. TPL is widely observed in biological systems exhibiting heavy-tailed distributions [Tay61, GFR+15, LR21]. In the context of single-cell transcriptomics, count data are standardly modeled using Poisson ($\beta = 1$) or Negative Binomial distributions ($\beta > 1$) [GKVO14, ATK+18], representing a canonical baseline for realistic overdispersion.

We ran the FF-based simulations with $N = 10$ replicates for Models $\Phi$, $\Psi$, and $\Lambda$ under four Gaussian noise levels ($\sigma \in \{0.5, 1, 2, 4\}$). For each experiment we computed the mean and variance of the output values per experimental conditions and fitted the log-log linear model (Figure S5). All panels yielded $R^2 > 0.6$—supporting adherence to TPL—and slopes $\beta \approx 1$ or slightly higher. This confirms that our simulators capture the Poisson-like stochasticity and structural overdispersion characteristic of real-world scRNA-seq data.

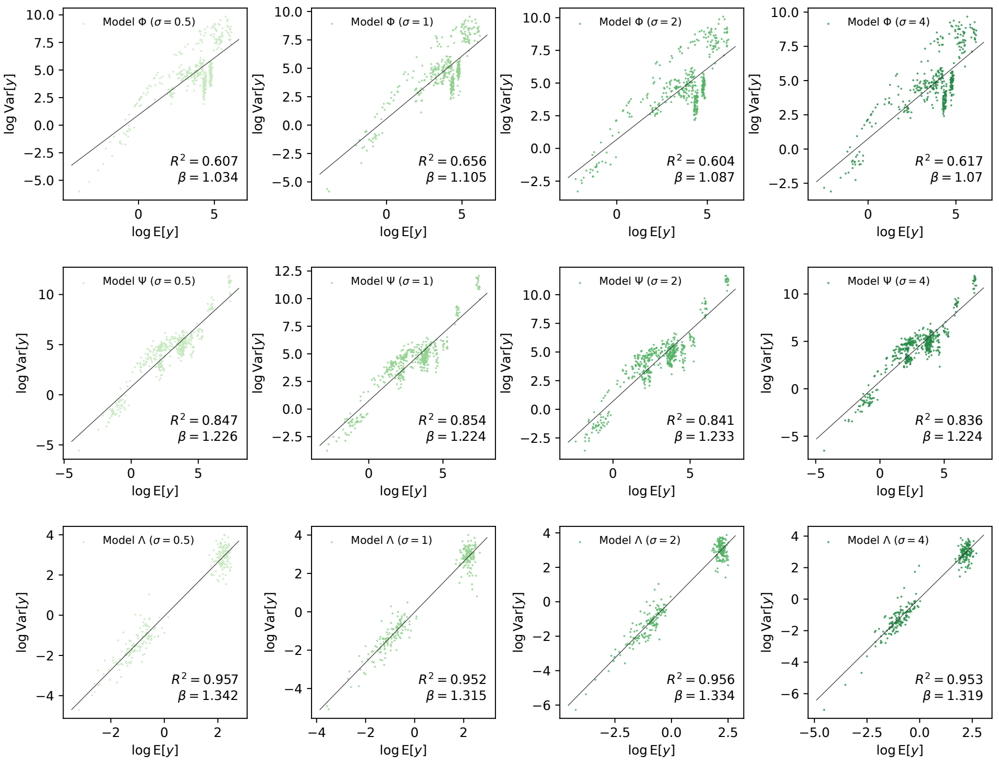


**Figure S5: TPL holds across the simulated models**

Each panel shows the relationship between the natural-log-transformed mean (horizontal axis) and variance (vertical axis) of the output values $y$. Rows correspond to Models $\Phi$ (top), $\Psi$ (center), and $\Lambda$ (bottom); columns correspond to Gaussian noise levels $\sigma = 0.5$ (left), $\sigma = 1$ (second left), $\sigma = 2$ (second right), and $\sigma = 4$ (right). The slopes ($\beta$) and $R^2$ of the linear regression models are reported in each panel.

#### E A generic workflow of C+LOO-based Perturb-seq vs. DOE-originated multivariate analysis

In a generic workflow of Perturb-seq experiments, genes are knocked down one at a time, and the effects are inferred by comparing the results of each single-gene perturbation against a control group. Typically, wild-type or sham conditions serve as this baseline. This pairwise comparison framework allows for an intuitive interpretation of gene function: one simply checks whether the KD of a specific gene causes a significant deviation from the intact state.

We demonstrated this standard approach using simulated Perturb-seq experiments with the C+LOO design (Model $\Phi$, $N = 3$). Following standard protocols, we performed Dunnett's test to compare the mean output values of single-gene KDs against the "all-genes-intact" control (Figure S6a). Since knocking down a regulator is expected to produce an effect opposite to its function, causal effects can be identified by detecting significant differences. This scheme effectively serves as a biologically meaningful proof-of-concept, directly linking the absence of a single gene to a phenotypic change. This intuitiveness is a unique advantage of the C+LOO design, whereas PB and other DOE-based designs often involve complex combinatorial perturbations that prioritize statistical efficiency over such direct interpretability.

Despite its straightforward nature, however, this pairwise approach creates a statistical bottleneck. From a DOE perspective, the standard Perturb-seq analysis implicitly treats multiple genetic perturbations as parallel levels of a single manipulation factor (i.e., an $(n + 1)$-level single-factor model). This structure makes inference heavily baseline-dependent. For instance, in a system with two factors A and B, pairwise comparison evaluates the effect of A by strictly comparing the "double-positive" (control) and "B-positive/A-negative" (A-KD) conditions (Figure S6d). Consequently, the effect of A is essentially evaluated under the specific influence of B.

This dependency becomes problematic when the baseline condition itself is unstable or heterogeneous. For example, in studies targeting highly heterogeneous diseases like glioblastoma multiforme[1], the effect of a KD might not be reproducible if the baseline gene expression profile varies between batches or samples. Furthermore, batch effects—systematic differences arising from experimental protocols or biological features—are a general concern in scRNA-seq [LBC+22]. Since individual perturbations are often replicated across different batches, relying on a single control condition as the absolute reference point renders the analysis vulnerable to such fluctuations.

In contrast, analyzing two-level $n$-factor designs with multivariate methods (MLR or ANOVA) allows for a neutral estimation of main effects (Figures S6b–S6c). Here, the main effect of a factor is defined as its average contribution independent of other factors [CSC20]. Returning to the two-factor example (A and B), the main effect of A is calculated by averaging (i) the difference between the "double-positive" and "B-positive/A-negative" conditions, and (ii) the difference between the "A-positive/B-negative" and "double-negative" conditions (Figure S6d). Crucially, no single condition acts as the exclusive baseline. This symmetric formulation effectively cancels out the influences of other variables (like B) and accommodates data variations, providing a robust framework for dealing with the noise inherent in scRNA-seq data. Indeed, our benchmarking results confirm the superiority of multivariate models over pairwise comparisons in terms of robustness to Gaussian noise (SI G.4).

However, applying this multivariate framework to the C+LOO design reveals a fundamental structural limitation: the lack of sufficient control conditions. DOE-based screening designs (e.g., FF or PB) are constructed to ensure orthogonality or near-orthogonality, containing a balanced set of conditions that allows main effects and interactions to be mathematically isolated. The C+LOO design, by definition, lacks the "multiple-KD" ("double-negative" in Figure S6d) conditions essential for canceling out influences from other factors. This imbalance creates a correlation structure inherent to C+LOO, meaning that estimates of main effects can bleed into interaction terms, and vice versa. As exemplified in Figures S6b–S6c, the estimated main effects can differ drastically depending on the experimental design, even when using the same multivariate analysis.

We evaluate design performance using MLR and ANOVA as the primary engines to compare C+LOO and PB from a DOE standpoint. Although ANOVA is formally undefined for C+LOO with $N = 1$, and one might resort to graphical methods like half-normal plots (Figure S6e) for effect screening [WG68, Dan59], we adopt a unified evaluation based on MLR/ANOVA under matched replication to ensure strict comparability.

#### F Impact of the inherent correlations: Interaction Leakage in the C+LOO design

When a variable $\mathbf{v} \sim \mathcal{N}_n(\boldsymbol{\mu}, \sigma^2 \mathbf{I}_n)$, and $X$ is the model matrix, the following equations hold:

$$\phi := \operatorname{rank}(X) \tag{S-48}$$

[1] a WHO grade IV adult brain tumor characterized by high levels of intertumoral and intratumoral heterogeneity [WKZ+21]

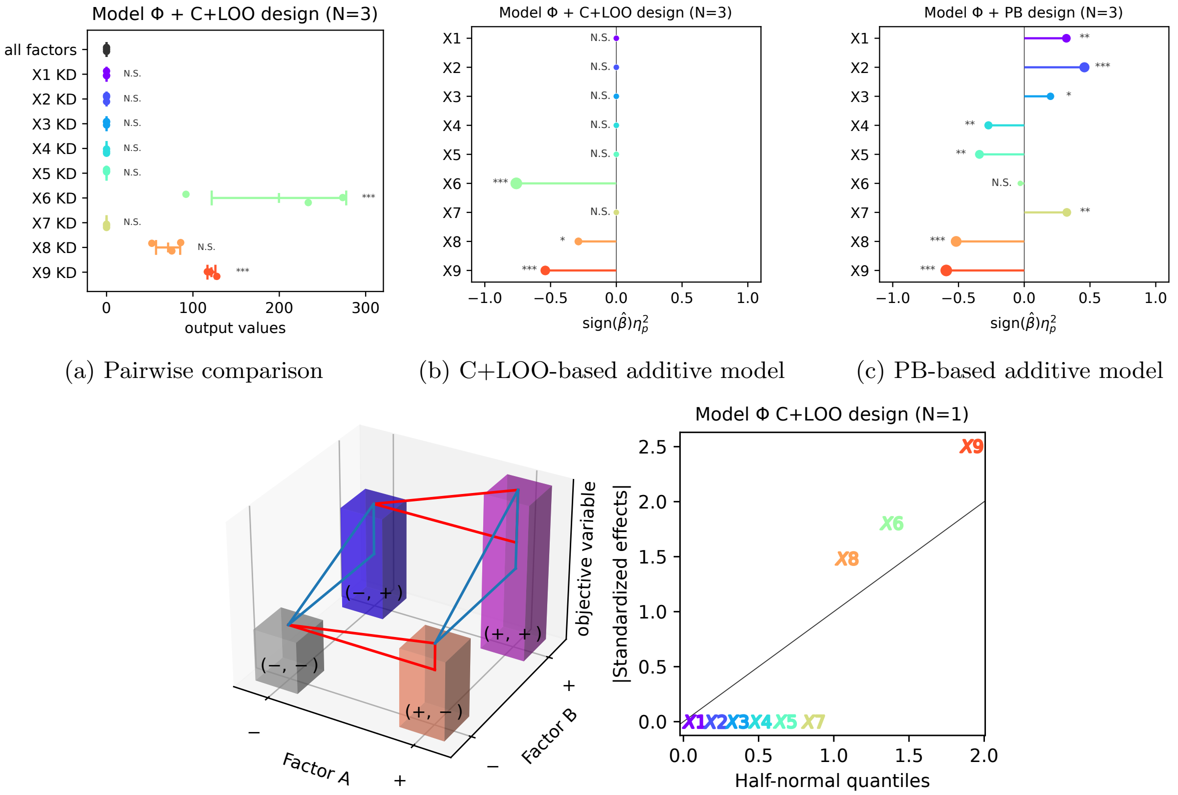


(a) Pairwise comparison (b) C+LOO-based additive model (c) PB-based additive model

(d) A $2 \times 2$ experiment (e) Half-normal plot

**Figure S6: Simulated experiments highlight the pros and cons of the C+LOO design**

**a**: The mean± SD values of the simulated Model $\Phi$ results from C+LOO-based experiments at $N = 3$. Each row represents the corresponding condition: either "all factors" (baseline) or a specific factor KD. Dunnett's test was performed against the "all factors" control ($*$, $p < 0.05$; $**$, $p < 0.01$; $***$, $p < 0.001$; N.S., not significant). The raw data points are overlaid. Note that KDs produce effects opposite to the genes' regulatory roles (e.g., "X6 KD" showing a significantly higher mean than "all factors" indicates that X6 functions as a downregulator). **b**–**c**: The signed partial eta-squared values ($\mathrm{sign}(\hat{\beta})\eta_p^2$), calculated as the product of the MLR coefficient sign ($\mathrm{sign}(\hat{\beta})$) and the partial eta-squared ($\eta_p^2$), for the additive models of Model $\Phi$ ($N = 3$). Statistical significance was determined using ANOVA ($*$, $p < 0.05$; $**$, $p < 0.01$; $***$, $p < 0.001$; N.S., not significant). The sign of the metric reflects the direction of the suggested regulatory effects. Marker sizes are scaled by $-\log_2 p$. **d**: An example representation of a $2 \times 2$ experiment. The presence (+) or absence (−) of the two factors, A and B, is indicated by the coordinates of the bars, which correspond to the output values of the associated conditions. **e**: The half-normal plot for the C+LOO-based Model $\Phi$ experiment at $N = 1$. The effects are suggested to be significant when the absolute values of the standardized effects (shown on the vertical axis) are larger than the half-normal quantiles (shown on the horizontal axis).

$$\lambda := \frac{\|P(X)\boldsymbol{\mu}\|^2}{\sigma^2} \tag{S-49}$$

$$\frac{\mathbf{v}^\top P(X)\mathbf{v}}{\sigma^2} \sim \chi^2_\phi(\lambda), \tag{S-50}$$

where $P(X) := X(X^\top X)^{-1}X^\top$ is the orthogonal projection matrix of $X$ and $\chi^2_\phi(\lambda)$ denotes a noncentral chi-squared distribution with $\phi$ degrees of freedom and a noncentrality parameter $\lambda$. Notably, the following equations hold for the definition of the orthogonal projection matrix:

- idempotency

$$P(X)^2 = P(X) \tag{S-51}$$

- symmetry

$$P(X)^\top = P(X) \tag{S-52}$$

As described in Methods 4.4.4, the error sum of squares $SS_E$ is defined as follows:

$$SS_E := \mathbf{y}^\top(\mathbf{I}_{n_{\max}} - P(X_\Omega))\mathbf{y}, \tag{S-53}$$

where $X_\Omega := [\,\mathbf{1}\ \mathbf{x}_1\ \cdots\ \mathbf{x}_n\,]$. Since $(\mathbf{I}_{n_{\max}} - P(X_\Omega))$ is also an orthogonal projection matrix showing idempotency and symmetry, the following equations hold:

$$\phi_E := n_{\max} - \operatorname{rank}(X_\Omega) \tag{S-54}$$

$$\lambda_E := \frac{\|(\mathbf{I}_{n_{\max}} - P(X_\Omega))\boldsymbol{\mu}\|^2}{\sigma^2} \tag{S-55}$$

$$\frac{SS_E}{\sigma^2} \sim \chi^2_{\phi_E}(\lambda_E), \tag{S-56}$$

where $\mathbf{y} \sim \mathcal{N}_{n_{\max}}(\boldsymbol{\mu}, \sigma^2\mathbf{I}_{n_{\max}})$.

The size of $\lambda_E$ is controlled by the model equation. When there is no influence of interaction terms to $\boldsymbol{\mu}$—i.e., $\boldsymbol{\mu} = X_\Omega\boldsymbol{\beta}_\Omega$—$\lambda_E$ is canceled as follows:

$$\begin{aligned}\lambda_E &= \frac{\boldsymbol{\mu}^\top(\mathbf{I}_{n_{\max}} - P(X_\Omega))^\top(\mathbf{I}_{n_{\max}} - P(X_\Omega))\boldsymbol{\mu}}{\sigma^2}\\ &= \frac{\boldsymbol{\beta}_\Omega^\top X_\Omega^\top(\mathbf{I}_{n_{\max}} - P(X_\Omega))^\top(\mathbf{I}_{n_{\max}} - P(X_\Omega))X_\Omega\boldsymbol{\beta}_\Omega}{\sigma^2}\\ &= 0.\end{aligned} \tag{S-57}$$

Therefore, $SS_E/\sigma^2 \sim \chi^2_{\phi_E}$, a chi-squared distribution without any noncentrality parameter. However, if there are active interactions—thus, $\boldsymbol{\mu} = X_\Omega\boldsymbol{\beta}_\Omega + X_{\text{int}}\boldsymbol{\beta}_{\text{int}}$ where $X_{\text{int}}$ and $\boldsymbol{\beta}_{\text{int}}$ refer to either the model matrix or the coefficient vector for the interaction terms—$\lambda_E$ is formulated as follows:

$$\begin{aligned}\lambda_E &= \frac{\|(\mathbf{I}_{n_{\max}} - P(X_\Omega))(X_\Omega\boldsymbol{\beta}_\Omega + X_{\text{int}}\boldsymbol{\beta}_{\text{int}})\|^2}{\sigma^2}\\ &= \frac{\|(\mathbf{I}_{n_{\max}} - P(X_\Omega))(X_{\text{int}}\boldsymbol{\beta}_{\text{int}})\|^2}{\sigma^2}.\end{aligned} \tag{S-58}$$

In regular Type II ANOVA, $\boldsymbol{\beta}_{\text{int}}$ is assumed to be $\mathbf{0}$ so that $F_i := (SS(X_i)/\phi_i)/(SS_E/\phi_E)$ follows a noncentral $F$-distribution $F_{\phi_i,\phi_E}(\lambda_i)$; otherwise, $F_i$ follows a doubly noncentral $F$-distribution $F_{\phi_i,\phi_E}(\lambda_i, \lambda_E)$.

Furthermore, the model matrix can shape the size of $\lambda_E$. Other than $\boldsymbol{\beta}_{\text{int}}$ and $\sigma$, the difference between $X_{\text{int}}^\top X_{\text{int}}$ and $X_{\text{int}}^\top P(X_\Omega)X_{\text{int}}$ plays the deterministic role in the scale of $\lambda_E$ as follows:

$$\begin{aligned}\lambda_E &= \frac{\boldsymbol{\beta}_{\text{int}}^\top X_{\text{int}}^\top(\mathbf{I}_{n_{\max}} - P(X_\Omega))X_{\text{int}}\boldsymbol{\beta}_{\text{int}}}{\sigma^2}\\ &= \frac{\boldsymbol{\beta}_{\text{int}}^\top(X_{\text{int}}^\top X_{\text{int}} - X_{\text{int}}^\top P(X_\Omega)X_{\text{int}})\boldsymbol{\beta}_{\text{int}}}{\sigma^2}.\end{aligned} \tag{S-59}$$

Interestingly, $X_{\text{int}}^\top X_{\text{int}} - X_{\text{int}}^\top P(X_\Omega)X_{\text{int}}$ can be measured exclusively from the model matrix, but without needing any parameters from observations. Since $X_{\text{int}}^\top P(X_\Omega)X_{\text{int}}$ represents the influences from interaction terms projected in the model based on $X_\Omega$, we created a metric *Interaction Leakage* to evaluate the impact of $X_{\text{int}}^\top P(X_\Omega) X_{\text{int}}$ relative to $X_{\text{int}}^\top X_{\text{int}}$ as follows:

$$\text{Interaction Leakage} := \frac{\text{tr}(X_{\text{int}}^\top P(X_\Omega) X_{\text{int}})}{\text{tr}(X_{\text{int}}^\top X_{\text{int}})}. \tag{S-60}$$

For example, we computed the Interaction Leakage for the C+LOO, PB, and FF designs at numbers of factors $1 \leq n \leq 12$ when assign the second-order interactions to $X_{\text{int}}$ (Figure S7). The C+LOO designs constantly show high Interaction Leakage, suggesting the "leaky" nature of the C+LOO design where the interaction terms percolated into the additive models via factor correlations even without explicitly assuming them.

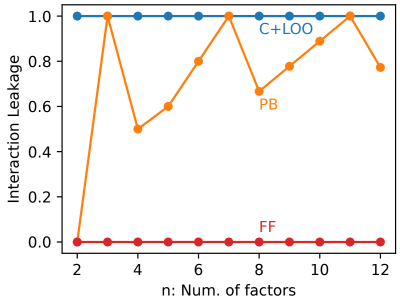


**Figure S7: Interaction leakage of the experimental designs**

The Interaction Leakage values for the additive model matrices $X_\Omega$ and the second-order interactions $X_{\text{int}}$ based on the C+LOO (blue), PB (orange), and FF (green) designs at different numbers of factors ($1 \leq n \leq 12$).

Notably, the C+LOO and PB designs at $n \geq 3$ show non-zero Interaction Leakage, which decrease $\lambda_E$. Accordingly, we assumed $\lambda_E \approx 0$ and performed ANOVA avoiding doubly noncentral $F$-distributions.

### G Correspondence between simulator configurations and experimental design performance

To investigate the behavior of the C+LOO across multiple conditions, we varied the Gaussian noise terms in Models $\Phi$, $\Psi$, and $\Lambda$ to compare the performance of the C+LOO and PB designs.

To understand how the factors (X1–X9) influence the output value across different noise conditions ($\sigma \in \{0.5, 1, 2, 4\}$), we generated a hundred different FF-based simulations per noise level and computed the main effects according to their definitions. Based on their 95% bootstrap CIs, we classified the nine factors into the three categories: upregulators (denoted as "Up"), downregulators (denoted as "Down"), and non-significant effects (denoted as "N.S.").

The estimated main effects were regarded as ground truths, and the performance of the C+LOO and PB designs was compared in terms of the consistency of result interpretation with these ground truths. Weighted Cohen's $\kappa$ was calculated from ANOVA models under varying replication levels ($N$) and Gaussian noise conditions. Since pairwise comparison is the standard analytical approach, C+LOO-based Dunnett's test models were also benchmarked. Power analyses were conducted for all ANOVA and Dunnett's test models.

Notably, the Gaussian noise level was set to $\mathcal{N}(0, 1)$[2] in the main text; thus, we focus on the other conditions to see if the characteristics of the two designs are preserved across different conditions.

#### G.1 Model $\Phi$: a basic model simulating signaling cascades

First, we established Model $\Phi$, incorporating nine factors (X1–X9) that regulate the output value through multiple parallel pathways. To determine which attributes of signaling cascades influence the performance of C+LOO designs, we assigned upregulators, downregulators, and dual-function regulators (capable of both up- and down-regulation) to replicate the complex regulatory mechanisms found in biology. The estimated ground truth labels for the main effects across different noise levels are shown in Figure S8a.

As shown in Figures S8b–S8c, the PB-based models yielded higher $\kappa$ values than the C+LOO-based models across all replication levels and noise conditions, whereas the two designs exhibited similar $1 - \beta$ curves. This

[2]in other words, $\sigma^2 = 1$

suggests that the C+LOO design is not an ideal choice for systems driven by multiple independent cascades with predominantly additive effects, as represented in Model Φ.

##### G.2 Model Ψ: a highly complex regulatory circuit of interactions

To simulate a scenario with a more intricate network of interactions than Model Φ, we developed Model Ψ, which represents a phenomenon regulated by a highly complex cascade involving both cooperative and antagonistic interactions. The ground truth main effects of the nine factors are shown in Figure S9a.

In this model, the C+LOO-based models generally yielded higher $\kappa$ values than the PB-based models across almost all replication numbers and Gaussian noise levels (Figure S9b)—even though the two designs showed similar patterns in the $1-\beta$ curves (Figure S9c). Given that $\rho_{i\times j,i} = \frac{2}{3}$ for the C+LOO design of nine factors, the leaky behavior of the C+LOO design—where main effects are estimated under the influences of their associated interaction terms (SI F)—may have aligned well with the complex interaction patterns represented in Model Ψ.

##### G.3 Model Λ: a model with poorly curated candidates

Model Λ was established to simulate a scenario where the candidate genes are poorly filtered so that the majority of them show no influence on the target phenomenon. As intended, the main effects were identified as "N.S." according to the definition, except for X1 and X2 (Figure S10a).

In terms of experimental design performance, C+LOO slightly outperformed PB under low noise, whereas the performance of PB gradually improved as the Gaussian noise level increased, eventually outperforming C+LOO at $\sigma = 4$ (Figure S10b). The performance decline of C+LOO-based models and the relative superiority of PB-based models under high Gaussian noise may be attributable to the instability of variance partitioning in sparse factor scenarios. When the number of effective factors is significantly lower than the design dimension, the balance between signal and error variances becomes highly sensitive to detailed model parameters such as coefficients and noise levels. Therefore, sparse networks might be uniquely difficult when predicting the relative performance between C+LOO and PB designs solely based on network structures.

Interestingly, the PB-based ANOVA models for X1 and X2 maintained their statistical power even under high noise levels (Figure S10c), which shows the noise stability of the PB design.

##### G.4 C+LOO-based multivariate models vs. Dunnett's test models

As shown in Figures S8–S10, in simulations based on the C+LOO design, the performance of the multivariate models and Dunnett's test-based models was comparable, although the multivariate models performed slightly better overall. We attribute this similarity to the "leaky" structure of the design (SI F), in which the estimated main effects in C+LOO-based multivariate models remain implicitly influenced by interaction effects. In the power analysis, however, the multivariate models demonstrated higher statistical power, particularly under high-noise conditions. These results suggest that substituting the conventional pairwise comparison-based Perturb-seq analysis with a DOE-based multivariate analysis is not only feasible but also offers a robust alternative that improves stability against noise.

#### H Investigating the role of C+LOO-inherent correlations via D-optimization

In SI G, we found that the network structure of the simulated model is a key determinant of experimental design performance. Experimental designs with distinct correlation profiles offered respective advantages in different simulators depending on the network structure of the governing mechanisms.

To isolate and evaluate the importance of correlation structures in the experimental design performance, we applied D-optimization to correct the correlation patterns in C+LOO designs and examined how this modification affected the performance in terms of the $\kappa$ values to the ground truth.

With D-optimization, experimental designs can be augmented with additional runs to improve the D-criterion value (Figures S11a–S11b) and reduce factor correlations[3]. We applied D-optimization to the C+LOO designs for nine factors at $N = 3$ across various levels, resulting in total numbers of trials $n_{\max} \in \{36, 39, 42, 45, 48\}$. As expected, the D-criterion values (indicative of reduced correlations) decrease as $n_{\max}$ increases (Figure S11c). As a result, the DO-C+LOO design exhibited improved D-optimality with increasing $n_{\max}$ but could not outperform the original PB design[4] in terms of D-criterion values (Figure S11d).

[3] Since D-criterion shows the degree of factor independence [dABK+95], we leveraged it as an indicator of factor correlation.
[4] The superiority of the PB designs in terms of D-criterion can be derived from the Hadamard-matrix-based structure.

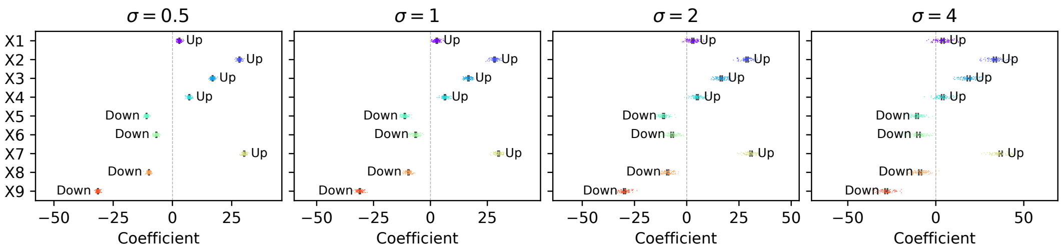


(a) Ground truth main effects

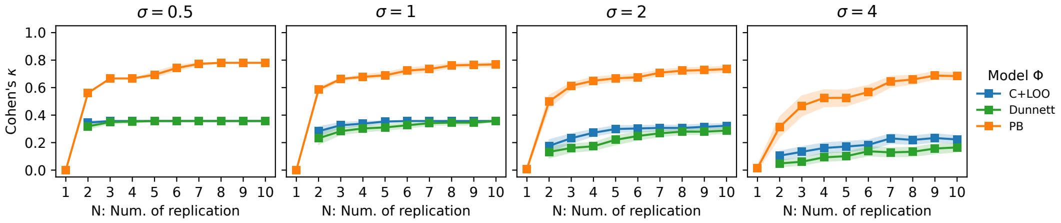


(b) Benchmarking

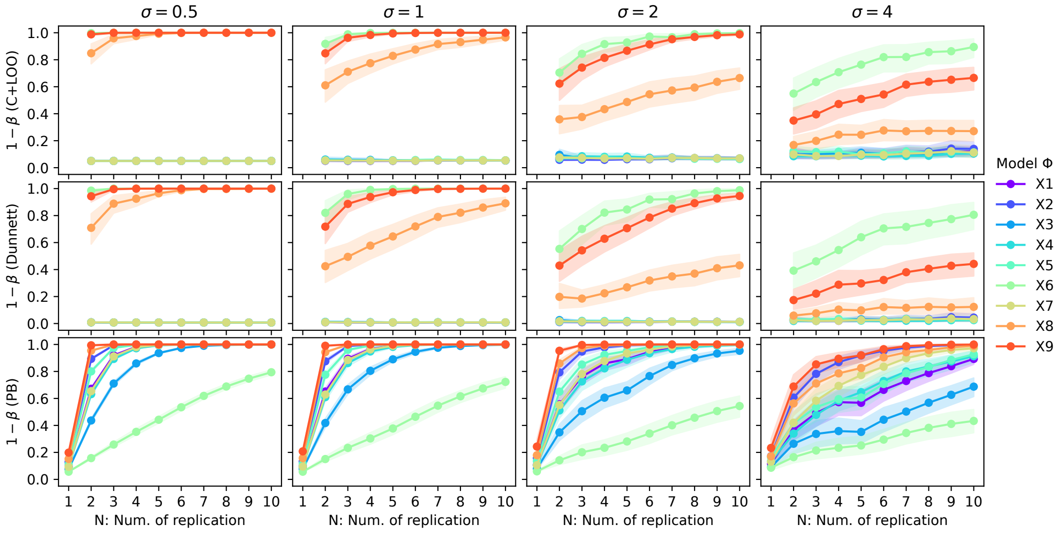


(c) Power analysis

**Figure S8: Benchmarking experimental designs with Model Φ**

**a**: Main effects of X1–X9 in Model Φ estimated from the FF-based simulations according to the definition of the main effect. The plots represent 100 different results generated with distinct random seeds and the error bars show the 95% bootstrap CIs (the number of bootstraps $n_{\text{boot}} = 10000$). Based on the CIs, we assigned interpretations (Up, N.S., or Down). **b**: Line charts showing $\kappa$ values for the C+LOO (blue) and PB (orange) designs as well as C+LOO-based Dunnett's test (green) at replication levels $1 \leq N \leq 10$. The Gaussian noise terms $\varepsilon_{i,j}, \varepsilon_{y,j}$ were generated from the normal distributions of different standard deviations: $\sigma \in \{0.5, 1, 2, 4\}$. **c**: The statistical power $1 - \beta$ of ANOVA for the C+LOO (top panels) and PB (bottom panels) designs and Dunnett's tests (middle panels) performed in Figure S8b at $\alpha = 0.05$. In Figures S8b–S8c, the markers show the mean of results from 30 different random seeds, and the shaded areas represent the 95% bootstrap CIs ($n_{\text{boot}} = 1000$).

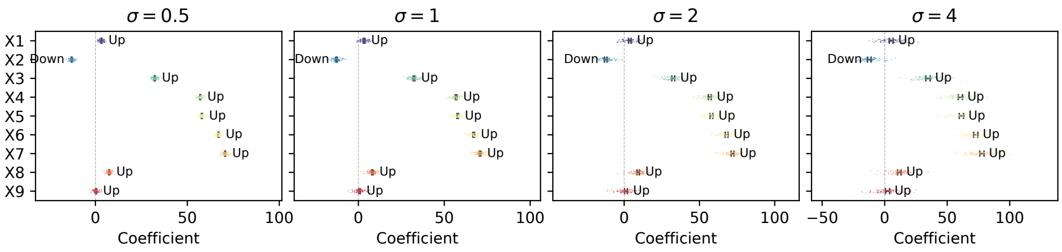


(a) Ground truth main effects

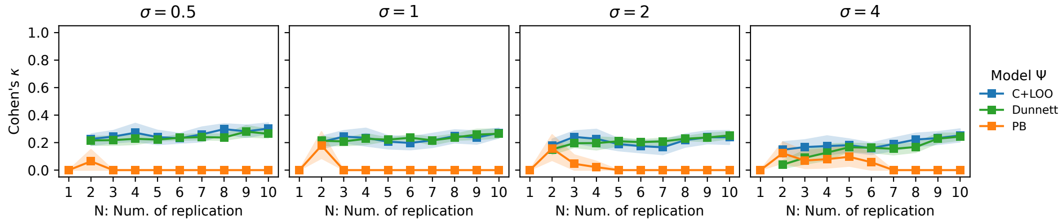


(b) Benchmarking

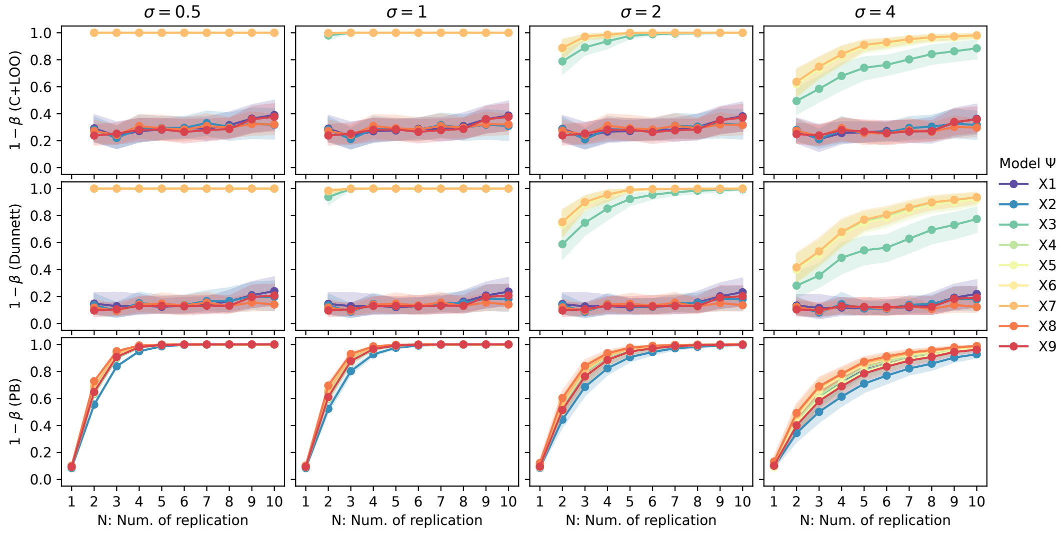


(c) Power analysis

**Figure S9: Benchmarking experimental designs with Model Ψ**

**a**: Main effects of X1–X9 in Model Ψ estimated from the FF-based simulations according to the definition of the main effect. The plots represent 100 different results generated with distinct random seeds and the error bars show the 95% bootstrap CIs (the number of bootstraps $n_{\text{boot}} = 10000$). Based on the CIs, we assigned interpretations (Up, N.S., or Down). **b**: Line charts showing $\kappa$ values for the C+LOO (blue) and PB (orange) designs as well as C+LOO-based Dunnett's test (green) at replication levels $1 \leq N \leq 10$. The Gaussian noise terms $\varepsilon_{i,j}, \varepsilon_{y,j}$ were generated from the normal distributions of different standard deviations: $\sigma \in \{0.5, 1, 2, 4\}$. **c**: The statistical power $1 - \beta$ of ANOVA for the C+LOO (top panels) and PB (bottom panels) designs and Dunnett's tests (middle panels) performed in Figure S9b at $\alpha = 0.05$. In Figures S9b–S9c, the markers show the mean of results from 30 different random seeds, and the shaded areas represent the 95% bootstrap CIs ($n_{\text{boot}} = 1000$).

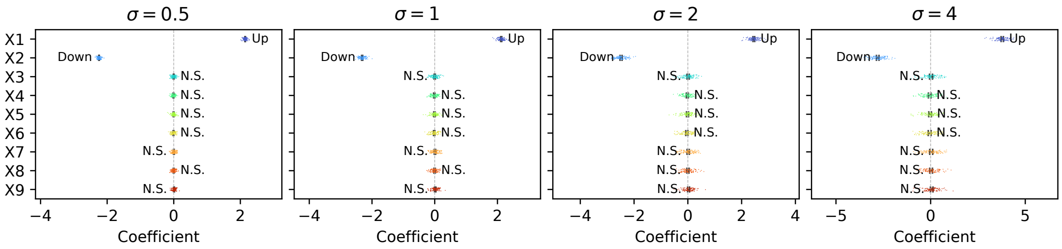


(a) Ground truth main effects

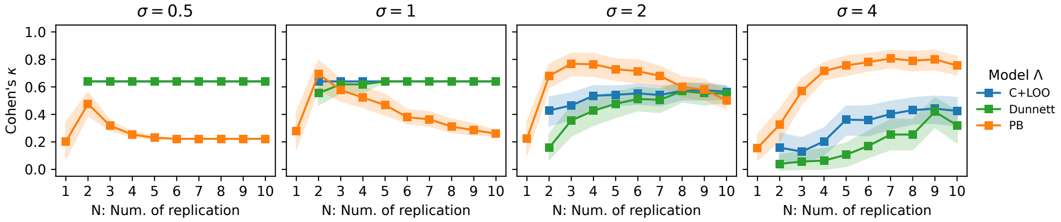


(b) Benchmarking

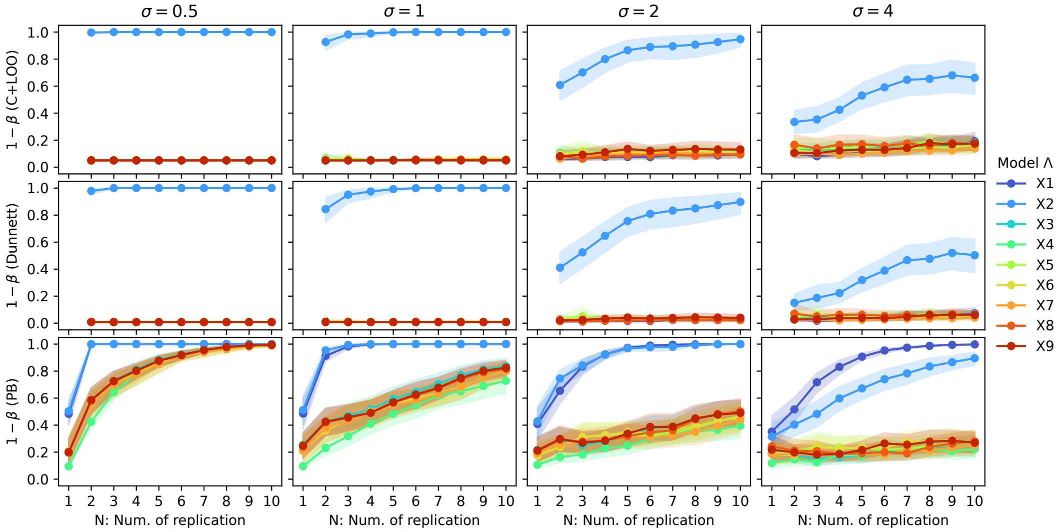


(c) Power analysis

**Figure S10: Benchmarking experimental designs with Model Λ**

**a**: Main effects of X1–X9 in Model Λ estimated from the FF-based simulations according to the definition of the main effect. The plots represent 100 different results generated with distinct random seeds and the error bars show the 95% bootstrap CIs (the number of bootstraps $n_{\text{boot}} = 10000$). Based on the CIs, we assigned interpretations (Up, N.S., or Down). **b**: Line charts showing $\kappa$ values for the C+LOO (blue) and PB (orange) designs as well as C+LOO-based Dunnett's test (green) at replication levels $1 \leq N \leq 10$. The Gaussian noise terms $\varepsilon_{i,j}, \varepsilon_{y,j}$ were generated from the normal distributions of different standard deviations: $\sigma \in \{0.5, 1, 2, 4\}$. Note that the C+LOO-based multivariate models and Dunnett's test models showed identical curves under $\sigma = 0.5$. **c**: The statistical power $1 - \beta$ of ANOVA for the C+LOO (top panels) and PB (bottom panels) designs and Dunnett's tests (middle panels) performed in Figure S10b at $\alpha = 0.05$. In Figures S10b–S10c, the markers show the mean of results from 30 different random seeds, and the shaded areas represent the 95% bootstrap CIs ($n_{\text{boot}} = 1000$).

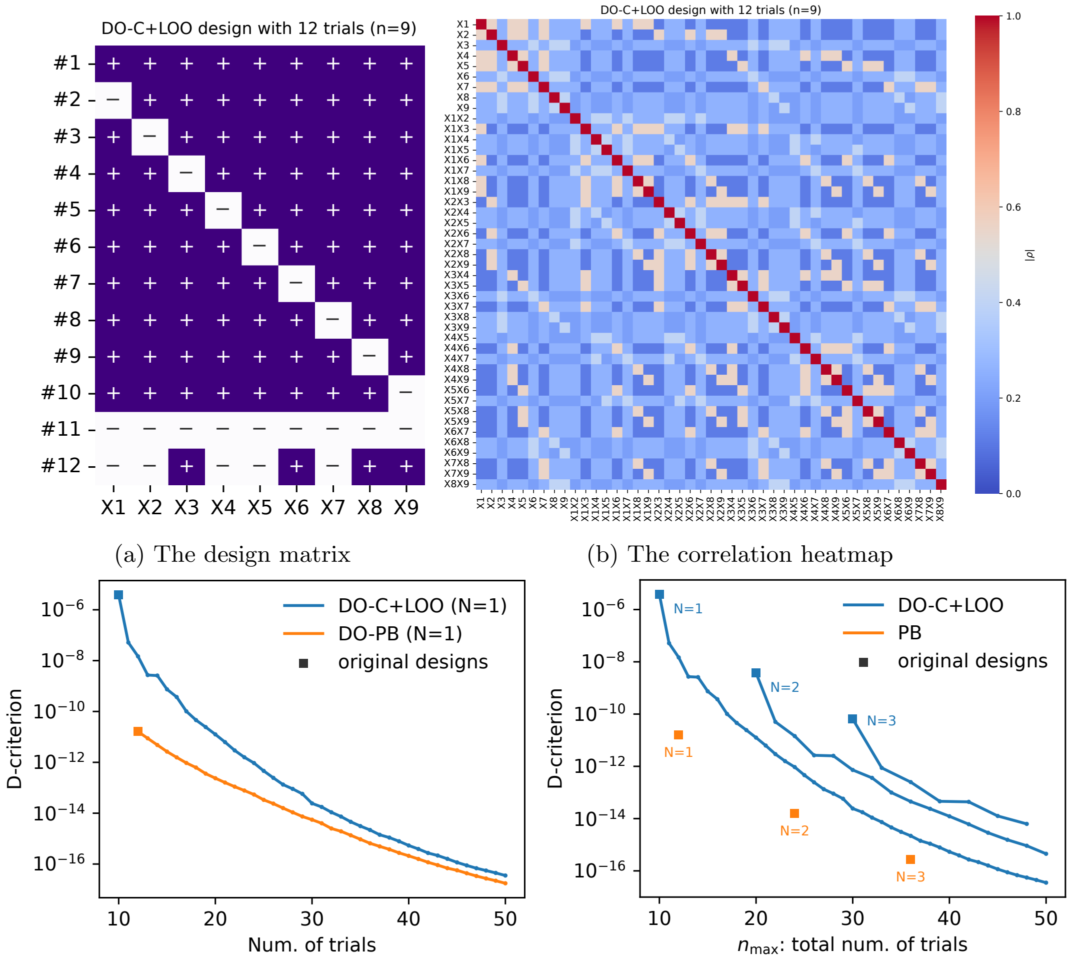


(a) The design matrix (b) The correlation heatmap

(c) DO-C+LOO vs DO-PB (d) DO-C+LOO vs PB

**Figure S11: Construction and D-optimality evaluation of the DO-C+LOO design**

**a–b**: Examples of the DO-C+LOO design matrix for nine factors with 12 trials (+, addition of factors; −, subtraction of factors) and the heatmap of $|\rho|$—the absolute values of the correlation coefficients—among main effects and interactions. **c–d**: D-criterion values for the model matrices of DO-C+LOO-based (blue) and DO-PB-based (orange) multivariate models for nine factors with different numbers of trials at (c) $N = 1$ and (d) multiple replication levels ($N = 1$–$3$). The square markers represent the original C+LOO and PB designs.

By comparing the $\kappa$ values of those DO-C+LOO designs and the original PB designs across the three simulated models (Figure S12), we reaffirmed the correspondence between the network structures and correlation patterns by reproducing results consistent with SI G. For reference, we also visualized the statistical power of the DO-C+LOO-based additive models (Figure S13).

In Model $\Phi$, where PB performed better than C+LOO, none of the DO-C+LOO designs outperformed the original PB designs across different Gaussian noise levels (Figure S12a). This suggests that experimental designs perform better in systems regulated by parallel cascades—as modeled in Model $\Phi$—when they become more D-optimal (i.e., less correlated).

On the other hand, in Model $\Psi$, the original PB designs performed the worst across all Gaussian noise conditions (Figure S12b), implying that the unique correlation patterns of the C+LOO design may have adapted to the complexity of interactions as emulated in Model $\Psi$.

In Model $\Lambda$, results consistent with Figure S10b were observed: C+LOO and DO-C+LOO designs outperformed PB under small Gaussian noise, whereas they showed relatively poor performance with large Gaussian noise (Figure S12c). Notably, DO-C+LOO did not show a monotonic increase in $\kappa$ scores with increasing $n_{\max}$, which further emphasizes the complex behavior of experimental design performance under Model $\Lambda$. Thus, the complexity of the correspondence between model configurations and $\kappa$ scores is illuminated, especially when the regulatory network is sparse.

These findings illustrate the difficulty of establishing a universal criterion for evaluating experimental designs independently of the network structures underlying target phenomena. Notably, the performance of DO-C+LOO designs did not exhibit a monotonic trend, even though D-optimization consistently reduced D-criterion values. This underscores the challenge of predicting optimal experimental designs solely based on design matrices, without accounting for background models. Although alternative optimization criteria exist—including the A-criterion [dABK+95]—they may face similar challenges. Consequently, these results support the importance of benchmarking experimental designs using simulated models.

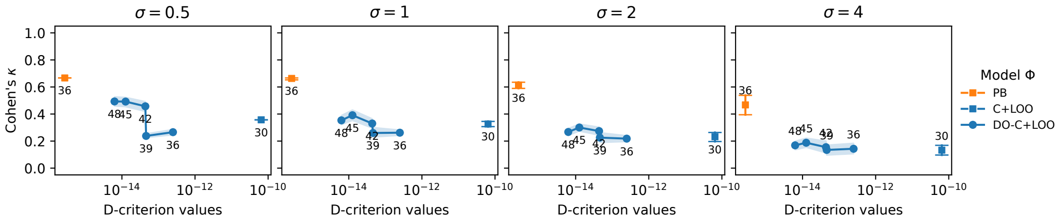


(a) Benchmarking (Model Φ)

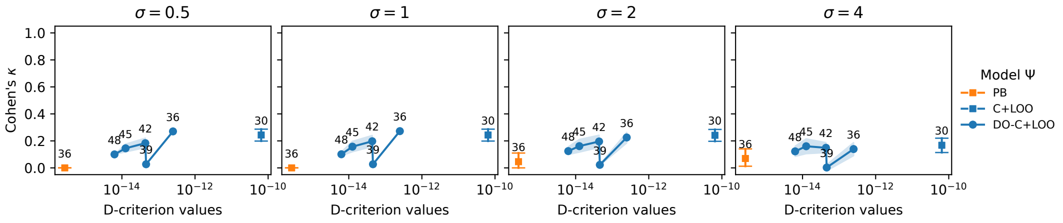


(b) Benchmarking (Model Ψ)

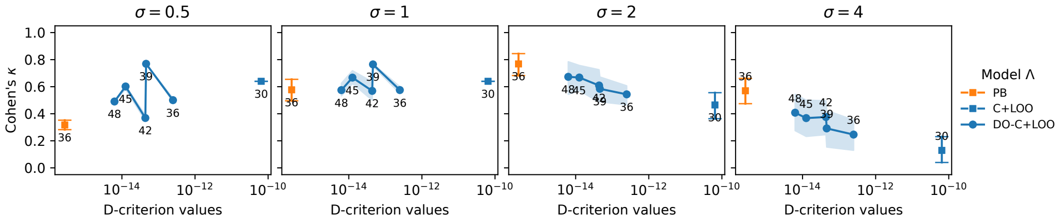


(c) Benchmarking (Model Λ)

**Figure S12: Performance comparison between the DO-C+LOO and PB designs**

**a–c**: Line charts showing $\kappa$ values for the original C+LOO (blue square), DO-C+LOO (blue circle), and PB (orange square) designs for nine factors at the replication level $N = 3$. The Gaussian noise terms $\varepsilon_{i,j}, \varepsilon_{y,j}$ were generated from the normal distributions of different standard deviations: $\sigma \in \{0.5, 1, 2, 4\}$. The markers show the mean of results from 30 different random seeds, and the shaded areas represent the 95% bootstrap CIs (the number of bootstraps $n_{\text{boot}} = 1000$). The numbers near the markers represent $n_{\max}$ values.

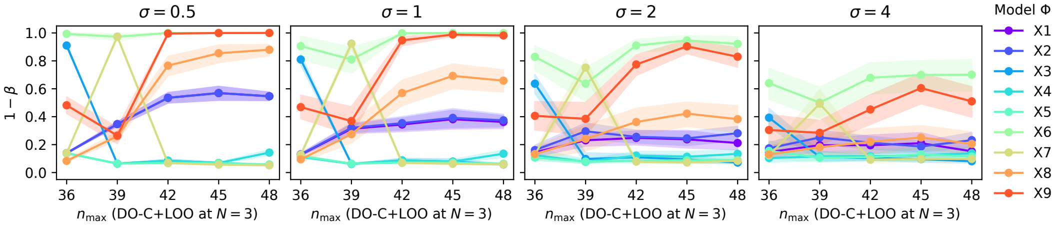


(a) Power analysis (Model Φ)

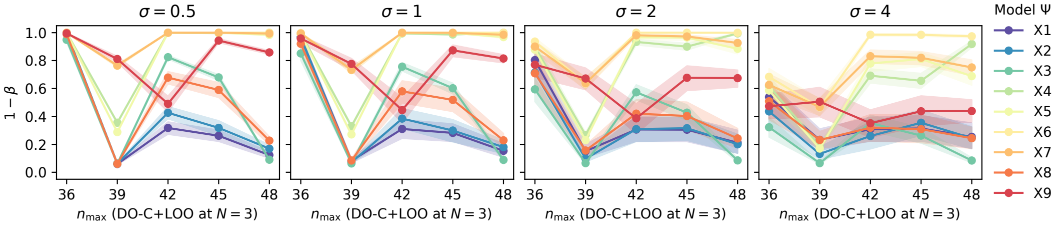


(b) Power analysis (Model Ψ)

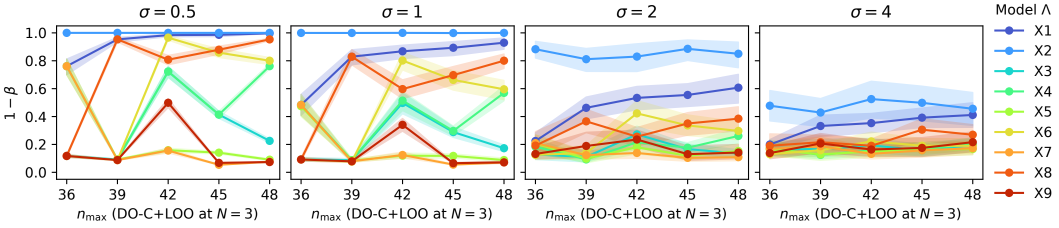


(c) Power analysis (Model Λ)

**Figure S13: Statistical power of the DO-C+LOO-based additive models**

**a–c**: Line charts showing $1 - \beta$ values for the DO-C+LOO-based additive models at various $n_{\max}$ values when the replication level was set to $N = 3$. The Gaussian noise terms $\varepsilon_{i,j}, \varepsilon_{y,j}$ were generated from the normal distributions of different standard deviations: $\sigma \in \{0.5, 1, 2, 4\}$. The markers show the mean of results from 30 different random seeds, and the shaded areas represent the 95% bootstrap CIs (the number of bootstraps $n_{\text{boot}} = 1000$).

### I PBSI

#### I.1 Identification of PBSI

To develop a simplified metric with improved predictive performance for the "PB-suited" class based on SHAP analysis, we calculated arithmetic means for all possible combinations of the top six features showing the highest AUROC scores in the ESM4 dataset—$1 - \mathrm{MaxRW}_{(+)}/\mathrm{R}^*$, $1 - \mathrm{RW}_{(+)}\%$, $1 - \mathrm{MaxRW}_{\&}/\mathrm{R}^*$, $\mathrm{P}_{(+)}\%$, $1 - \mathrm{R}^*_{(+)}/\mathrm{R}^*$, and P%—yielding 57 composite indices named "arithmetic1" through "arithmetic57" (Figure S14a).

We evaluated the predictive performance of each composite metric individually in the ESM4 dataset and found that "arithmetic12" achieved the highest AUROC (Figures S14b–S14c). This feature combination—comprising $\mathrm{P}_{(+)}\%$, $1 - \mathrm{MaxRW}_{(+)}/\mathrm{R}^*$, and $1 - \mathrm{RW}_{(+)}\%$ (Figure S14a)—was adopted as the PBSI.

Since PBSI is simply defined as the arithmetic mean of its components, we evaluated the effect of not optimizing the weights by performing logistic regression using the same features. The AUROC difference was minimal (Figure S14d), suggesting that PBSI performs comparably well even without fitting.

#### I.2 Scalability of PBSI

To validate the scalability of PBSI beyond four-factor networks, we developed the Exhaustive Search Model of nine factors (ESM9) framework. Given that the structural search space for nine factors expands to $3^{45}$ configurations, exhaustive enumeration is computationally intractable. We therefore randomly sampled 20,000 representative network structures to construct a validation dataset (Figures S15a–S15b). Using the same classification criteria as ESM4, we evaluated the predictive performance of PBSI (Figure S15b).

Despite the increased complexity, PBSI outperformed all other individual features (Figures S15c–S15d); while the predictive performance of individual features varied—with some retaining moderate discriminative power and others degrading to near-random levels—PBSI consistently achieved the highest ranking. Notably, PBSI achieved the highest average precision (AP) significantly outperforming all the other features and the baseline (which is approximately 0.62 according to the class balance displayed in Figure S15b), underscoring the robustness of predictive performance relative to the other features. Although the absolute performance suggests that perfect prediction is challenging when estimating appropriate designs from highly diverse GRN structures like ESM9 using simple metrics, these results demonstrate that PBSI remains the most robust and scalable indicator available.

To further examine the scalability of PBSI in comparison with other composite indices, we constructed ROC curves in the ESM9 dataset as well (Figure S16a). Among the indices whose rankings improved in terms of AUROC—such as "arithmetic41," "arithmetic43," and "arithmetic45," which were not within the top five in the ESM4 AUROC scores—PBSI ("arithmetic12") maintained the highest position (Figure S16b). Although its AUROC decreased in the ESM9 dataset relative to the ESM4 dataset, PBSI consistently exhibited the best predictive performance, thereby outperforming all other composite indices.

As with the ESM4 dataset, we also constructed logistic regression models for the ESM9 dataset to compare the predictive performance of PBSI—defined as a simple arithmetic mean—with that of the fitted models in which feature weights were optimized. PBSI exhibited nearly identical AUROC performance to the logistic regression model (Figure S16c), suggesting its utility as a model-free metric, even when applied to systems with a different number of factors from which it was originally derived.

#### I.3 Explainability of PBSI: validation via representative topological features

The PBSI was designed to be interpretable, with components characterizing specific network topologies: high values imply parallel cascades with minimal interaction, while low values imply complex, converging interactions. While these tendencies are consistent with the empirical hypotheses from our initial benchmarks (Models $\Phi$ and $\Psi$), verifying whether these topological definitions truly drive design suitability requires a rigorous validation process.

Accordingly, we adopted a two-step approach:

1. **Calibration and *sanity check***: Determining decision thresholds using the ESM9 dataset and verifying them with the original benchmark models (Models $\Phi$ and $\Psi$) are correctly classified.

2. **Structural validation**: The core verification step. We tested whether the topological archetypes defined by PBSI—specifically, parallel vs. entangled structures—truly represent the PB- and C+LOO-suited toplogies. To do this, we introduced new simulators (Models $\Pi$, $\Delta$, and $\Sigma$ within the ESM9 architecture) explicitly constructed to embody these archetypes.

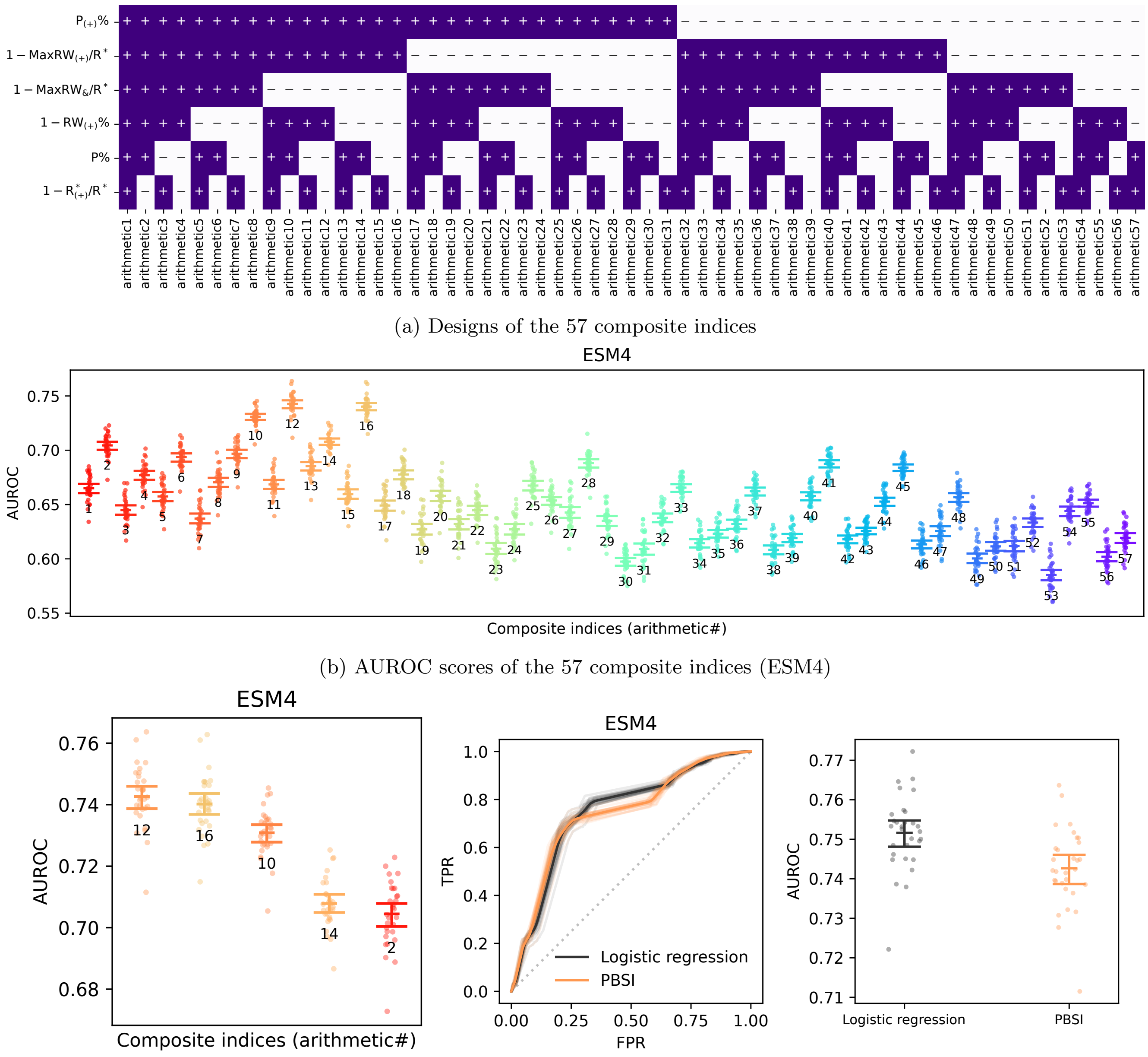


(a) Designs of the 57 composite indices

(b) AUROC scores of the 57 composite indices (ESM4)

(c) The top five indices (ESM4)

(d) PBSI vs. logistic regression (ESM4)

**Figure S14: An exhaustive search for a better index at the "PB-suited" class prediction**

**a**: Structures of the 57 composite indices (+, inclusion of features; −, absence of features). **b**–**c**: (b) AUROC scores of all 57 and (c) the top five indices with the highest mean AUROC in the ESM4 dataset. Plotting raw AUROC scores from each fold, the error bars show the mean and 95% bootstrap CIs. The numbers adjacent to the plots indicate the serial numbers of the composite indices. The"arithmetic12", which achieved the highest AUROC, was adopted as the PBSI. **d**: ROC curves (left) and AUROC scores (right) of the logistic regression model (black) and PBSI (orange). For the ROC curves, semi-transparent lines show the raw curves for each of the 30 cross-validation folds, overlaid with the mean curve. For the AUROC plots, each marker corresponds to raw data from an individual fold, and the mean and 95% bootstrap CIs are directly annotated on the plot in addition to the error bars.

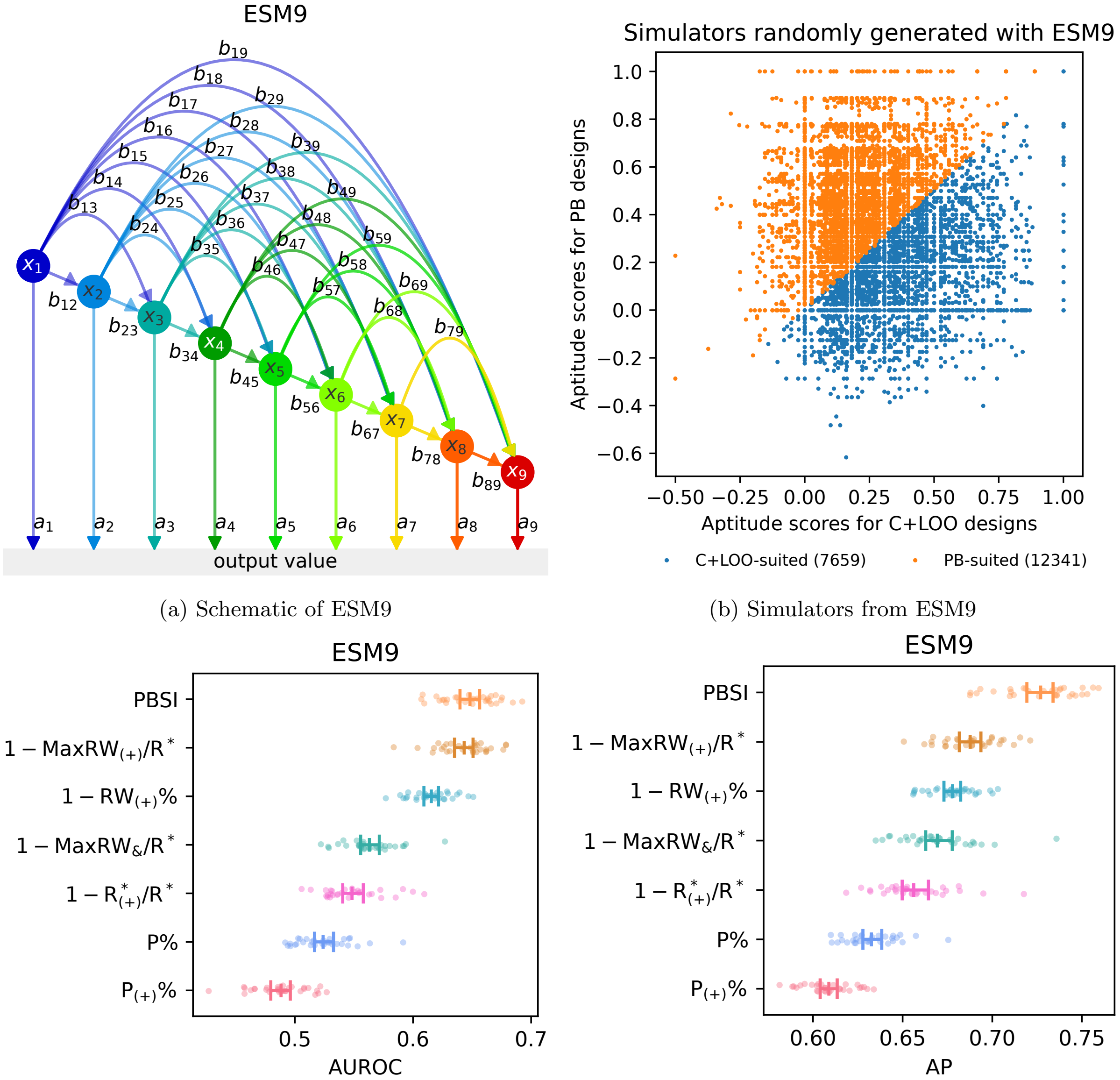


(a) Schematic of ESM9

(b) Simulators from ESM9

(c) AUROC (ESM9)

(d) AP (ESM9)

**Figure S15: PBSI is proposed as a simple, scalable, parameter-free metric recommending replacement of the C+LOO design with the PB design**

**a**: Example network structure generated from ESM9. **b**: Classification of the 20,000 randomly sampled ESM9 simulators. To reduce computational costs, representative network structures were randomly sampled rather than exhaustively enumerated. Using aptitude scores for C+LOO ($\kappa^*_{\mathrm{C}}$) and PB ($\kappa^*_{\mathrm{P}}$), simulators were binned into "C+LOO-suited" (blue; $\kappa^*_{\mathrm{C}} \geq \kappa^*_{\mathrm{P}}$) and "PB-suited" (orange; $\kappa^*_{\mathrm{C}} < \kappa^*_{\mathrm{P}}$) classes. The numbers in parentheses indicate the number of simulators in each category. **c**–**d**: AUROC (c) and AP (d) curves of PBSI and the top six features with the highest AUROC scores in Figure 5e. The error bars indicate the mean and 95% bootstrap CIs.

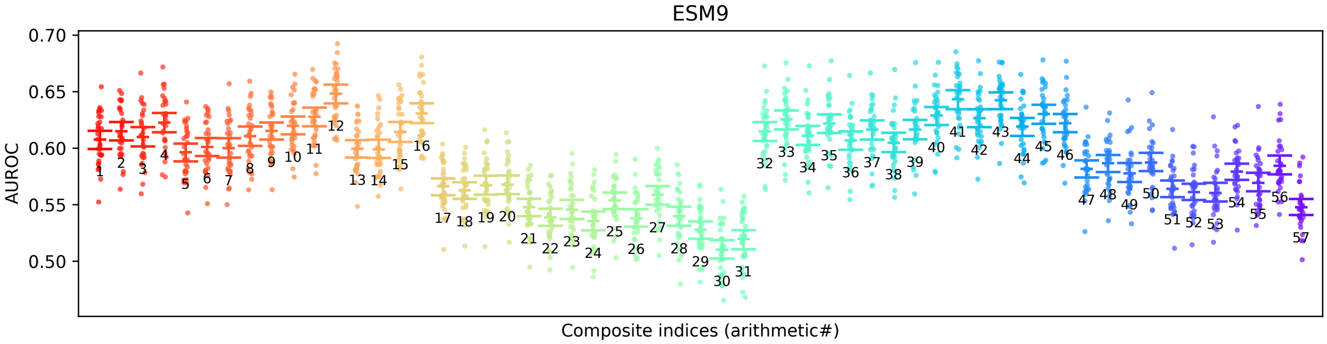


(a) AUROC scores of the 57 composite indices (ESM9)

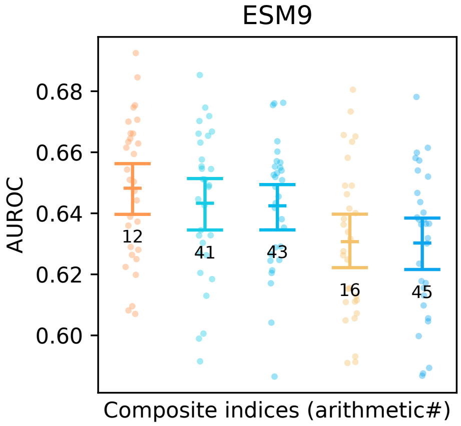


(b) The top five indices (ESM9)

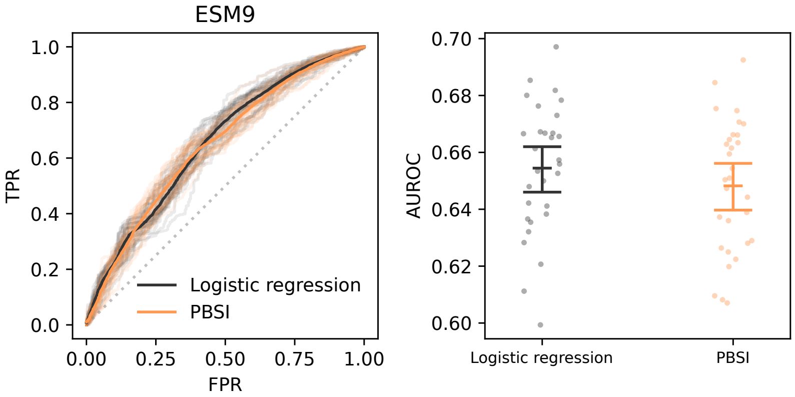


(c) PBSI vs. logistic regression (ESM9)

**Figure S16: PBSI (arithmetic12) outperformed other composite indices for randomly sampled ESM9 models**

**a–b**: AUROC scores of all 57 (a) and the top five indices (b) with the highest mean AUROC in the ESM9 dataset. Plotting raw AUROC scores from each fold, the error bars show the mean and 95% bootstrap CIs. The numbers adjacent to the plots indicate the serial numbers of the composite indices. The PBSI (“arithmetic12”) outperformed all other indices, consistent with the results from the ESM4 datasets. **c**: ROC curves (left) and AUROC scores (right) of the logistic regression model (black) and PBSI (orange). For the ROC curves, semi-transparent lines show the raw curves for each of the 30 cross-validation folds, overlaid with the mean curve. For the AUROC plots, each marker corresponds to raw data from an individual fold, and the mean and 95% bootstrap CIs are directly annotated on the plot in addition to the error bars.

By utilizing nine-factor based models, we also provided additional validation of the scalability of PBSI, which was originally constructed on four-factor networks.

##### I.3.1 Threshold calibration and sanity check using benchmark models

First, to establish a quantitative criterion, we determined optimal PBSI thresholds using the ESM9 dataset. Thresholds maximizing accuracy, $F_1$ score, and TPR−FPR were identified (Figure S17a), collectively suggesting a practical threshold of approximately 0.6.

As a *sanity check*, we applied these thresholds to Models Φ and Ψ (Figure S17b). Model Λ was excluded due to its inconsistent design preference (SI G.3). While the $F_1$-based threshold proved conservative, the thresholds for accuracy and TPR − FPR successfully recommended the optimal designs consistent with our simulation results; specifically, they correctly classified Model Φ as PB-suited and Model Ψ as C+LOO-suited (Figures 3d–3e).

This confirmed that the metric reproduces the expected behavior on the models from which our empirical intuition was derived.

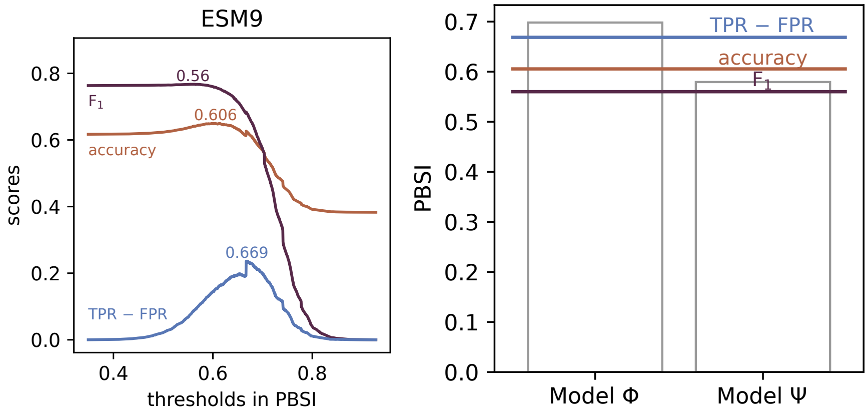


(a) Best thresholds (ESM9) (b) PBSI (Models Φ and Ψ)

**Figure S17: Threshold calibration and sanity check using Models Φ and Ψ**

**a**: To determine the optimal threshold of PBSI, accuracy (tangerine), $F_1$ (plum), and TPR − FPR (blue) scores were calculated for each Plackett–Burman suitability index value. The numbers along the curves indicate the argmax values for each performance metric. **b**: PBSI scores for Models Φ and Ψ together with the thresholds for accuracy (tangerine), $F_1$ (plum), and TPR − FPR (blue) fine-tuned in Figure S17a. Model Λ, which consisted of only two active factors, was excluded from this evaluation because no consistent trend was observed in the relative performance between the C+LOO and PB designs (see Results 2.2 and SI G.3 for details).

##### I.3.2 Structural validation: verifying the topological archetypes of design suitability

To verify the consistency of our structural hypothesis—that "parallel" and "entangled" topologies are indeed the archetypes of PB and C+LOO suitability—we performed an inverse test. We introduced three simulators based on the ESM9 architecture, Models Π, Δ, and Σ, explicitly designed to yield distinctively high or low PBSI scores based on these topological definitions (Methods 4.8.8).

- **Model Π (high PBSI archetype)**: Designed to have PBSI = 1, representing a pure parallel cascade topology (Figures S18a–S18b). Benchmarks confirmed the PB design's superiority (Figures S18d–S18e), demonstrating that the parallel structure captured by high PBSI is indeed a key driver of PB suitability.
- **Model Δ (low PBSI archetype)**: Designed to yield a low PBSI value, representing a pure entangled topology (Figures S19a–S19b). Benchmarks showed the C+LOO design's superiority (Figures S19d–S19e), confirming that the dense convergence of positive gene–gene interactions captured by low PBSI necessitates the robustness of C+LOO.
- **Model Σ (another high PBSI archetype)**: Designed to yield a high PBSI value despite featuring complex upstream positive regulations (structurally entangled). Crucially, the terminal regulation is negative, which

effectively suppresses $\text{MaxRW}_{(+)}/\text{R}^*$ and $\text{RW}_{(+)}\%$ (Figures S20a–S20b). Benchmarks confirmed the slight superiority of PB (Figures S20d–S20e), demonstrating that complex gene–gene interactions do not severely compromise PB suitability provided the ultimate regulatory effect is downregulation.

By confirming that these intentionally constructed archetypes yield the predicted design preferences, we validated the logical consistency between the PBSI components and the resulting experimental design suitability.

Interestingly, Model Π and Model Δ exhibited main effects differing by orders of magnitude ($10^3$) despite using identical parameters, solely due to differences in GRN structure (Figures S18c and S19c). Given that gene expression values (both real and simulated) are non-negative, amplification typically occurs via chains of upregulation, leading to extremely high main effect values. Therefore, we identified the entangled topology composed of chains of positive interaction as the primary driver of "signal amplification."

In contrast, Model Σ, which shares identical entangled upstream regulations with Model Δ, exerts strong downregulation via the terminal negative pathway from X9 (Figure S20c). Despite its structural complexity, Model Σ maintains a high PBSI and demonstrates the relative superiority of the PB design, qualifying it as a "PB-suited" topology. This finding implies more than a mere artificial manipulation of PBSI via a terminal negative pathway; it suggests that under the non-negativity constraint of gene expression data, complex gene–gene interactions do not induce strong nonlinear effects (i.e., signal amplification) unless they result in upregulation via positive pathways. Indeed, while the regulations upstream of X9 are identical between Models Δ and Σ, the magnitudes of their main effects differ by $10^2$. This confirms that without direct upregulation of the final readout, structural entanglement alone does not generate the type of signal amplification that necessitates C+LOO designs.

##### I.3.3 Theoretical insight: model misspecification under heteroscedasticity

Our analysis revealed that while the PB design is robust across most network topologies, the C+LOO design exhibits relatively high performance specifically under signal amplification. Here, we attempt a unified explanation for the performance trade-off between PB and C+LOO from the perspective of how design-specific correlation structures influence estimation within the framework of multivariate analysis (e.g., ANOVA/MLR).

As illustrated in Figure 1b and Figure S2c, both PB and C+LOO designs possess distinct correlation structures where interaction terms—variables not explicitly included in the additive model (Methods 4.4.2)—are partially confounded with the main effects. When computing the Ordinary Least Squares (OLS) estimator using the orthogonal projection matrix $P(\mathbf{X}) := (\mathbf{X}^\top\mathbf{X})^{-1}\mathbf{X}^\top$ (where $\mathbf{X}$ denotes the model matrix; refer to Equations (60) and (67) for details), these design-specific correlations cause the influence of unmodeled interaction terms to propagate into the main effect estimates. In this context, the PB design maintains near-orthogonality, whereas the C+LOO design inherently contains a more specific, structured correlation pattern.

Standard methods like ANOVA and MLR assume homoscedasticity (constant variance). However, real biological systems can exist on a spectrum between perfect homoscedasticity and severe heteroscedasticity. The validity of the design comparison depends on where the system lies on this spectrum. In systems approximating homoscedasticity (e.g., Model Π; see also Figure S21a), the Gauss–Markov theorem holds [SL03], rendering the OLS estimator the Best Linear Unbiased Estimator (BLUE). Under these conditions, "external" correlation structures are detrimental to accurate estimation. Therefore, the PB design, which minimizes structural confounding and maintains orthogonality, provides superior effect estimation compared to C+LOO.

Conversely, in systems with clear heteroscedasticity, the Gauss–Markov theorem does not hold, and the OLS estimator is no longer BLUE. Consequently, applying general linear models (including Dunnett's test, MLR, and ANOVA) constitutes a model misspecification, meaning the validity of the model's conclusions (e.g., $p$-values and confidence intervals) is not guaranteed [SL03]. In this regime, estimators are still subject to the propagation of unmodeled effects via design-specific correlations. However, if the design possesses a correlation structure that coincidentally aligns with the true network topology, this "confounding" can act as a heuristic correction. This can result in a higher sign consistency with the ground truth main effects (i.e., higher Weighted Cohen's $\kappa$) compared to orthogonal designs, despite the theoretical violation of linear assumptions.

In summary, comparing PB and C+LOO designs in heteroscedastic systems requires evaluating which correlation structure is empirically beneficial for the specific case. Our exhaustive search using ESM4 and ESM9 confirmed that the PB design is robust for most topologies. However, in the exceptional case of signal amplification, the correlation structure of the C+LOO design better reflected the system's structure. Signal amplification generates significant heteroscedasticity as noise is amplified at each node (e.g., Model Δ; see also Figure S21b). Under such model misspecification, the C+LOO design, which retains high correlations such as $\rho_{i\times j,i}$, acted as a "niche adaptation" that matched the distorted signal structure more effectively than the PB design, which enforces separation ($\rho_{i\times j,i} = 0$).

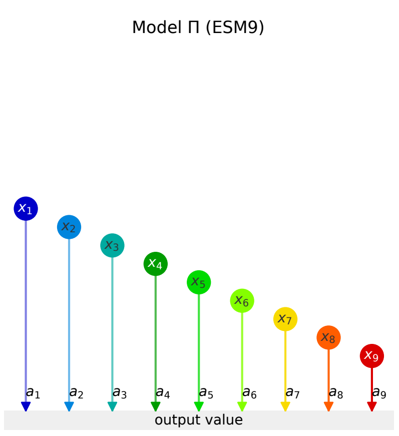


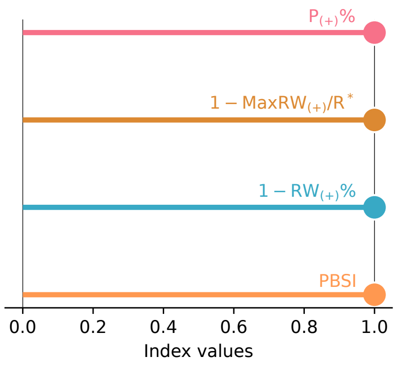


(a) Model Π (b) Index scores

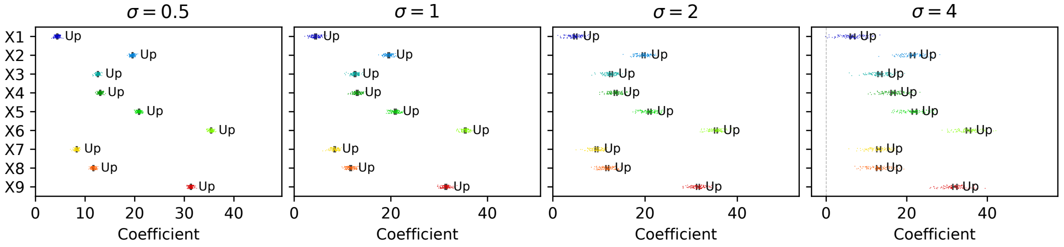


(c) Ground truth main effects

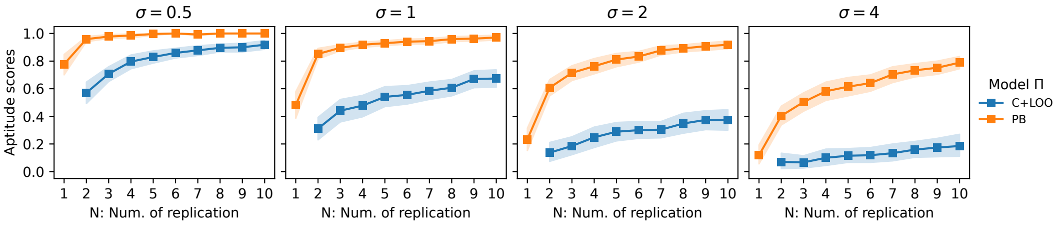


(d) Benchmarking

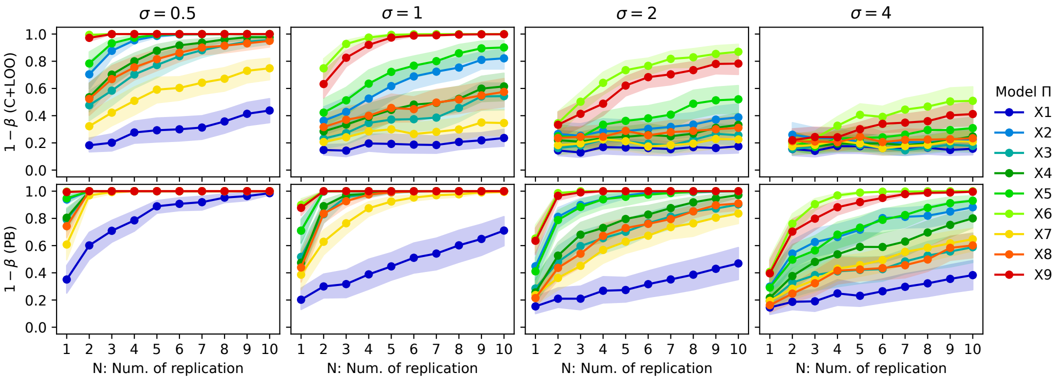


(e) Power analysis

**Figure S18: Benchmarking experimental designs with Model Π**

**a**: Schematic of Model Π. **b**: $\mathrm{P}_{(+)}\%$, $1-\mathrm{MaxRW}_{(+)}/\mathrm{R}^{*}$, $1-\mathrm{RW}_{(+)}\%$, and PBSI scores of Model Π. **c**: Main effects of X1–X9 in Model Π estimated from the FF-based simulations according to the definition of the main effect. The plots represent 100 different results generated with distinct random seeds and the error bars show the 95% bootstrap CIs (the number of bootstraps $n_{\mathrm{boot}} = 10000$). Based on the CIs, we assigned interpretations (Up, N.S., or Down). **d**: Line charts showing $\kappa$ values for the C+LOO (blue) and PB (orange) designs at replication levels $1 \leq N \leq 10$. The Gaussian noise terms $\varepsilon_{i,j}, \varepsilon_{y,j}$ were generated from the normal distributions of different standard deviations: $\sigma \in \{0.5, 1, 2, 4\}$. **e**: The statistical power $1-\beta$ of ANOVA for the C+LOO (top panels) and PB (bottom panels) designs at $\alpha = 0.05$. In Figures S18d–S18e, the markers show the mean of results from 30 different random seeds, and the shaded areas represent the 95% bootstrap CIs ($n_{\mathrm{boot}} = 1000$).

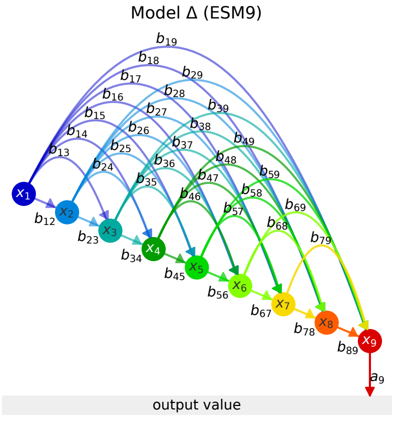


(a) Model $\Delta$

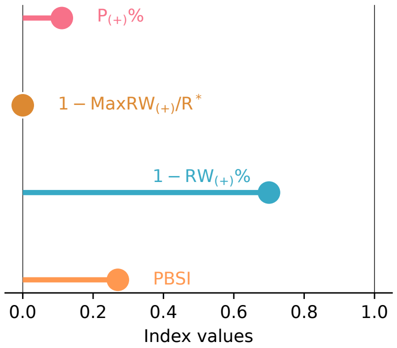


(b) Index scores

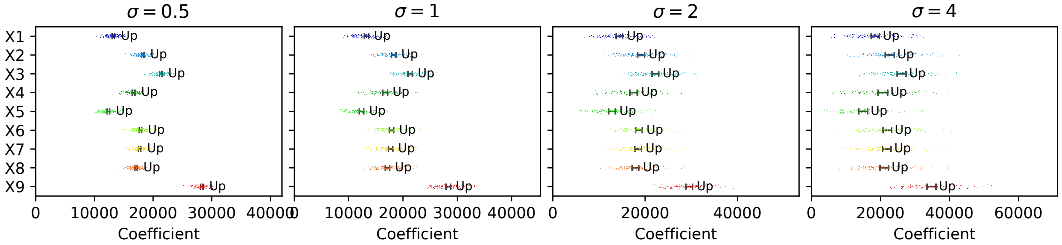


(c) Ground truth main effects

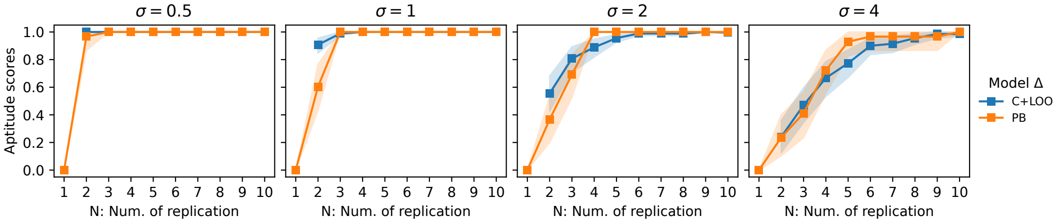


(d) Benchmarking

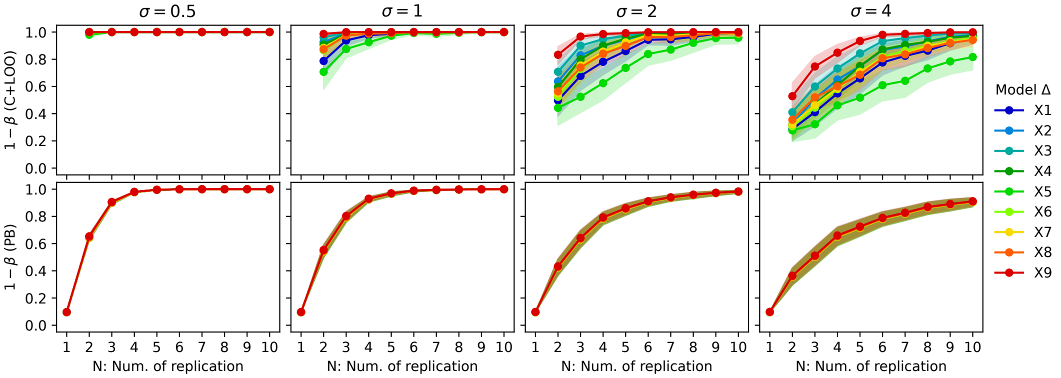


(e) Power analysis

**Figure S19: Benchmarking experimental designs with Model $\Delta$**

**a**: Schematic of Model $\Delta$. **b**: $\mathrm{P}_{(+)}\%$, $1 - \mathrm{MaxRW}_{(+)}/\mathrm{R}^*$, $1 - \mathrm{RW}_{(+)}\%$, and PBSI scores of Model $\Delta$. **c**: Main effects of X1–X9 in Model $\Delta$ estimated from the FF-based simulations according to the definition of the main effect. The plots represent 100 different results generated with distinct random seeds and the error bars show the 95% bootstrap CIs (the number of bootstraps $n_{\mathrm{boot}} = 10000$). Based on the CIs, we assigned interpretations (Up, N.S., or Down). **d**: Line charts showing $\kappa$ values for the C+LOO (blue) and PB (orange) designs at replication levels $1 \leq N \leq 10$. The Gaussian noise terms $\varepsilon_{i,j}, \varepsilon_{y,j}$ were generated from the normal distributions of different standard deviations: $\sigma \in \{0.5, 1, 2, 4\}$. **e**: The statistical power $1 - \beta$ of ANOVA for the C+LOO (top panels) and PB (bottom panels) designs at $\alpha = 0.05$. In Figures S19d–S19e, the markers show the mean of results from 30 different random seeds, and the shaded areas represent the 95% bootstrap CIs ($n_{\mathrm{boot}} = 1000$).

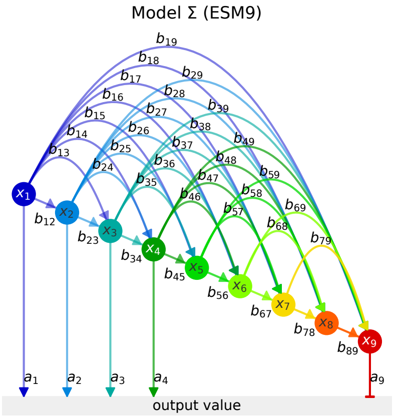


(a) Model Σ

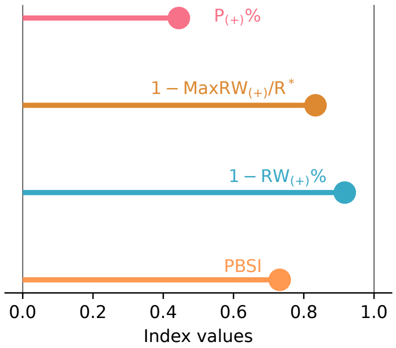


(b) Index scores

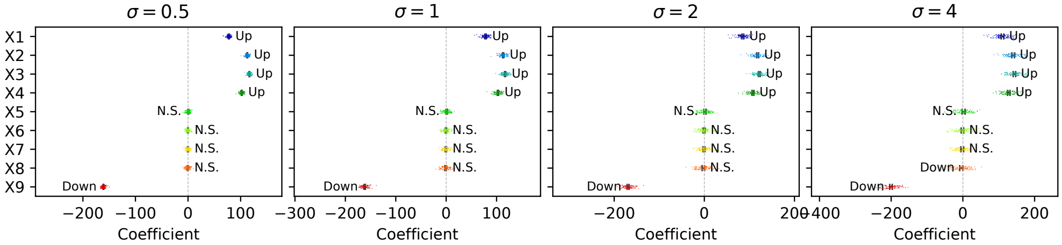


(c) Ground truth main effects

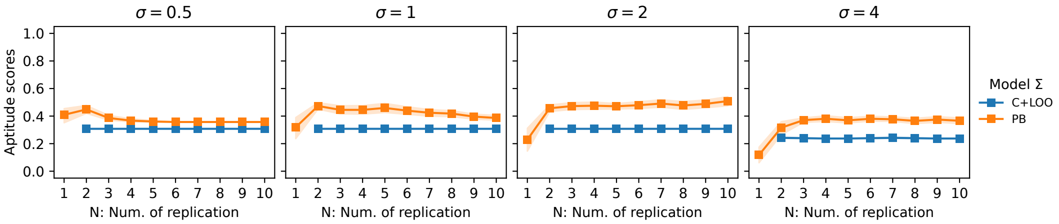


(d) Benchmarking

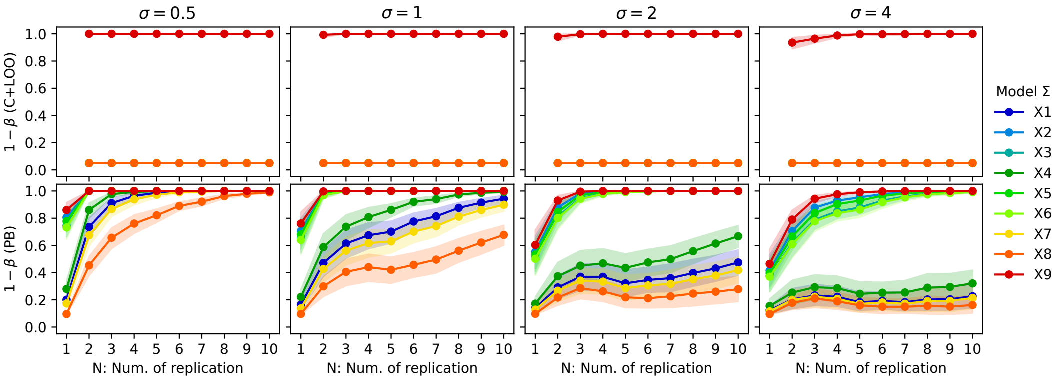


(e) Power analysis

**Figure S20: Benchmarking experimental designs with Model Σ**

**a**: Schematic of Model Σ. **b**: $\mathrm{P}_{(+)}\%$, $1-\mathrm{MaxRW}_{(+)}/\mathrm{R}^*$, $1-\mathrm{RW}_{(+)}\%$, and PBSI scores of Model Π. **c**: Main effects of X1–X9 in Model Π estimated from the FF-based simulations according to the definition of the main effect. The plots represent 100 different results generated with distinct random seeds and the error bars show the 95% bootstrap CIs (the number of bootstraps $n_{\mathrm{boot}} = 10000$). Based on the CIs, we assigned interpretations (Up, N.S., or Down). **d**: Line charts showing $\kappa$ values for the C+LOO (blue) and PB (orange) designs at replication levels $1 \le N \le 10$. The Gaussian noise terms $\varepsilon_{i,j}, \varepsilon_{y,j}$ were generated from the normal distributions of different standard deviations: $\sigma \in \{0.5, 1, 2, 4\}$. **e**: The statistical power $1-\beta$ of ANOVA for the C+LOO (top panels) and PB (bottom panels) designs at $\alpha = 0.05$. In Figures S20d–S20e, the markers show the mean of results from 30 different random seeds, and the shaded areas represent the 95% bootstrap CIs ($n_{\mathrm{boot}} = 1000$).

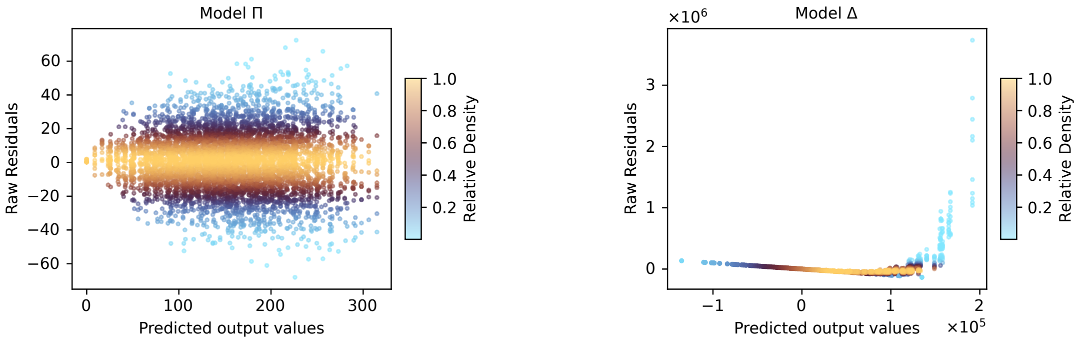


(a) Homoscedastic system (Model Π) (b) Heteroscedastic system (Model Δ)

**Figure S21: Homoscedasticity vs. heteroscedasticity represented in residual plots**

**a–b**: Residual density plots for Models Π (a) and Δ (b) under FF designs ($N = 10, \sigma = 1$). The horizontal axis represents the predicted output values $\hat{y}$ calculated as the product of the model matrix $\mathbf{X}_\Omega$ and the coefficient vector $\hat{\boldsymbol{\beta}} := [\hat{\mu}, \hat{M}^1, \ldots, \hat{M}^9]^\top$. Here, $\hat{\mu}$ denotes the global mean computed from FF simulations ($N = 100, \sigma = 1$), and $\hat{M}^1, \ldots, \hat{M}^9$ denote the main effects averaged over 100 independent FF simulations ($N = 1, \sigma = 1$). The vertical axis displays the raw residuals ($y - \hat{y}$). Points are colored by "relative density," defined as the Gaussian kernel density of the raw residuals normalized by the maximum density value. Model Π exhibits an almost homoscedastic distribution (horizontal band), whereas Model Δ shows severe structural heteroscedasticity (variance expansion).

##### I.3.4 Summary of topological interpretation

The validation of PBSI, which can be computed solely from network features, strongly suggests that network structure plays a critical role in determining the suitability of experimental designs. Consistent with the empirical assumptions derived from Models Φ and Ψ, as well as the inferences from the three PBSI components and the archetypal Models Π, Δ, and Σ, the following structural interpretation of PBSI is supported:

- **High PBSI**: While a parallel structure is a quintessential example of topological features favoring PB designs, high PBSI is fundamentally characterized by the *absence* of structural features that necessitate C+LOO (i.e., signal amplification). As demonstrated by Model Σ, even structurally complex networks can exhibit high PBSI and PB suitability if they lack the positive edges connecting factors to the output value, which are required for amplification. Thus, the most parsimonious interpretation is that the PB design serves as the default superior strategy, with C+LOO becoming advantageous only under specific exceptional conditions.
- **Low PBSI**: Rather than entanglement *per se*, a specific type of entanglement—characterized by cascades involving unbroken chains of positive regulations that must explicitly culminate in positive terminal edges to the output—is a reliable topological indicator of C+LOO preference. Crucially, as highlighted by the contrast between Models Δ and Σ, even dense positive upstream cascades do not trigger the "signal amplification" necessitating C+LOO designs if the final link to the output is negative.

Collectively, these results demonstrate that PBSI functions as a scalable and explainable predictor of PB suitability even in nine-factor networks, providing a solid theoretical basis for its application in real-world scenarios.

Ultimately, this implies that the biological phenomenon of "signal amplification" serves as the physical manifestation of structural noise inflation (heteroscedasticity) that violates the fundamental assumption of linear screening designs. Consequently, unless such specific amplification structures are present, the orthogonality of PB designs remains robust, supporting their broad applicability in transcriptomic studies.

#### I.4 Biological relevance of PBSI-related GRN topologies

The partial validation of PBSI's effectiveness, computed solely from network features, suggests that network structure plays a critical role in determining the suitability of experimental designs. Ultimately, the discussion regarding topology converges on whether the phenomenon of "signal amplification" renders the orthogonal PB design suboptimal.

PBSI includes $1 - \text{MaxRW}_{(+)}/\text{R}^*$ and $1 - \text{RW}_{(+)}\%$ as its components, suggesting that PB designs can outperform C+LOO designs in systems characterized by parallel GRN topologies. Such scenarios are highly realistic in practice.

For instance, a genome-wide Perturb-seq study demonstrated that gene functions often cluster into distinct modules with high intra-module correlation but minimal inter-module correlation, as observed in the distinct functional clusters of the Integrator complex [RSP+22]. Although that study was genome-wide, if a focused Perturb-seq experiment were designed to target genes across such distinct, non-interacting modules, the underlying GRN would functionally resemble a set of parallel cascades—indicating a "PB-suited" landscape. In such cases, our findings suggest that the PB design would be statistically superior to C+LOO. Furthermore, as validated in this study, the PB design exhibits relative superiority in any system where the "signal amplification" that undermines OLS-based frameworks is absent.

Conversely, the C+LOO design remains superior to PB designs under mechanisms involving chains of positive regulations that amplify biological signals. In a general biological context, MAPK signaling [ZL02] serves as a representative example of signal amplification via such cascades. A mathematical analysis of MAPK signaling based on Boolean networks [KK22] reported that kinases located further downstream in protein-level interactions tend to exhibit higher activity. This mirrors the phenomenon observed in our Model $\Delta$, where X9, positioned at the bottom of a complex chain of positive regulations, exhibits the strongest positive main effect (Figure S19c).

However, it is crucial to examine whether these interpretations hold in real scRNA-seq data. Since the MAPK example primarily involves regulation at the level of protein phosphorylation, it is necessary to assess whether similar structural interpretations are transferable to transcriptomic data. Furthermore, it is essential to confirm the consistency of the hypothesis derived from our simulations: that even in the presence of complex interactions, the resulting signal amplification is minimal if the chains of positive edges are interrupted. While comparing experimental designs directly in the context of Perturb-seq remains challenging as discussed in SI D.2, PBSI can be defined for any GRN, including those inferred from scRNA-seq data. Therefore, to further elaborate on the biological relevance of PBSI and GRN topology, we examined whether the GRN structures inferred from scRNA-seq data for specific biological phenomena align with the trends in PBSI values.

Assuming a preliminary experiment phase where experimental designs are evaluated prior to Perturb-seq, we inferred GRNs from plain scRNA-seq data without perturbations. To ensure realistic conditions such as sequencing depth, we extracted a sub-dataset corresponding to the non-targeting (control) condition from the genome-wide K562 Perturb-seq dataset acquired by Replogle et al. (2022) [RSP+22]. Using this real-world dataset for GRN inference allowed us to mimic the sample and library sizes typical of a realistic preliminary experiment for Perturb-seq.

Given that Perturb-seq is designed to investigate the correspondence between interventions and phenotypes (SI D.1), we constructed GRNs to examine the impact of specific biological contexts—namely TNF signaling, Myc response, apoptosis, and the G2/M phase—on the S-phase score (Figure 6a) as a concrete example. We then evaluated PBSI and its constituent network indices for these inferred networks. Note that the GRNs in this article are treated as DAGs with Boolean-network-like edges, consisting of target genes and output values (corresponding to a single marker gene expression level or a phenotypic score), consistent with our minimalist simulator architecture. To infer such DAGs from scRNA-seq data, which include nodes like the S-phase score that are not explicitly present as gene features, we employed Bayesian network-based causal discovery. The signs of the edges were subsequently inferred from correlation coefficients. An overview of this workflow is illustrated in Figure S22.

In this section, we first detail the construction of GRNs favoring PB or C+LOO designs, providing supplementary information to Figure 6, followed by the presentation of additional concrete examples.

##### I.4.1 Examples of PB-suited GRN topologies favoring orthogonality

In this section, we examine TNF signaling and apoptosis as concrete examples of parallel GRN topologies. Given that high-PBSI topologies encompass a broad spectrum defined essentially by the absence of the exceptional conditions necessitating C+LOO, it is difficult to isolate a unique structural signature for intentional construction. Therefore, we aimed to construct a parallel topology from the data as the most representative archetype of PB suitability. It is important to note that the term "parallel GRN" here refers specifically to the structural relationship between the selected target genes and their impact on the S-phase score. The topology is determined by the choice of target genes and output values, rather than being an intrinsic property of the biological context itself. However, to avoid purely theoretical or artificial examples, we selected biological phenomena where a parallel interpretation is consistent with general biological knowledge.

We first consider the impact of TNF signaling on the S-phase score. Calculating the absolute Pearson correlations for genes listed in the `TNF-alpha signaling via NF-kB` category reveals a heatmap with a distinct block-diagonal structure: clusters of highly correlated genes form "red boxes" along the diagonal, surrounded by broad areas of weak correlation ("blue area") (Figure S23a). This pattern suggests the potential for modeling the system with a parallel mechanistic diagram. This interpretation is biologically plausible, given that TNF-$\alpha$ and NF-$\kappa$B

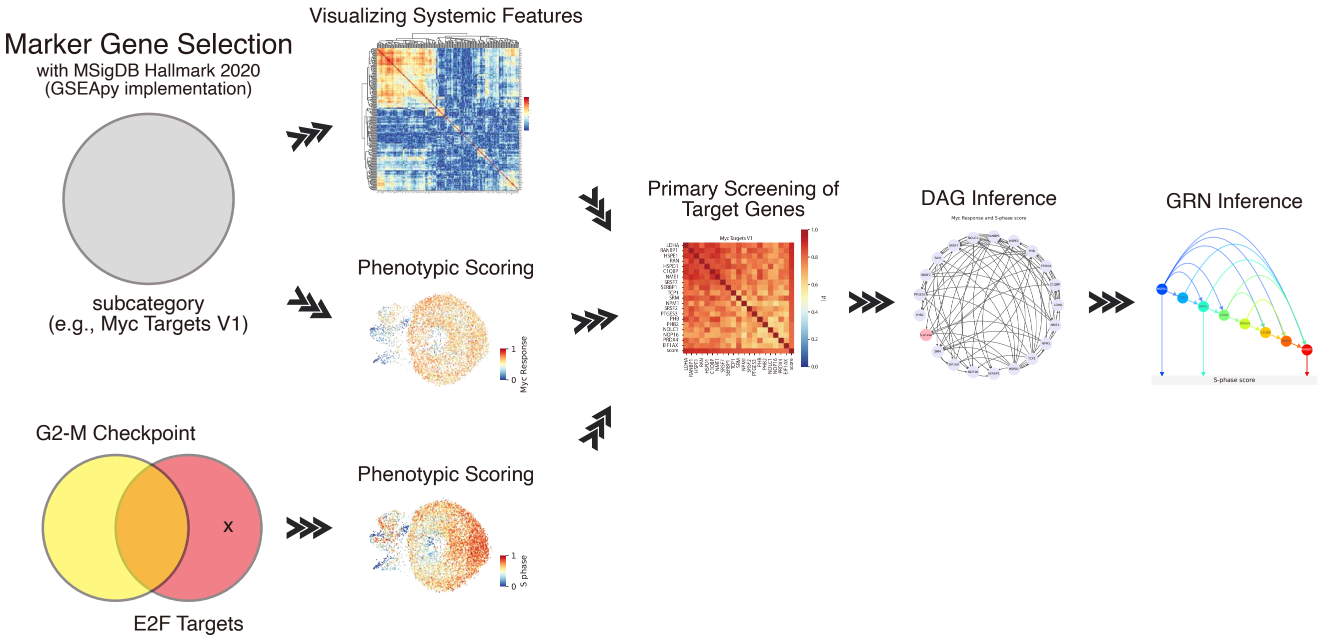


**Figure S22: Outline of the GRN inference workflow employed in this study**

The workflow consists of the following steps: **Marker gene selection**: Gene sets were retrieved from the MSigDB Hallmark 2020 collection [LBT+15] via `GSEApy` [FLP23]. **Phenotypic scoring**: Phenotypic scores were computed based on the selected genes. Specifically for the S-phase score, genes from the "E2F Targets" category were used, explicitly excluding those overlapping with the "G2-M Checkpoint" category to ensure phase specificity. **Visualizing systemic features**: Gene–gene correlations were visualized using MetaCell-aggregated pseudobulk data. This step aims to characterize the underlying biological context (e.g., "Myc Targets V1"), analogous to formulating experimental hypotheses or organizing domain knowledge in actual Perturb-seq studies. Since the target GRN constitutes an *induced subgraph* of the global GRN within a specific cellular state [OKO23], the selection of target genes is critical for determining whether the resulting topology is parallel or entangled. Thus, this visualization provides cues on whether to select highly independent or highly correlated gene sets. **Primary screening of target genes**: For demonstration purposes—rather than investigating a specific biological phenomenon based on *a priori* hypotheses—target genes were screened in a data-driven manner. Based on the systemic features observed above, we screened for genes exhibiting either low or high mutual correlations. The phenotypic score served as a reference to assess the relationship between individual genes and the systemic context (see Methods 4.9.3). **DAG inference**: A DAG comprising the screened genes and the S-phase score was inferred using Bayesian network-based causal discovery. Edges represent statistical dependencies identified at the pseudobulk level. **GRN inference**: The final GRN was constructed by extracting the induced subgraph where the S-phase score serves as the terminal (sink) node; genes remaining in this graph were defined as the final candidates. Edge signs were assigned based on the sign of the Pearson correlation coefficient between each gene and the S-phase score. The PBSI of the resulting GRN was then evaluated. Refer to SI I.4.3 for detailed considerations on the GRN inference logic.

function as central hubs regulating a diverse array of genes related to inflammation and cell survival [HG08]. Furthermore, since these factors are reported to regulate the S-phase and cell cycle progression through multiple distinct pathways [VAS94, HKE+99], the construction of a parallel GRN was anticipated.

To explicitly construct a parallel GRN, we selected the top 20 genes with the lowest mutual correlations using Recursive Feature Elimination with LASSO regression (see Methods 4.9.3 for details). The absolute Pearson correlations between these selected genes and the phenotypic score (Figure 6b)—calculated from all genes in the `TNF-alpha signaling via NF-kB` list—are shown in Figure 6d. We then inferred a DAG for these 20 genes and the S-phase score (Figure S23b). The resulting GRN topology exhibited fragmented cascades and a high PBSI score (Figures 6f–6g). Interestingly, while the inferred GRN contains some complex chains of positive gene–gene interactions, the edges directed toward the S-phase score (referred to as pathways in this study) are exclusively negative. This structural configuration prevents signal amplification, consistent with the high PBSI score. This result substantiates that a system predicted to be parallel based on systemic feature visualization indeed yielded a GRN structure with high PBSI.

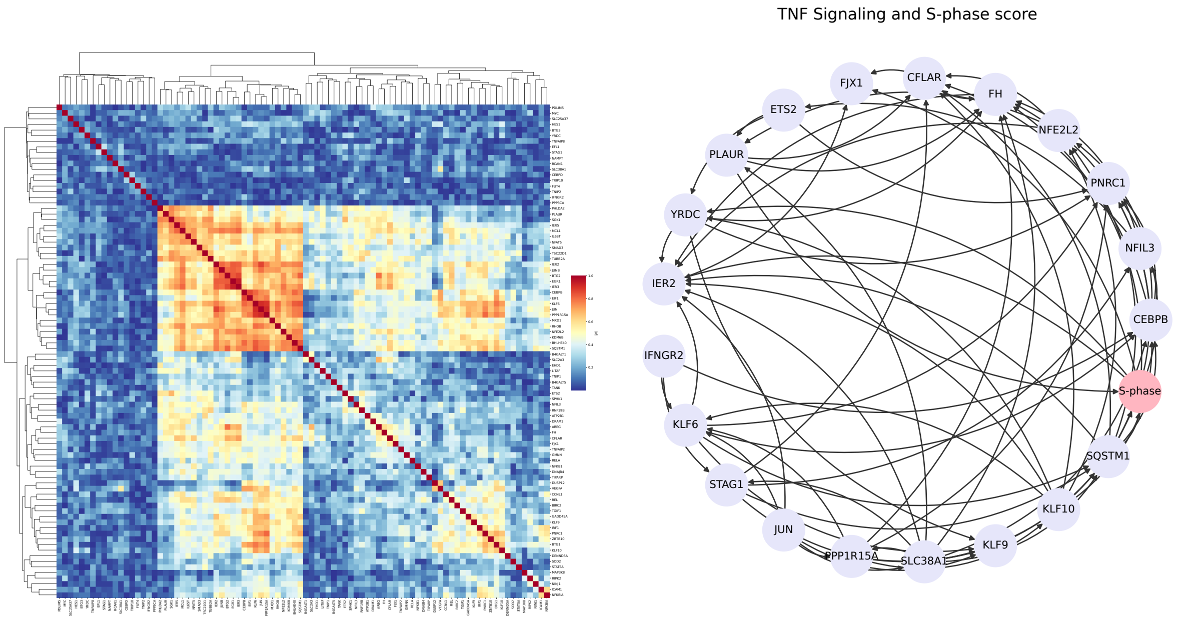


(a) Systemic features of TNF signaling

(b) Inferred DAG (TNF signaling)

**Figure S23: TNF signaling as a representative example of a parallel GRN topology**

**a**: Heatmap displaying the absolute Pearson correlation coefficients among all genes associated with TNF-$\alpha$ signaling via NF-$\kappa$B (MSigDB Hallmark 2020). **b**: The inferred DAG representing the effect of TNF signaling on the S-phase score. Note that edges here represent structural dependencies only; unlike the final GRN, signs (positive/negative edges) have not yet been assigned.

As an additional concrete example, we investigated the relationship between apoptosis-associated genes and the S-phase score. It is well-established that apoptosis regulation and cell cycle progression are intimately coupled via multiple mechanisms [PKG00, CL11], implying the presence of parallel gene modules. Consistent with this expectation, the heatmap of absolute Pearson correlations for all genes in the `Apoptosis` category exhibited a distinct block-diagonal structure (Figure S24a).

Using the Apoptosis score calculated from these genes as a reference (Figure S24b), we selected the top 20 genes with low mutual correlations (Figure S24c). Subsequent inference of the DAG and GRN using these selected genes resulted in a topology characterized by fragmented cascades and a high PBSI score (PBSI $> 0.6$), notably featuring negative terminal edges even within complex interaction chains (Figures S24d–S24f).

Consequently, these results consistently underscore that in multiple biological contexts where parallel gene modules are anticipated, the inferred GRNs exhibit high PBSI scores, indicating suitability for the PB design. Furthermore, this empirical evidence aligns with the theoretical insights derived from the comparison of Models $\Delta$ and $\Sigma$: specifically, it confirms that PB suitability remains robust even in the presence of intra-module interaction complexity, provided that the terminal regulatory edges are negative. Crucially, this finding implies more than just a

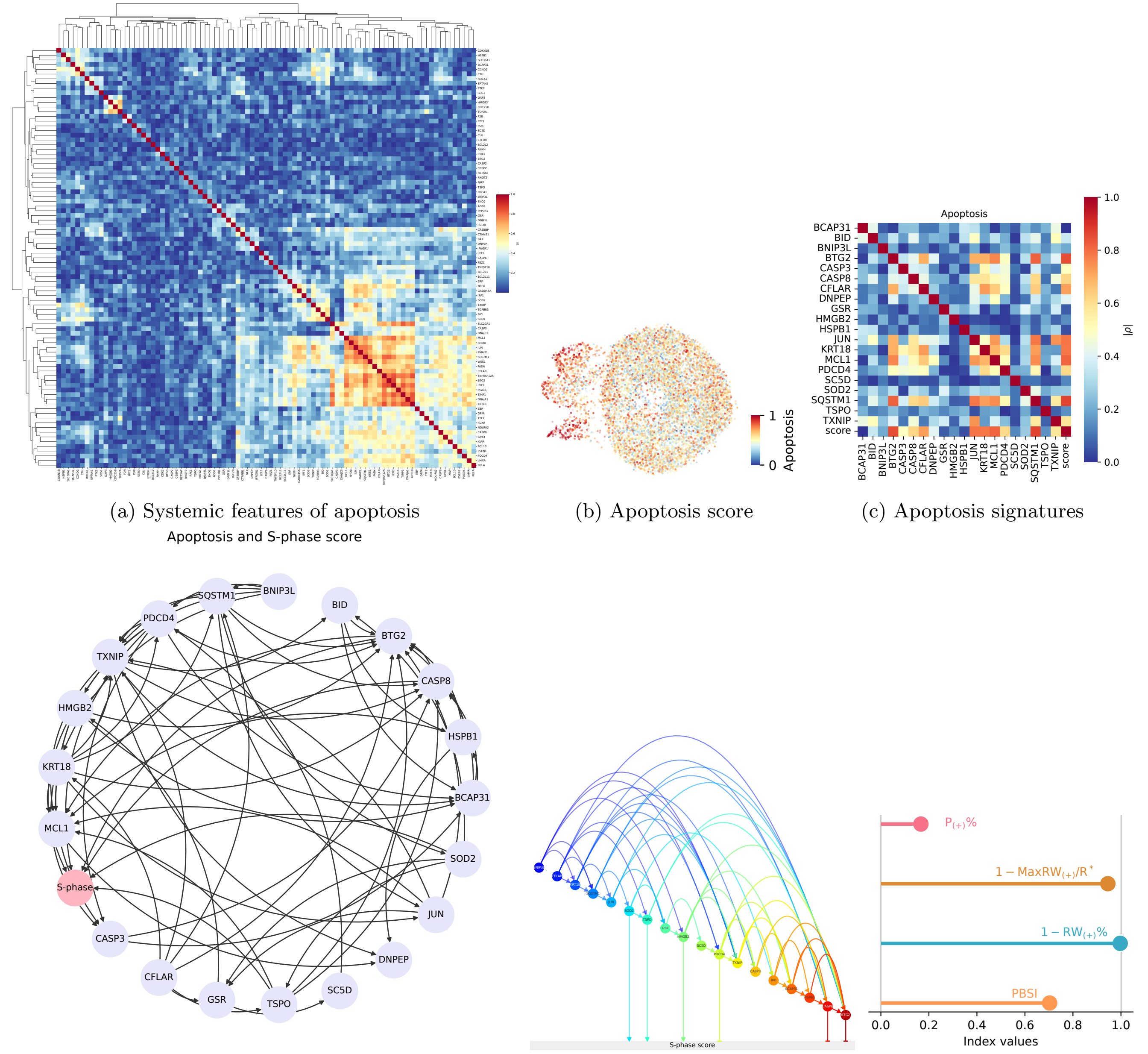


(a) Systemic features of apoptosis (b) Apoptosis score (c) Apoptosis signatures

(d) Inferred DAG (apoptosis) (e) GRN (apoptosis) (f) Indices (apoptosis)

**Figure S24: Apoptosis as another example of a parallel GRN topology**

**a**: Heatmap displaying the absolute Pearson correlation coefficients among all genes associated with apoptosis (MSigDB Hallmark 2020). **b**: UMAP plot colored by the apoptosis score. **c**: The absolute Pearson correlation coefficients among the 20 low-correlated genes and the phenotypic score. **d–f**: The inferred DAG (d), the inferred GRN representing the effect of apoptosis on the S-phase score (e), and its four topology indices (f). The high PBSI score ($> 0.6$) indicates that the GRN, consisting of low-correlated apoptosis-related genes, exhibits fragmented cascades, as reflected by the high values of $1 - \text{MaxRW}_{(+)}/\text{R}^*$ and $1 - \text{RW}_{(+)}\%$. Although the topology includes some entangled substructures, the presence of negative terminal edges effectively prevents signal amplification. Note that edges in (d) represent structural dependencies only; unlike the final GRN, signs (positive/negative edges) have not yet been assigned.

methodological demonstration of constructing additive systems; it supports the view that the insights derived from our minimalist simulators are reproducible in real-world scRNA-seq data possessing inherent biological complexity.

##### I.4.2 Examples of C+LOO-suited GRN topologies defying orthogonal decomposability

In this section, we examine Myc response and G2/M-phase as concrete examples of entangled GRN topologies enabling signal amplification. Crucially, as demonstrated in our theoretical analysis, the mere presence of an entangled structure within the target gene module is insufficient to confirm signal amplification; the complex interactions must ultimately exert a positive effect on the evaluated phenotype. Therefore, we selected the Myc response and G2/M-phase as specific cases where prior knowledge suggests both a highly interconnected (entangled) structure and a positive correlation with the S-phase score, thereby fulfilling the prerequisites for signal amplification.

We first consider the impact of Myc response on the S-phase score. Calculating the absolute Pearson correlations for genes listed in the `Myc Targets V1` category reveals a large cluster of highly correlated genes ("red square") accompanied by moderate off-diagonal correlations ("yellow shadow"), creating a macroscopically diffuse region of red-to-yellow gradients (Figure S25a). This pattern suggests the potential for modeling the system as an entangled mechanistic network.

In comprehensive lists such as the MSigDB Hallmark 2020 collection, not all genes are necessarily highly correlated. However, by strategically selecting target genes from such diffuse, highly interconnected areas, we can increase the complexity of interactions within the target set. While Myc is strongly associated with cell cycle progression—evidenced by the high correlation between the Myc V1 score and the proliferation marker MKI67 in metastatic breast cancer [SOET20, BDL15]—predicting the resulting topology is non-trivial. An increase in direct positive paths ($P_{(+)}\%$) from targets to the S-phase score would ostensibly favor a parallel topology. However, the balance between these direct effects and the extensive inter-gene crosstalk (entanglement) cannot be determined solely from qualitative biological knowledge. Therefore, we focused our verification on determining whether chains of positive regulations emerge when target genes are selected to maximize gene–gene interactions.

To explicitly construct an entangled GRN, we selected the top 20 genes with the highest mutual correlations (see Methods 4.9.3). The correlation structure of these selected genes is shown in Figure 6e. We then inferred a DAG for these 20 genes and the S-phase score (Figure S25b). The resulting GRN topology exhibited a dense, interconnected structure and a low PBSI score ($< 0.6$) (Figures 6h–6i). Notably, the terminal edges emerging from these complex interactions are positive, thereby enabling the signal amplification mechanism characterized in Model $\Delta$. This result demonstrates that a target gene–readout pair anticipated to exhibit signal amplification indeed yielded a GRN structure with low PBSI, aligning with the theoretical preference for the amplification-adapted C+LOO design over the orthogonality-enforcing PB design.

Next, we analyzed the association between the G2/M-phase checkpoint genes and the S-phase score. We selected this category because the biological overlap between `G2/M Checkpoint` genes and `E2F Targets` (which constitute the S-phase score) allows us to anticipate a positive correlation with the phenotype for at least a subset of genes. Similar to the Myc response, determining whether the underlying biological context is purely entangled solely from a large-scale gene function thesaurus is challenging. Indeed, the heatmap of gene–gene correlations revealed a pattern characterized by a large cluster of high correlations ("red square") surrounded by diffuse moderate correlations ("yellow shadow") (Figure S26a), presenting a structural ambiguity similar to that observed in the Myc analysis.

To clarify this ambiguity and explicitly test the impact of potential entanglement, we selected the top 20 genes with the highest correlations to the G2/M-phase score (Figures S26b–S26c), intending to construct a maximally entangled topology. The resulting DAG and GRN exhibited a densely interconnected structure with a low PBSI score ($< 0.6$), confirming the formation of an entangled topology (Figures S26d–S26f). Crucially, consistent with the positive correlation anticipated from the biological overlap, the complex interactions within this GRN formed chains of positive edges leading to the S-phase score, notably culminating in positive terminal edges. This structure enables the signal amplification effect, thereby supporting the suitability of the amplification-adapted C+LOO design in this context as well.

Collectively, the comprehensive analysis of these four biological contexts—spanning from TNF signaling and Apoptosis to Myc response and G2/M phase—provides empirical support for the theoretical framework established by our simulators. The workflow presented here successfully constructed biologically plausible GRN topologies, where the calculated PBSI scores aligned consistently with their structural interpretations, demonstrating that researchers can infer optimal experimental designs through a quantitative scheme derived solely from plain scRNA-seq data. This suggests that PBSI accurately captures the subtle distinction between "complex but additive" systems (e.g., Apoptosis) and "amplifying and nonlinear" systems (e.g., Myc). Consequently, these findings extend beyond merely assisting practical design selection for Perturb-seq studies; they support the core theoretical insight

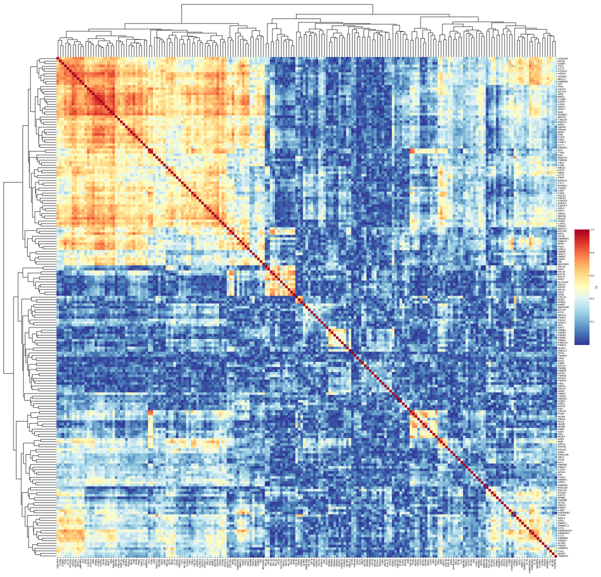

(a) Systemic features of Myc response

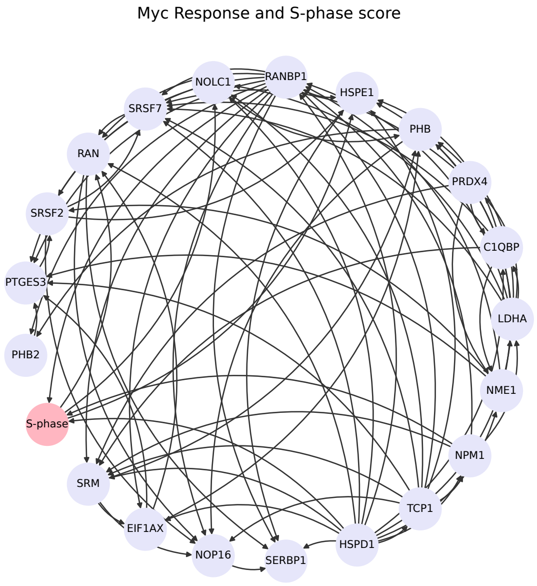


(b) Inferred DAG (Myc response)

**Figure S25: Myc response as a representative example of a parallel GRN topology**

**a**: Heatmap displaying the absolute Pearson correlation coefficients among all genes associated with Myc targets (MSigDB Hallmark 2020). **b**: The inferred DAG representing the effect of Myc response on the S-phase score. Note that edges here represent structural dependencies only; unlike the final GRN, signs (positive/negative edges) have not yet been assigned.

derived from our minimalist simulators—specifically, that the suboptimality of orthogonal PB designs due to signal amplification-induced nonlinearity is not a hypothetical artifact, but a tangible reality in biological systems.

##### I.4.3 Supplementary Note: Validity of using statistical independence and correlation as proxies for causality

While statistical independence (or zero correlation) does not theoretically preclude causality—as interactions can manifest as non-monotonic functions (e.g., U-shaped responses) with negligible linear correlation—inferring structural edges from statistical dependence is a standard methodological convention, foundational to constraint-based causal discovery frameworks [SGS00, OKO23]. Moreover, biological regulatory networks are inherently sparse [BO04, GF05], constrained by biochemical specificity (e.g., transcription factor binding preferences [LJC$^+$18]) and physical chromatin organization (e.g., topologically associating domains [LKH$^+$15]). Crucially, these constraints extend beyond "physical models" of direct binding to shape the "influence models" inferred from transcriptomic data, as the latter implicitly reflect the underlying physical architecture [GF05].

Consequently, observing a lack of correlation is far more likely to stem from the topological absence of regulatory edges rather than the perfect cancellation of complex nonlinear interactions. Furthermore, apparent non-monotonic responses often emerge as network-level properties derived from combinations of monotonic interactions (analogous to neural networks approximating nonlinear functions via ReLU activation). Even if direct non-monotonic edges exist in specific cases, they do not negate our broader argument; we employ correlation here as a practical heuristic to demonstrate that "PB-suited" GRNs are statistically plausible outcomes of candidate selection in real-world Perturb-seq studies. Thus, treating uncorrelated factors as functionally independent serves as the most structurally parsimonious and rational default.

Conversely, the presence of correlation does not guarantee direct causality, often arising from spurious associations. In the context of causal discovery, "spurious correlations" caused by conditioning on observed variables (e.g., colliders) can generally be resolved by examining conditional independence to refine the DAG structure. However, correlations arising from unobserved confounders or variables excluded from the model pose a more significant challenge. To mitigate this, we employed a strategy that combines rigorous variable selection grounded in domain knowledge (outsourced here to the curated MSigDB gene sets) with careful scrutiny of the data structure via

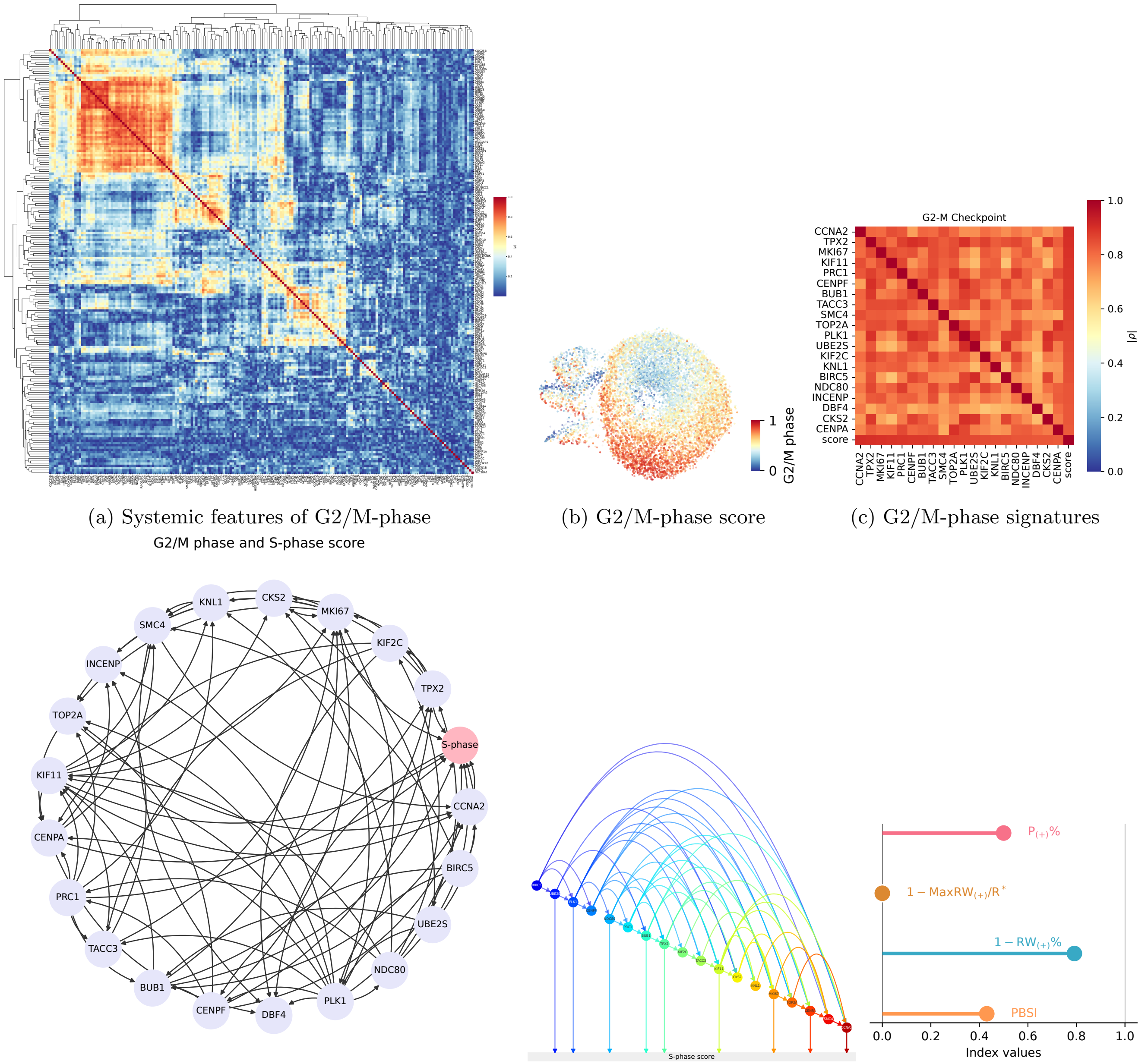


(a) Systemic features of G2/M-phase

(b) G2/M-phase score

(c) G2/M-phase signatures

(d) Inferred DAG (G2/M-phase)

(e) GRN (G2/M-phase)

(f) Indices (G2/M-phase)

**Figure S26: G2/M-phase as another example of an entangled GRN topology**

**a**: Heatmap displaying the absolute Pearson correlation coefficients among all genes associated with G2/M checkpoint (MSigDB Hallmark 2020). **b**: UMAP plot colored by the G2/M-phase score. **c**: The absolute Pearson correlation coefficients among the top 20 high-correlated genes and the phenotypic score. **d**–**f**: The inferred DAG (d), the inferred GRN representing the effect of G2/M-phase on the S-phase score (e), and its four topology indices (f). The low PBSI score ($< 0.6$) indicates that the GRN, consisting of high-correlated G2/M-phase-related genes, exhibits chains of positive gene–gene interactions, as reflected by the low values of $1 - \mathrm{MaxRW}_{(+)}/\mathrm{R}^*$ and $1 - \mathrm{RW}_{(+)}\%$. Note that edges in (d) represent structural dependencies only; unlike the final GRN, signs (positive/negative edges) have not yet been assigned.

visualization, rather than relying blindly on automated inference.

Regarding the influence of exogenous variables, it is essential to recognize the intrinsic limitations of observational data. Since scRNA-seq data (count matrices) are mathematically treated as a vector space where measured samples and genes act as dual bases, the gene functions represented within the data structure are inherently defined and constrained by the observed sample space. Without introducing external structural priors as domain knowledge, extrapolating gene functions beyond the measurement scope or visualizing the effects of unobserved exogenous variables is theoretically ill-posed. Unlike interventional systems such as Perturb-seq—where causal effects on targeted variables are verifiable and non-targeted variables can be partly controlled (e.g., via dummy variables)—observational plain scRNA-seq data, particularly from a single batch, lack such interventional anchors, making the technical handling of unobserved confounders difficult.

In light of these limitations, we emphasize that the networks constructed in this study are strictly defined as "inferred GRNs" and do not necessarily correspond to the ground-truth biological networks. Specifically, the structure learning algorithm employed here—Hill Climb Search based on the Bayesian Information Criterion score—was selected for its flexibility in DAG design but is a greedy algorithm known for potential issues with reproducibility and local optima. Therefore, the inferred topologies should be interpreted not as definitive biological discoveries, but as provisional inference models sufficient for assessing the statistical properties of PBSI.

### J Practical guidelines for experimental design in multivariate biology

In this section, we synthesize the findings of this study into practical guidelines for planning experimental designs in multivariate biological experiments. While this article has primarily focused on Perturb-seq as a case study, the C+LOO design has a long tradition in biological experimentation—exemplified by the discovery of the four pluripotency-inducing factors [TY06]. Since the mathematical properties of the C+LOO design discussed herein are inherent to the design matrix itself, the following discussions are applicable not only to Perturb-seq but also to other experimental modalities employing similar screening strategies.

#### J.1 Selection of candidate genes

##### Plausibility of candidates matters to choosing an optimal experimental design

As demonstrated in Results 2.2 and SI G.3, when the selection of candidate genes is poor (i.e., the number of factors $n$ is significantly larger than the number of active factors $n^*$; $n \gg n^*$), the optimal experimental design fluctuates depending on the noise level. As noted in SI D.2, while we could benchmark C+LOO and PB designs under specific noise levels using ground truth labels in our simulations, such *a priori* knowledge is rarely available in practice. Therefore, it is practically crucial to ensure that the candidate genes selected for the experiment are as *plausible* as possible to avoid the sparse factor scenario.

##### A large candidate gene list is preferable in C+LOO

When adopting the C+LOO design, maintaining a sufficiently large size for the candidate gene list is critical from a statistical perspective. As shown in Results 2.1 and Figure 1c, there exists an $n$-dependent trade-off among the five correlation functions (Equations (10)–(14); derived in SI B). Specifically, correlations involving a shared factor ($\rho_{i\times j,i}$ and $\rho_{i\times j,i\times l}$) increase in magnitude with $n$, whereas correlations between distinct factors ($\rho_{i,j}$, $\rho_{i\times j,k}$, and $\rho_{i\times j,k\times l}$) tend toward zero. Although the severity and patterns of these inherent correlations vary depending on the number of factors, certain types of correlations are unavoidable, underscoring the strong association between the C+LOO design and factor correlations.

Among the five correlation types, $\rho_{i,j}$ and $\rho_{i\times j,i}$ are particularly unique to the C+LOO design. In the field of DOE, main effects are generally regarded as fundamental variables [Lan03]; thus, orthogonal designs such as the FF and PB ensure $\rho_{i,j} = 0$ to enable a clear distinction between main effects (SI C). These designs also typically yield $\rho_{i\times j,i} = 0$, allowing the effect of $\mathbf{x}_i$ to be evaluated independently of $\mathbf{x}_j$ and their interaction $\mathbf{x}_{i\times j}$. In contrast, C+LOO designs produce non-zero values for both $\rho_{i,j}$ and $\rho_{i\times j,i}$, causing the estimated main effects to be mutually confounded. The presence of $\rho_{i,j}$ is especially undesirable because confounding $\mathbf{x}_i$ with $\mathbf{x}_j$ leads to serious interpretability issues.

To mitigate this, a large $n$ is preferable in the C+LOO design. According to Equation (10), $n \geq 4$ is required to achieve $|\rho_{i,j}| < 0.3$, and $n \geq 11$ is necessary to suppress it to $|\rho_{i,j}| < 0.1$.

In contrast, C+LOO designs with large $n$ produce non-zero $\rho_{i\times j,i}$, implying that the main effect $\mathbf{x}_i$ is assessed under the implicit influence of all other factors $\mathbf{x}_j$ via their interactions $\mathbf{x}_{i\times j}$. Although $\rho_{i\times j,i}$ (and $\rho_{i\times j,i\times l}$) can be suppressed with small $n$, these effects persist to a meaningful degree even at large $n$—for example, $\rho_{i\times j,i}$ ranges from 0.5 to $\sqrt{1/2} \approx 0.707$, according to Equation (16).

While this dilemma is unavoidable due to the $n$-dependent trade-off, this residual factor correlation, particularly $\rho_{i\times j,i}$, constitutes the "leaky" nature of the C+LOO design. As discussed below, this property is a unique characteristic not found in orthogonal DOE designs and can function positively in certain biological contexts. Therefore, the persistence of these correlations at large $n$ should not be viewed entirely negatively; rather, the design choice should be based on whether the specific biological system benefits from this leakage.

In this regard, maximizing $n$ to suppress the more problematic $\rho_{i,j}$ is generally the recommended strategy for C+LOO designs.

**Preliminary validation is also practically important for dealing with budgetary constraints**

However, it is crucial to consider budgetary constraints. In scRNA-seq, there is often a trade-off between sample size and sequencing depth under limited budgets [LMPH18]. Similarly, since replication is critical in Perturb-seq, a trade-off exists between the number of target genes and replication levels. For reference, it has been reported that $N = 8$–$12$ replicates per condition are required for robust differential expression analysis in mouse bulk RNA-seq [HSN+25]; securing such replication numbers in high-throughput screens inevitably strains the budget. While ample funding would allow for the examination of numerous factors with maximized replication and sequencing depth, the extent to which these parameters can be compromised depends heavily on the biological phenomena and sample characteristics. Therefore, we strongly recommend conducting preliminary experiments or simulations to validate detectability.

Using regular scRNA-seq datasets without perturbation—which can be more accessible in terms of experimental cost or are already available in databases such as the Human Cell Atlas [RTL+17]—researchers can prioritize plausible genes. Furthermore, by employing scRNA-seq simulators such as Splatter [ZPO17] to virtually adjust sequencing depth, one can verify whether candidate genes can be appropriately detected (e.g., whether they appear in appropriate positions on a volcano plot as differentially expressed genes). This approach facilitates finding an optimal balance between sequencing depth and replication number before performing the actual Perturb-seq experiment.

#### J.2 Experimental design choice: C+LOO vs. PB

As summarized in SI I.3.4, a strategic approach involves adopting the PB design as the default baseline, reserving the C+LOO design only for cases where strong signal amplification is suspected. This recommendation reflects the broad applicability of the PB design, supported by our exhaustive analysis of ESM4 and random sampling of ESM9 (Figure 5b and Figure S15b), which demonstrated that the majority of network topologies are classified as "PB-suited."

Conversely, in systems characterized by strong signal amplification, the "leaky" nature of the C+LOO design enables a comprehensive evaluation that implicitly includes interaction effects. While main effects are, by definition, calculated by averaging out the influence of other factors to isolate the specific impact of the factor of interest [CSC20], in systems where gene–gene interactions lead to significant signal amplification, the extreme magnitude of these amplified values persists after averaging, thereby substantially inflating the estimated main effects. This phenomenon represents a violation of the homoscedasticity assumption underlying the standard calculation of main effects. In such scenarios, the C+LOO design, despite possessing unfavorable properties from the standpoint of classical DOE, becomes relatively adaptive; its inherent correlation structure effectively propagates the impact of interaction terms into the main effect estimates, thereby yielding results that paradoxically align closer with the intended inference than those from a strictly orthogonal design.

However, as discussed in SI I.3.2, the manifestation of signal amplification requires that all constituent edges within an interaction chain be positive. Qualitatively identifying such conditions in general biological systems requires precise and extensive domain knowledge. Therefore, rather than relying solely on qualitative assumptions, we recommend the workflow demonstrated in SI I.4: utilizing unperturbed scRNA-seq data to perform GRN inference and making quantitative decisions based on PBSI. As established in SI I.3.1, a value of PBSI $> 0.6$ serves as the practical threshold for recommending the PB design.

Regarding the methodology for PBSI calculation, while this study implemented a Bayesian network-based GRN inference to prioritize the estimation of edges for variables not explicitly present in scRNA-seq data (e.g., phenotypic scores), the framework is methodology-agnostic. Any algorithm capable of inferring GRNs (including output values) from standard unperturbed scRNA-seq datasets—such as SCENIC [AGBM+17] or decoupleR [BiMVSB+22]—can be applied to calculate PBSI.

From a wet-lab perspective, implementing PB designs with CRISPR-based perturbations involves simultaneous KDs of multiple genes. This introduces technical challenges, such as controlling off-target effects and optimizing viral titers to ensure efficient multiplicity of infection. Consequently, the practical number of factors $n$ manageable in a PB design is likely limited to approximately $n < 10$. For screening larger gene sets, the C+LOO design—which does not require simultaneous perturbations—might be considered. However, researchers must be cautious, as C+LOO performance degrades significantly in noise-prone systems, particularly those with parallel GRN topologies (Figures S8b and S18d). In many cases, it may be strategically wiser to reduce the number of target genes to fit within the constraints of a PB design, thereby reallocating resources to increase replication numbers or sequencing depth to enhance measurement quality.

Thus, when planning large-scale experiments, rigorous preliminary experiments are essential, including the option of narrowing down targets to a critical subset to maintain a manageable $n$. The approach demonstrated in SI I.4, which leverages plain scRNA-seq data to discuss gene selection in the context of data structure, provides a rational framework for such *a priori* optimization.

#### J.3 Strategies in the absence of preliminary scRNA-seq data

Thus far, we have emphasized the importance of preliminary experiments using scRNA-seq for candidate gene selection and experimental design decisions. However, we must also consider scenarios where preliminary plain scRNA-seq is unavailable. While well-funded projects may afford preliminary sequencing, budget-constrained projects or experiments using non-sequencing, low-throughput readouts may lack this data.

In such situations, it is necessary to perform candidate gene selection and GRN inference in a data-free manner. We recommend leveraging domain knowledge or public databases such as KEGG [KG00] to construct hypothetical GRNs and calculate PBSI. However, interactions documented in global databases are not guaranteed to be active in the specific cellular state of interest. Although the global maps of canonical KEGG pathways aggregate distinct cellular contexts [KG00], the topology of the GRN relevant to an experiment depends entirely on the selection of candidate genes. Indeed, the *distributional hypothesis of gene function* posits that gene functions are defined by their context-specific collocations [KPG+24]. This implies that interactions documented in global reference maps can be highly context-dependent rather than universally preserved; in any specific cellular state, many "known" edges functionally vanish. Therefore, it is crucial to construct the GRN blueprint under the critical supervision of the experimenter's domain expertise, selectively incorporating only biologically plausible edges.

The ability to infer appropriate experimental designs from such qualitative information represents a distinct advantage of PBSI. Unlike generative model-based simulations, which necessitate plain scRNA-seq data as a prerequisite for design comparison, PBSI allows for the estimation of suitable designs even from a rough sketch of the GRN derived from fragmented information. In this regard, PBSI is expected to serve as a smooth bridge between knowledge-driven and data-driven research. By guiding appropriate design selection to avoid the pitfall of naively adopting the C+LOO design, PBSI facilitates robust Perturb-seq planning even when relying on qualitative inference.

When the full picture of the GRN or edge directions is undetermined (note that PBSI targets DAGs), we recommend the following heuristic:

- If signaling amplification is strongly suspected, adopt the C+LOO design to leverage interaction leakage.
- In all other cases, the PB design is the more robust and safer choice.

Empirical evidence from Models $\Phi$, $\Psi$, $\Lambda$, $\Pi$, $\Delta$, and $\Sigma$ confirms that the PB design maintains relatively stable statistical power compared to C+LOO, even under difficult conditions such as high noise and low replication numbers (Figures S8c–S10c and S18e–S20e). Furthermore, in our exhaustive analysis of ESM4 and random sampling of ESM9, the majority of networks were classified as "PB-suited" (Figure 5b and Figure S15b). Therefore, unless there is a compelling reason to suspect amplification-driven dominance, treating orthogonal designs (such as PB) as the default is a scientifically rational strategy.

Regarding experimental costs, although the C+LOO design can reduce the number of batches compared to PB, the difference is at most $3N$ relative to the replication number $N$ (Figure 1d). For a large number of factors $n$, this relative impact is minor. While the impact is relatively larger when $n$ is small, as mentioned earlier, a large $n$ is recommended for C+LOO to avoid undesirable factor correlations (such as $\rho_{i,j}$). Thus, even in these situations, the guideline to prioritize the PB design remains valid.

#### J.4 Analytical framework: Shifting from pairwise comparisons to MLR/ANOVA

As mentioned in SI G.4, our evaluation of Models $\Phi$, $\Psi$, and $\Lambda$ compared two analytical approaches for C+LOO-based data: conventional pairwise comparisons using Dunnett's test versus multivariate analysis (MLR and ANOVA) for estimating main effects. The results suggested that multivariate analysis yields equivalent or slightly superior performance in terms of Cohen's $\kappa$ (Figures S8b–S10b). While this similarity can be attributed to the "leaky" nature of the C+LOO design which implicitly incorporates interaction effects, it highlights a critical opportunity: performance can be enhanced strictly *post-hoc* by merely updating the analytical framework, incurring no additional financial cost.

Furthermore, a noteworthy finding is that in terms of statistical power, multivariate analysis outperformed Dunnett's test, particularly under high-noise conditions (Figures S8c–S10c). This indicates that a simple transition from Dunnett's test to MLR/ANOVA significantly contributes to the stability of data interpretation.

Multivariate analysis is, of course, not new to biology. Especially in the modern era, where high-throughput quantitative data is easily accessible, MLR has established itself as a fundamental analytical tool in experimental biology as well. Thus, the application of multivariate models is not a novel concept imported solely through the recent integration of biology and DOE. Rather, the prevalence of pairwise comparisons likely stems from a conceptual coupling: the C+LOO design is historically rooted in the philosophy of assessing differences against a fixed "baseline," leading to a default association where employing the C+LOO design implies conducting pairwise testing.

Conversely, while ANOVA is widely used in biology, its application is often limited to simple one- or two-way designs. It is unclear whether the know-how to perform $n$-way ANOVA on highly unbalanced datasets like C+LOO has sufficiently permeated the experimental biology community. Although statistically standard, we have explicitly detailed the procedure for conducting Type II ANOVA and subsequent power analysis within the general linear model framework in Methods 4.4.4. This formulation empowers researchers to implement rigorous statistical testing and power calculations for *any* arbitrary design matrix, including D-optimized designs. To lower the implementation barrier, we provide the corresponding code in our GitHub repository (https://github.com/yo-aka-gene/WhyDOE). Since the core logic relies on simple matrix operations—aside from specific functions in `statsmodels` [SP10] or `SciPy` [VGO+20]—the framework is highly accessible and adaptable.

Looking forward, the importance of preliminary experiments and simulations will only increase. Large-scale generative models for Perturb-seq consistently report higher accuracy in predicting unseen single-gene perturbations compared to combinatorial ones [VTWP+25]. Consequently, generating virtual data based on the C+LOO design—which by definition excludes simultaneous KDs—represents the most feasible and reliable near-term application of these models. In such scenarios, researchers could generate virtual C+LOO-based Perturb-seq data from plain scRNA-seq inputs to perform power analysis, enabling rigorous sample size planning. As discussed in SI J.1, given that replication number trades off against sequencing depth under strict budgetary constraints, the ability to perform precise sample size design is of paramount importance.

We anticipate that such DOE-oriented methodologies will become increasingly generalized, transforming not only experimental design but also the fundamental frameworks of data analysis in multivariate biology.

#### J.5 Result interpretation: on translating readouts from statistical analysis into biological language

Finally, we must address the fundamental limitations regarding the interpretation of results derived from Perturb-seq and DOE. As demonstrated in Figure S7, *Interaction Leakage* (Equation (S-60)) can be non-zero in both C+LOO and PB designs (except for specific minimal cases like $n = 2$ in PB). Even when employing high-resolution designs like the FF design or calculating main effects strictly by definition, interactions can implicitly influence main effect estimates in biological settings. This is particularly true in systems characterized by signal amplification, where the scale of responses differs vastly between positive and negative conditions. In such domains, the mathematical operation of "centering"—assuming symmetric cancellation of effects around a zero mean—lacks biological plausibility due to the non-negative nature of gene expression.

Consequently, there is no guarantee that mechanistic explanations derived from these experiments will be reproducible across different experimental contexts (e.g., different species, cellular states, cell culture protocols, or perturbation conditions). Generalizing Perturb-seq results as immutable "intrinsic properties" of genes is therefore fraught with risk. Fundamentally, in high-dimensional transcriptomic data, gene functions and sample characteristics constitute a dual basis system. A gene's functional identity is mathematically defined only within the vector space spanned by the observed samples (i.e., the cellular states under specific perturbations). This implies that *extrapolability* is never axiomatic; the function observed in one context does not necessarily hold in another.

This limitation is inevitable even with MLR/ANOVA, which mitigates context-dependency arising from baseline-dependency in pairwise comparisons (introduced in SI E).

Therefore, we urge researchers to exercise caution against allowing the interpretation of experimental results to detach from their premises. Perturb-seq results should be viewed not as universal biological truths, but as context-dependent descriptions of system behavior under specific constraints. This perspective resonates with the implications of the *distributional hypothesis of gene function* [KPG+24] and reinforces our recommendation that constructing hypothetical GRNs from global databases requires critical expert supervision to ensure context-specific validity (SI J.3).

#### J.6 Perspectives for detecting combinatorial effects

In this paper, we did not discuss the framework of verifying combinatorial effects by creating double-KD conditions for a specific subset of candidate genes in addition to the C+LOO design, a practice seen in recent studies. These methods excel in flexibility, allowing for *post-hoc* verification of combinatorial effects based on the C+LOO design. However, since the C+LOO design itself exhibits $n$-dependent changes in factor correlation structure, and adding specific gene pairs *post-hoc* further alters this structure in an *ad-hoc* manner, it is difficult to discuss the properties of such approaches from a comprehensive, theoretical standpoint.

On the other hand, DOE provides established frameworks for analyzing such combinatorial effects. While the PB design leaves the majority of two-factor interaction terms correlated with each other (Figure S2c), designs with higher resolution, such as definitive screening designs [JN11], can resolve this issue while allowing for the verification of two-factor interaction terms. Furthermore, these methods allow for combinatorial optimization using response surface methodology, holding significant potential for application in biological experiments [MSS24].

Although a comparative discussion of these methods is beyond the scope of this paper, we anticipate further developments in this area.

[KPG+24] Jason J Kwon, Joshua Pan, Guadalupe Gonzalez, William C Hahn, and Marinka Zitnik. On knowing a gene: A distributional hypothesis of gene function. *Cell Systems*, 2024.

[Lan03] Øyvind Langsrud. ANOVA for unbalanced data: Use Type II instead of Type III sums of squares. *Statistics and computing*, 13(2):163–167, 2003.

[LBC+22] Malte D Luecken, Maren Büttner, Kridsadakorn Chaichoompu, Anna Danese, Marta Interlandi, Michaela F Müller, Daniel C Strobl, Luke Zappia, Martin Dugas, Maria Colomé-Tatché, et al. Benchmarking atlas-level data integration in single-cell genomics. *Nature methods*, 19(1):41–50, 2022.

[LBT+15] Arthur Liberzon, Chet Birger, Helga Thorvaldsdóttir, Mahmoud Ghandi, Jill P Mesirov, and Pablo Tamayo. The molecular signatures database hallmark gene set collection. *Cell systems*, 1(6):417–425, 2015.

[LJC+18] Samuel A Lambert, Arttu Jolma, Laura F Campitelli, Pratyush K Das, Yimeng Yin, Mihai Albu, Xiaoting Chen, Jussi Taipale, Timothy R Hughes, and Matthew T Weirauch. The human transcription factors. *Cell*, 172(4):650–665, 2018.

[LKH+15] Darío G Lupiáñez, Katerina Kraft, Verena Heinrich, Peter Krawitz, Francesco Brancati, Eva Klopocki, Denise Horn, Hülya Kayserili, John M Opitz, Renata Laxova, et al. Disruptions of topological chromatin domains cause pathogenic rewiring of gene-enhancer interactions. *Cell*, 161(5):1012–1025, 2015.

[LMPH18] Atefeh Lafzi, Catia Moutinho, Simone Picelli, and Holger Heyn. Tutorial: guidelines for the experimental design of single-cell RNA sequencing studies. *Nature protocols*, 13(12):2742–2757, 2018.

[LR21] W Brent Lindquist and Svetlozar T Rachev. Taylor's law and heavy-tailed distributions. *Proceedings of the National Academy of Sciences*, 118(50):e2118893118, 2021.

[MSS24] Seonyun Moon, Anna Saboe, and Michael J Smanski. Using design of experiments to guide genetic optimization of engineered metabolic pathways. *Journal of Industrial Microbiology and Biotechnology*, 51:kuae010, 2024.

[NIGM18] Chigozie Nwankpa, Winifred Ijomah, Anthony Gachagan, and Stephen Marshall. Activation functions: Comparison of trends in practice and research for deep learning. *arXiv preprint arXiv:1811.03378*, 2018.

[OKO23] Yuji Okano, Yoshitaka Kase, and Hideyuki Okano. A set-theoretic definition of cell types with an algebraic structure on gene regulatory networks and application in annotation of RNA-seq data. *Stem cell reports*, 18(1):113–130, 2023.

[PKG00] Bruna Pucci, Margaret Kasten, and Antonio Giordano. Cell cycle and apoptosis. *Neoplasia*, 2(4):291–299, 2000.

[PMC+21] Lucrezia Patruno, Davide Maspero, Francesco Craighero, Fabrizio Angaroni, Marco Antoniotti, and Alex Graudenzi. A review of computational strategies for denoising and imputation of single-cell transcriptomic data. *Briefings in bioinformatics*, 22(4):bbaa222, 2021.

[RHL24] Yusuf Roohani, Kexin Huang, and Jure Leskovec. Predicting transcriptional outcomes of novel multigene perturbations with GEARS. *Nature Biotechnology*, 42(6):927–935, 2024.

[RSP+22] Joseph M Replogle, Reuben A Saunders, Angela N Pogson, Jeffrey A Hussmann, Alexander Lenail, Alina Guna, Lauren Mascibroda, Eric J Wagner, Karen Adelman, Gila Lithwick-Yanai, et al. Mapping information-rich genotype-phenotype landscapes with genome-scale Perturb-seq. *Cell*, 185(14):2559–2575, 2022.

[RTL+17] Aviv Regev, Sarah A Teichmann, Eric S Lander, Ido Amit, Christophe Benoist, Ewan Birney, Bernd Bodenmiller, Peter Campbell, Piero Carninci, Menna Clatworthy, et al. The human cell atlas. *elife*, 6:e27041, 2017.

[SGS00] Peter Spirtes, Clark N Glymour, and Richard Scheines. *Causation, prediction, and search.* MIT press, 2000.

[SL03] George AF Seber and Alan J Lee. *Linear regression analysis.* John Wiley & Sons, 2003.

[SOET20] Amy Schulze, Masanori Oshi, Itaru Endo, and Kazuaki Takabe. Myc targets scores are associated with cancer aggressiveness and poor survival in er-positive primary and metastatic breast cancer. *International journal of molecular sciences*, 21(21):8127, 2020.

[SP10] Skipper Seabold and Josef Perktold. Statsmodels: econometric and statistical modeling with python. *SciPy*, 7(1), 2010.

[Tay61] LR Taylor. Aggregation, mean and variance. *Nature*, 189:732–735, 1961.

[TBR+15] Jun Tsuyama, Jens Bunt, Linda J Richards, Hiroko Iwanari, Yasuhiro Mochizuki, Takao Hamakubo, Takuya Shimazaki, and Hideyuki Okano. MicroRNA-153 regulates the acquisition of gliogenic competence by neural stem cells. *Stem cell reports*, 5(3):365–377, 2015.

[TY06] Kazutoshi Takahashi and Shinya Yamanaka. Induction of pluripotent stem cells from mouse embryonic and adult fibroblast cultures by defined factors. *Cell*, 126(4):663–676, 2006.

[VAS94] Sabine Volland, Eberhard Amtmann, and Gerhard Sauer. TNF accelerates the S-phase of the cell cycle in tumor cells. *International journal of cancer*, 56(5):698–705, 1994.

[VGO+20] Pauli Virtanen, Ralf Gommers, Travis E Oliphant, Matt Haberland, Tyler Reddy, David Cournapeau, Evgeni Burovski, Pearu Peterson, Warren Weckesser, Jonathan Bright, et al. SciPy 1.0: fundamental algorithms for scientific computing in Python. *Nature methods*, 17(3):261–272, 2020.

[VTWP+25] Ramon Viñas Torné, Maciej Wiatrak, Zoe Piran, Shuyang Fan, Liangze Jiang, Sarah A Teichmann, Mor Nitzan, and Maria Brbić. Systema: a framework for evaluating genetic perturbation response prediction beyond systematic variation. *Nature Biotechnology*, pages 1–10, 2025.

[WG68] Martin B Wilk and Ram Gnanadesikan. Probability plotting methods for the analysis of data. *Biometrika*, 55(1):1–17, 1968.

[WKZ+21] Wei Wu, Jessica L Klockow, Michael Zhang, Famyrah Lafortune, Edwin Chang, Linchun Jin, Yang Wu, and Heike E Daldrup-Link. Glioblastoma multiforme (GBM): An overview of current therapies and mechanisms of resistance. *Pharmacological research*, 171:105780, 2021.

[ZL02] Wei Zhang and Hui Tu Liu. MAPK signal pathways in the regulation of cell proliferation in mammalian cells. *Cell research*, 12(1):9–18, 2002.

[ZPO17] Luke Zappia, Belinda Phipson, and Alicia Oshlack. Splatter: simulation of single-cell RNA sequencing data. *Genome biology*, 18(1):174, 2017.
